# Ubiquity of the Symbiont *Serratia symbiotica* in the Aphid Natural Environment: Distribution, Diversity and Evolution at a Multitrophic Level

**DOI:** 10.1101/2021.04.18.440331

**Authors:** Inès Pons, Nora Scieur, Linda Dhondt, Marie-Eve Renard, François Renoz, Thierry Hance

## Abstract

Bacterial symbioses are significant drivers of insect evolutionary ecology. However, despite recent findings that these associations can emerge from environmentally derived bacterial precursors, there is still little information on how these potential progenitors of insect symbionts circulates in the trophic systems. The aphid symbiont *Serratia symbiotica* represents a valuable model for deciphering evolutionary scenarios of bacterial acquisition by insects, as its diversity includes intracellular host-dependent strains as well as gut-associated strains that have retained some ability to live independently of their hosts and circulate in plant phloem sap. These strains represent a potential reservoir for the emergence of new and more intimate symbioses. Here, we conducted a field study to examine the distribution and diversity of *S. symbiotica* found in aphid populations, as well as in different compartments of their surrounding environment. A total of 250 aphid colonies, 203 associated insects, and 161 plant samples associated with aphid colonies were screened for *S. symbiotica*. Twenty percent of aphids were infected with *S. symbiotica*, and the symbiont includes a wide diversity of strains with varied tissue tropism corresponding to different lifestyle. We also showed that the prevalence of *S. symbiotica* is influenced by seasonal temperatures. For the first time, we found that *S. symbiotica* was present in non aphid species and in host plants, and that the prevalence of the bacterium in these samples was higher when associated aphid colonies were infected. Furthermore, phylogenetic analyses suggest the existence of horizontal transfers between the different trophic levels examined. These results provide a completely new picture of the ubiquity of an insect symbiont in nature. They suggest that ecological interactions promote the dissemination of strains that are still free-living and poorly specialized, and for which plants are a proabable reservoir for the acquisition of new bacterial partners in insects.

## INTRODUCTION

Numerous studies have revealed the diversity of microbial symbiotic associations in insects and their role in adapting to particular lifestyles or diets deficient in certain elements (Moran 2007; Douglas 2011; McFall-Ngai et al. 2013). Many insect species that have specialized on nutritionally unbalanced foods depend on obligate symbionts to synthesize deficient nutrients in their diets (Douglas 1998; Moran and Baumann 2000; Baumann et al. 2013). Obligate nutritional symbioses represent key evolutionary innovations, allowing insects to diversify in ecological niches otherwise inaccessible (Wernegreen 2017). Insects can also harbor facultative symbionts that are often involved in more recent associations and affect a wide range of host life history traits, even if they are not essential for host survival (Oliver et al. 2010; Feldhaar 2011; Heyworth and Ferrari 2015). Depending on the ecological context, facultative symbionts may act as mutualistic partners providing benefits to hosts, but may also produce fitness costs (Russell and Moran 2006; Vorburger and Gouskov 2011; Oliver et al. 2014; Zytynska et al. 2021). They suggest that their persistence is determined by a balance of costs and benefits. In recent years, the origin of bacterial mutualism in insects has been well studied (Husník et al. 2011; Clayton et al. 2012; Sudakaran et al. 2017; Manzano-Marín et al. 2020), particularly in stink bugs that pick up environmental bacteria that become mutualistic partners in specialized cavities of their digestive tract (Kikuchi et al. 2007; Takeshita and Kikuchi 2017). However, there is still little data regarding how potential progenitors of insect symbionts circulate in the environment and to what extent these circulating bacteria can spread to different trophic levels.

Several successive evolutionary steps can be considered in the establishment of mutualistic symbioses between insects and bacteria: 1) an infection by a generalist bacterium that circulates in the trophic chain from the environment, 2) if this infection results in a loss of fitness, a reduction of virulence by selection of the most resistant hosts and the least virulent bacteria in successive coevolutionary processes, 3) a contribution of benefits in particular circumstances, benefits greater than fitness costs for the host, and which allow the establishment of a profitable association for both partners under specific conditions, 4) the perpetuation of the association by a permanent reacquisition via a circulation of the symbiont in the trophic chain, 5) the establishment of horizontal and/or vertical transfers, and 6) the development of an increasingly tight relationships between the partners with a reduction in the size of the bacterial genome (Hosokawa et al. 2016; Latorre and Manzano-Marín 2017; Takeshita and Kikuchi 2017; Gil and Latorre 2019). Depending on the degree of evolution of these systems, each of these stages should still be observed or in many cases, the intermediate stages have disappeared over time, making the reconstruction of this evolutionary history more complex. In this evolutionary scheme, transmission mechanisms are known to be instrumental in the establishment and evolution of symbioses in natural host populations (Bright and Bulgheresi 2010). Although vertical transmission is the primary way by which symbioses are stably maintained (Salem et al. 2015), symbionts can also undergo occasional horizontal transfers or be acquired directly from the environment (Kikuchi et al. 2007; Gehrer and Vorburger 2012; Caspi-Fluger Ayelet et al. 2012; Łukasik et al. 2015; Hosokawa et al. 2016). Novel associations could emerge through these latter mechanisms, subsequently allowing the rapid acquisition of new ecological traits by insects and the expansion of host ranges for symbiotic bacteria (Su et al. 2013; Oliver et al. 2014).

In nature, the occurrence of bacterial strains capable of living freely outside of insects but related to insect endosymbionts suggests their potential role as source of insect symbionts. For example, a strain of the bacterium *Sodalis* isolated from a hand wound in a human host but originated from a dead tree branch was found to share a very close relationship with *Sodalis glossinidius* and other menbers of Sodalis-allied clade of insect symbionts (Clayton et al. 2012; Chari et al. 2015). *Arsenophonus* bacteria are known to infest plants while members of the genus are widely distributed in insect populations (Bressan et al. 2009; Jousselin et al. 2013). *Erwinia* bacteria, generally described as phytopathogens, are also found associated with insects, such as aphids where it was first described as a gut associate (Harada and Ishikawa 1993; Harada et al. 1997; Nadarasah and Stavrinides 2011). More recently, some *Cinara* aphids have been found to harbor *Erwinia*-related obligate symbionts that complement the ancestral obligate symbiont *Buchnera aphidicola* (Meseguer et al. 2017). Genomic analyses indicate that these associations originate from the acquisition of free-living *Erwinia* strains, likely acquired horizontally throught plants, and have evolved into an intracellular lifestyle (Manzano-Marín et al. 2020). Taken together, all these findings support the hypothesis of the existence of an environmental pool of bacteria from which new intimate partnerships with insects can emerge. In addition to this, a growing number of studies have recently shown that symbiotic associations between insects and bacteria evolve in a very dynamic fashion, involving the acquisition of new symbionts, and/or the loss and replacement of established bacterial partners, even in the context of obligate associations established sometimes for millions of years (Koga and Moran 2014; Hosokawa et al. 2016; Husnik and McCutcheon 2016; Manzano-Marín et al. 2017; Sudakaran et al. 2017; Chong and Moran 2018; Matsuura et al. 2018; Mao and Bennett 2020). For example, genomic analyses in aphids indicate that the dependence of some species on co-obligate symbiotic bacteria has arisen independently many times during their evolutionary history (Manzano-Marín et al. 2016, 2017; Meseguer et al. 2017; Manzano-Marín et al. 2020; Monnin et al. 2020). The repeated replacement of pre-existing symbionts by other microbial partners is now considered a redundant evolutionary process that occurs in many insect species, suggesting 1) the continuous formation of new mutualistic associations in nature, and 2) the existence of a pool of environmental symbionts from which new intimate, facultative, or obligate associations are formed in nature.

*Serratia symbiotica*, one of the most common symbionts in aphids (Oliver et al. 2010; Henry et al. 2015; Zytynska and Weisser 2016; Monnin et al. 2020), is a valuable candidate to decipher the origin and the evolution of bacterial mutualism in insects. It includes a great diversity of strains associated with very distinct biological features and reflecting the various associations that bacteria can share with insects (Burke and Moran 2011; Manzano-Marín and Latorre 2016; Pons et al. 2019a; Monnin et al. 2020; Perreau et al. 2020). The strains studied in aphids of the subfamily *Aphidinae* have been first described as intracellulaire facultative partners because they can invade host cells (including bacteriocytes and sheath cells), and can be associated with protective phenotypes (against parasitoids and high temperatures) (Oliver et al. 2003; Burke et al. 2009; Heyworth and Ferrari 2015). *S. symbiotica* strains associated with aphids of the subfamilies Lachninae and Chaitophorinae are intracellular symbionts involved in co-obligate associations, compensating some metabolic capacities lost by the ancient obligate symbiont *B. aphidicola* (Lamelas et al. 2011; Manzano-Marín and Latorre 2014, 2016; Manzano-Marín et al. 2018; Monnin et al. 2020). Moreover, the species *S. symbiotica* also includes strains that are gut-associates (Renoz et al. 2018; Pons et al. 2019a; Perreau et al. 2020) and which can be cultivated freely on a pure artificial medium (Sabri et al. 2011; Grigorescu et al. 2017). These strains, slightly pathogenic to aphids, are considered to be at the pathogen-symbiont interface (Pons et al. 2019a; Perreau et al. 2020). We recently showed that these gut-associated strains that retain some free-living capacities, are extracellularly transmitted via contamination with honeydew (Pons et al. 2019a) and/or through the plant phloem (Pons et al. 2019b). They can be horizontally transferred between aphids through host plant, and their uptake by plant roots can induce new bacterial infections of aphids, as well as positive fitness effects on the host plant (Pons et al. 2019b). A field study also suggest that *S. symbiotica* may reside in the gut of ants tending aphids (Renoz et al. 2018). Taken together, these results indicates that certain *S. symbiotica* strains have a tremendous ability to circulate from one compartment to another, with an aptitude to perform horizontal transfers between both phylogenetically close and distant species in the environment. We suggest that these strains may thrive in the environment where aphids prosper and could provide an environmental source for the establishment of new symbiotic asociations between aphids and bacteria.

While *S. symbiotica* strains with a broad spectrum of infection may represent ideal candidates to refine our understanding of evolutionary scenarios of symbiont acquisition, we still do not know how *S. symbiotica* strains are distributed in nature, how they circulate and how they spread in aphid populations. To address these issues, we conducted a field study to provide comprehensive picture of the ubiquity of *S. symbiotica* across the food web. We investigated the distribution, diversity and evolution of *S. symbiotica* infections at a multitrophic level, including 58 aphid species, aphid-interacting insects (i.e. predators, parasitoids, etc.), and host plants. This study allowed us 1) to examine the prevalence of *S. symbiotica* in aphids and their environment, and to detect environmental and ecological factors that influence its prevalence, 2) to determine the diversity of tissue tropism exhibited by *S. symbiotica* in natural insect populations, and 3) to investigate the propensity of *S. symbiotica* to circulate and be transferred horizontally in the natural environment of aphids. Using diagnostic PCR, fluorescence *in situ* hybridization and phylogenetic approaches, we provide a picture of the diversity and the ubiquity of *S. symbiotica* in a natural environment. Our study highlights the presence of *S. symbiotica* strains at different levels of food webs and suggests the existence of ecological compartments for the exchange and acquisition of *S. symbiotica* in insects, serving as potential interfaces for the emergence of new symbiotic associations.

## MATERIALS AND METHODS

### Sample collection

To get a compregensive picture of the ubiquity of *S. symbiotica* in the natural aphid environment, we sampled 1) aphid colonies of different species, 2) insects potentially in interaction with aphids in the sampled colonies, and 3) the host plant. Sampling was carried out to maximize the diversity of aphid subfamilies and genera, representing 3 subfamilies of Aphididae, 27 genera and 58 species. Field specimens were sampled between May and August 2018 on various host plants at several locations in Belgium (Walloon Brabant province) (Table S1). The sampling also includes some aphids sampled in Italy, France, Germany and Rwanda (2017). A total of 614 samples were collected, containing 250 aphid colonies, 203 insects associated with aphid colonies (tending ants, hoverfly larvae, larvae and adults of ladybugs, parasitoids, bugs, aphid midge larvae, moth larvae, fly larvae and lacewing larvae), and 161 host plant samples (Table S1). To minimize the risk of pseudo-replicating, *S. symbiotica* infection was verified in only one pool of aphids per host plant. When insects were observed within aphid colonies, individuals were systematically collected and preserved in 90% ethanol at room temperature until use. Plant samples (stem and leaf pieces) associated with the sampled aphid colonies were also systematically collected and stocked at −80°C. For samples collected in Belgium, the mean daily temperatures and the maximum daily temperatures during the different months of sampling in Belgium were obtained via the database of IRM (Institut Royal Météorologique) of Belgium and the meteobelgique.be website (Table S1).

**Table 1.** Summary of the natural occurrence of *S. symbiotica* within 250 aphid colonies, across the different aphid genera. Aphids positive to specific primers correspond to aphids positive to the three primers designed to detect *S. symbiotica* strains with a potential gut localization.

| Aphid genus |  | Number of colonies | Positive to *S. symbiotica* |
| --- | --- | --- | --- |
| **Aphidinae** |  |  |  |
| Macrosophini |  |  |  |
|  | *Acyrthosiphon* | 3 | 0 |
|  | *Brachycaudus* | 3 | 1 |
|  | *Capitophorus* | 1 | 1 |
|  | *Cavariella* | 5 | 2 |
|  | *Corylobium* | 1 | 0 |
|  | *Dysaphis* | 1 | 0 |
|  | *Hyadaphis* | 1 | 0 |
|  | *Hyalopteroides* | 4 | 1 |
|  | *Hyperomyzus* | 10 | 0 |
|  | *Macrosiphoniella* | 8 | 1 |
|  | *Macrosiphum* | 5 | 4 |
|  | *Megoura* | 1 | 0 |
|  | *Metopeurum* | 2 | 0 |
|  | *Metopolophium* | 4 | 0 |
|  | *Myzus* | 1 | 0 |
|  | *Sitobion* | 8 | 1 |
|  | *Straticobium* | 1 | 0 |
|  | *Uroleucon* | 49 | 0 |
| Aphidini |  |  |  |
|  | *Aphis* | 121 | 35 |
|  | *Hyalopterus* | 3 | 0 |
|  | *Rhopalosiphum* | 1 | 0 |
|  | *Schizaphis* | 3 | 2 |
|  | *Toxoptera* | 6 | 2 |
| **Chaitophorinae** |  |  |  |
| Chaitophorini |  |  |  |
|  | *Chaitophorus* | 1 | 0 |
|  | *Periphyllus* | 1 | 1 |
| **Calaphidinae** |  |  |  |
| Panaphidini |  |  |  |
|  | *Pterocallis* | 1 | 0 |
|  | *Tinocallis* | 1 | 0 |
|  | Not identified | 4 | 0 |
| **Total** |  | **250** | **51** |

### DNA extraction

Insect DNA extraction was performed using a high salt-extraction method (Aljanabi and Martinez 1997). Extraction was carried out on a pool of two to six aphids from each colony and on a pool of two to three individuals for associated tending ants and larvae (Jousselin et al. 2013; Renoz et al. 2018). Pools were carried out to avoid the risk of missing infection, when *S. symbiotica* is present. For the other associated insects, extraction was performed on a single individual. Plant DNA was extracted using the CTAB method (Doyle 1991). Prior to extraction, plant samples were surface-sterilized with 99% ethanol, 10% bleach, and rinsed with sterile water.

### Insect identification

For insect species identification, the primers LepF and LepR (presented in Table S2) were used to amplify the target 658-bp fragment of cytochrome c oxidase subunit I (COI) gene (D’acier et al. 2014). PCR reactions were conducted in a final volume of 15 μl containing 1μl of the template DNA lysate, 0.5 μM of each primer, 200 μM dNTPs, 1 × buffer and 0.625 unit of Taq DNA polymerase (Roche).

**Table 2.** Summary of the natural occurrence of *S. symbiotica* within 161 host plants. Plants positive to specific primers correspond to plants positive to the three primers designed to detect *S. symbiotica* strains with a potential localization in aphid gut and free-living capacity. The number of plants associated with infected aphids, as well as the number of plants infected with *S. symbiotica* associated with infected aphids were considered.

| Plant genus | Number of samples | Positive to *S. symbiotica* | Plant associated with infected aphids |  | Infected plant associated with infected aphids |  |
| --- | --- | --- | --- | --- | --- | --- |
|  |  |  | Number | Infected Aphid genus | Number | Infected Aphid genus |
| *Achillea* | 1 | 0 | 1 | *Macrosiphoniella* | 0 | - |
| *Artemisia* | 5 | 0 | 0 | - | 0 | - |
| *Campanula* | 1 | 0 | 0 | - | 0 | - |
| *Centaurea* | 2 | 0 | 0 | - | 0 | - |
| *Centranthus* | 1 | 0 | 0 | - | 0 | - |
| *Chaenomeles* | 1 | 0 | 0 | - | 0 | - |
| *Chenopodium* | 2 | 0 | 0 | - | 0 | - |
| *Cirsium* | 26 | 4 | 7 | *Aphis*,<br>*Capitophorus*,<br>*Macrosiphum* | 3 | *Aphis* |
| *Clematis* | 1 | 0 | 1 | *Aphis* | 0 | - |
| *Conyza* | 1 | 0 | 0 | - | 0 | - |
| *Corylus* | 2 | 0 | 0 | - | 0 | - |
| *Crataegus* | 1 | 0 | 0 | - | 0 | - |
| *Crepis* | 1 | 0 | 0 | - | 0 | - |
| *Daucus* | 8 | 0 | 2 | *Aphis* | 0 | - |
| *Epilobium* | 10 | 0 | 2 | *Aphis* | 0 | - |
| *Epipactis* | 2 | 0 | 0 | - | 0 | - |
| *Eupatorium* | 2 | 0 | 1 | *Aphis* | 0 | - |
| *Gaillet* | 1 | 0 | 0 | - | 0 | - |
| *Glebionis* | 1 | 0 | 0 | - | 0 | - |
| *Hedera* | 1 | 0 | 1 | *Aphis* | 0 | - |
| *Heracleum* | 7 | 1 | 2 | *Aphis*, *Cavariella* | 0 | - |
| *Hibiscus* | 1 | 0 | 0 | - | 0 | - |
| *Jacobaea* | 6 | 1 | 2 | *Aphis* | 0 | - |
| *Lactuca* | 2 | 0 | 0 | - | 0 | - |
| *Laphangium* | 2 | 0 | 2 | *Aphis*,<br>*Brachycaudus* | 0 | - |
| *Leucanthemum* | 3 | 0 | 2 | *Aphis* | 0 | - |
| *Lythrum* | 1 | 0 | 0 | - | 0 | - |
| *Malus* | 1 | 1 | 0 | - | 0 | - |
| *Oenothera* | 1 | 0 | 0 | - | 0 | - |
| *Phragmites* | 3 | 0 | 0 | - | 0 | - |
| *Populus* | 1 | 0 | 0 | - | 0 | - |
| *Ranunculus* | 1 | 1 | 0 | - | 0 | - |
| *Robinia* | 2 | 0 | 0 | - | 0 | - |
| *Rosa* | 3 | 0 | 2 | *Macrosiphum* | 0 | - |
| *Rumex* | 8 | 2 | 2 | *Aphis* | 1 | *Aphis* |
| *Salvia* | 1 | 0 | 0 | - | 0 | - |
| *Sambucus* | 1 | 1 | 1 | *Aphis* | 1 | *Aphis* |
| *Solanum* | 2 | 1 | 1 | *Aphis* | 1 | *Aphis* |
| *Sonchus* | 32 | 1 | 0 | - | 0 | - |
| *Spiraea* | 2 | 0 | 0 | - | 0 | - |
| *Symphytum* | 1 | 0 | 0 | - | 0 | - |
| *Tanacetum* | 6 | 1 | 0 | - | 0 | - |
| *Verbena* | 2 | 0 | 0 | - | 0 | - |
| *Vesca* | 1 | 0 | 0 | - | 0 | - |
| *Vicia* | 1 | 0 | 0 | - | 0 | - |
| **Total** | **161** | **14** | **37** | **-** | **6** | **-** |

The thermocycling profile consisted of 94°C for 1 min; 6 cycles of 94 °C for 1 min, 45 °C for 1 min and 30 s, and 72 °C for 1 min and 15 s; followed by 36 cycles of 94°C for 1 min, 51°C for 1 min and 30 s, and 72 °C for 1 min and 15 s; with a final 5 min extension period of 72°C. Amplicons were then purified before sequencing (Macrogen Inc., Amsterdam). The resulting sequences were cleaned and aligned using Geneious® v9.1.5 (Kearse et al. 2012) and insects were identified by comparing resulting COI sequence data to the GeneBank nucleotide database using BLAST.

### Diagnostic screening for *S. symbiotica*

All samples were tested for the presence of *S. symbiotica* using PCR assays based on the 16S rRNA gene using the specific primers 16SA1 and PASScmp (presented in Table S2). PCR amplification was performed as previously described (Pons et al. 2019a). PCR reactions were conducted in a final volume of 15 μl containing 1μl of the template DNA lysate, 0.5 μM of each primer, 200 μM dNTPs, 1X buffer and 0.625 unit of Taq DNA polymerase (Roche). The thermocycling profile consisted of 95°C for 5min; 35 cycles at 95 °C for 30 sec, 55 °C for 1 min 30 s and 72 °C for 1 min 30s; 72°C for 7 min. DNA from an infected line of the pea aphid *A. pisum* was used as a positive control (Burke et al. 2009) and in addition to the negative control, DNA from an uninfected line of the black bean aphid *A. fabae* was used as a negative control (Vorburger and Gouskov 2011). Amplicons were purified and sequenced in both directions (Macorgens Inc., Amsterdam). Sequence alignments were done using Geneious® v9.1.5 (Kearse et al. 2012) and compared to sequences on GenBank using BLAST. The sequences were deposited in GenBank.

### Phylogenetic analyses

The diversity of *S. symbiotica* strains was also characterized using the partial sequence of four housekeeping genes *accD*, *gyrB*, *murE* and *recJ* (Table S2) (Henry et al. 2013; Łukasik et al. 2015). DNA samples that were found positive for *S. symbiotica* were subjected to PCR amplification under the following conditions: initial denaturation at 94°C for 5 min; 35 cycles of denaturation at 94°C for 30 s, annealing at 60-64°C depending on primers for 30s, extension at 72°C for 1 min; a final extension at 72°C for 5 min (Renoz et al. 2018). Amplicons were then purified and sequenced in both directions (Macrogen Inc., Amsterdam). Sequences obtained were cleaned and aligned using Geneious® v9.1.5 (Kearse et al. 2012).

Phylogenetic associations were analyzed for 54 *S. symbiotica* strains obtained in this work, sequences from the three cultivable *S. symbiotica* strains that were isolated in our laboratory (Sabri et al. 2011; Foray et al. 2014; Grigorescu et al. 2017; Renoz et al. 2020), eight sequences from *S. symbiotica* that were found in a recent field study (from aphids collected in Belgium in summer 2015) (Renoz et al. 2018) and twelve sequences from *S. symbiotica* whose genomes have been sequenced and available in Genbank (Burke and Moran 2011; Lamelas et al. 2011; Manzano-Marín and Latorre 2014; Manzano-Marín et al. 2016; Meseguer et al. 2017; Manzano-Marín et al. 2018; Nikoh et al. 2019; Monnin et al. 2020). *Serratia proteamaculans* 568 was used as outgroup. All sequences are available in GenBank (Table S3). SeaView v4.6.1 was used to align sequences, and all ambiguously aligned regions identified by GBlocks v0.91b (Castresana 2000) were eliminated, as were regions of incomplete data at the 3’ and 5’ ends of the targeted regions. PartitionFinder v1.1.0 (Lanfear et al. 2012) was used to determine the substitution model for each gene. Phylogenetic analysis was performed by Bayesian inference methods with MrBayes v3.2.7a (Ronquist et al. 2012). Indels were treated as missing data. Four Markov chain Monte Carlo (MCMC) simulations were run independently for 10,000,000 generations. Trees and model parameters were sampled every 10,000 generations. Convergence of the MCMCs was estimated in three ways: (1) the standard deviation of split frequencies was < 0.01, (2) visual inspection of the plot of the log-likelihood score at each sampling point suggested that the four chains reached stationarity, and (3) the posterior probability plots of all splits for paired MCMC runs showed high correlation, which diagnosed convergence among the four chains (Nylander et al. 2008). The trees of the burn-in for each run were excluded from the tree set, and the remaining trees from each run were combined to form the full sample of trees assumed to be representative of the posterior probability distribution.

**Table 3.** Summary of the natural occurrence of *S. symbiotica* within 203 insects associated with aphid colonies. Insects positive to specific primers correspond to insects positive to the three primers designed to detect *S. symbiotica* strains with a potential localization in aphid gut and free-living capacity. The number of insects infected with *S. symbiotica* associated with infected aphids was considered.

| Insect genus |  | Number of samples | Positive to *S. symbiotica* | Infected insect associated with infected aphids |  |
| --- | --- | --- | --- | --- | --- |
|  |  |  |  | Number | Infected Aphid genus |
| Ants |  |  |  |  |  |
|  | *Camponotus* | 3 | 0 | - | - |
|  | *Formica* | 1 | 0 | - | - |
|  | *Lasius* | 78 | 7 | 6 | Capitophorus, Aphis |
|  | *Linepithema* | 2 | 2 | 2 | *Aphis* |
|  | *Myrmica* | 7 | 1 | 0 | - |
|  | Not identified | 7 | 0 | - | - |
| Aphid Midge Larvae |  |  |  |  |  |
|  | *Aphidoletes* | 7 | 2 | 1 | *Aphis* |
| Bugs |  |  |  |  |  |
|  | *Coreus* | 2 | 1 | 1 | *Aphis* |
|  | *Deraeocoris* | 1 | 0 | - | - |
|  | *Dictyla* | 2 | 1 | 0 | - |
|  | *Dicyphus* | 1 | 0 | - | - |
|  | *Dolycoris* | 3 | 0 | - | - |
|  | *Graphosoma* | 1 | 0 | - | - |
|  | *Malacocoris* | 1 | 0 | - | - |
|  | *Orius* | 1 | 0 | - | - |
|  | *Orthops* | 1 | 0 | - | - |
|  | *Pinalitus* | 1 | 0 | - | - |
|  | Not identified | 1 | 0 | - | - |
| Fly Larvae |  |  |  |  |  |
|  | *Chamaemyiidae* | 1 | 1 | 1 | *Aphis* |
| Hoverfly Larvae |  |  |  |  |  |
|  | *Episyrphus* | 6 | 1 | 1 | *Aphis* |
|  | *Eupeodes* | 3 | 0 | - | - |
|  | *Paragus* | 7 | 0 | - | - |
|  | *Platycheirus* | 2 | 0 | - | - |
|  | *Scaeva* | 11 | 3 | 1 | *Capitophorus* |
|  | *Syrphus* | 5 | 0 | - | - |
| Lacewing Larvae |  |  |  |  |  |
|  | *Chrysoperla* | 1 | 0 | - | - |
| **Ladybugs** |  |  |  |  |  |
|  | *Coccinella* | 8 | 1 | 0 | - |
|  | *Harmonia* | 7 | 1 | 1 | *Aphis* |
|  | *Hippodamia* | 2 | 0 | - | - |
|  | *Propylea* | 2 | 0 | - | - |
|  | *Rhyzobius* | 1 | 0 | - | - |
|  | *Scymnus* | 1 | 0 | - | - |
|  | Not identified | 1 | 0 | - | - |
| **Moth Larvae** |  |  |  |  |  |
|  | *Eupithecia* | 3 | 1 | 1 | *Aphis* |
|  | *Hecatera* | 1 | 0 | - | - |
|  | *Mompha* | 1 | 0 | - | - |
|  | *Pieris* | 1 | 0 | - | - |
| **Parasitoids** |  |  |  |  |  |
|  | *Aphelinus* | 1 | 0 | - | - |
|  | *Aphidius* | 2 | 1 | 0 | - |
|  | *Binodoxys* | 2 | 0 | - | - |
|  | *Diglyphus* | 1 | 0 | - | - |
|  | *Eulophinae* | 1 | 0 | - | - |
|  | *Lysiphlebus* | 7 | 2 | 0 | - |
|  | *Pachyneuron* | 1 | 0 | - | - |
|  | *Pnigalio* | 1 | 0 | - | - |
|  | *Tetrastichinae* | 3 | 0 | - | - |
| **Total** |  | **203** | **25** | **15** | - |

### Localization of *S. symbiotica*

To determine the tissue tropism of *S. symbiotica* strains in aphids and in the other insects sampled in the colonies, whole-mount fluorescence *in situ* hybridization (FISH) was carried out as described in (Koga et al. 2009; Renoz et al. 2018; Pons et al. 2019a). All insect samples positive to *S. symbiotica* were tested. The following oligonucleotide probes were used: Cy5-ApisP2a (5’-Cy5-CCTCTTTTGGGTAGATCC-3’) targeting 16S rRNA of *B. aphidicola* and Cy3-PASSisR (5’-Cy3-CCCGACTTTATCGCTGGC-3’) targeting 16S rRNA of *S. symbiotica*. Insect tissues were stained with SYTOX Green. Samples (between 1 to 6 per colony) were whole mounted and observed under a Zeiss LSM 710 confocal microscope. Negative controls consisted of aphids not infected with *S. symbiotica* and stained with the two probes or infected aphids with no-probe staining. Positive controls comprised of artificially and naturally infected aphids.

## Statistical analyses

We analyzed whether the proportion of aphids harboring *S. symbiotica* differs according to their host plant range, as carry out in (Henry et al. 2015): specialized (feeding on a single plant species or group of closely related species, N=13), restricted (feeding on a single plant family, N=91), polyphagous (feeding on various plant families, N=105), or obligatory host-plant alternating (N=31). The presence of *S. symbiotica* in aphids was analyzed using generalized linear models (GLM) with a binomial error structure and a logit-link function, and the degree of plant specialization by aphids was the explanatory variable. We also tested if the proportion of aphids infected by *S. symbiotica* differs in populations collected from different plant species, using Pearson’s Chi-square statistic. Aphids on plant species from which we had less than five collection, and where *S. symbiotica* symbiont infected < 2% of the individuals on a plant, were excluded from the analysis as recommended in (Henry et al. 2015). We also examined the effect of the temperature (mean daily and maximum daily) on the proportion of aphids harboring *S. symbiotica*. We tested if *S. symbiotica* infection was influenced by high seasonal temperatures, because this symbiont is known to protect its host from heat shock, especially aphids collected from arid regions (Russell and Moran 2006; Henry et al. 2013). The presence of *S. symbiotica* in aphids was analyzed on pooled data using generalized linear models (GLM) with a binomial error structure and a logit-link function, and the temperature was the explanatory variable. The relationship between mean daily temperatures or maximum daily temperatures and the proportion of aphids harboring *S. symbiotica* was tested on pooled data using non-parametric Spearman’s correlation coefficients. We also analyzed if the proportion of tending ants, associated other insects and host plants harboring *S. symbiotica* were correlated with the infection status of aphid colonies. The presence of *S. symbiotica* in these different samples was analyzed using generalized linear models (GLM) with a binomial error structure and a logit-link function. The infection status of the aphid colonies was the explanatory variable and for the associated insects, the insect feeding specialization (parasitoids, phytophagous or predators) was also one explanatory variable. Statistical analyses were performed using the software R version 3.5.3 (R Development Core team, 2019), using the *GrapheR* package.

## RESULTS

### Distribution of *S. symbiotica* in natural aphid population

The presence of *S. symbiotica* was examined in 250 aphid colonies from 58 species of the Aphididae family (Table S1). These species were mostly part of the Macrosiphini (31 species) and Aphidini (23 species) tribes belonging to the Aphidinae subfamily (Table S1). Other species were members of the subfamilies Chaitophorinae (2 species) and Calaphidinae (2 species) (Table S1). The tribe Aphidini was the most collected group with 121 colonies belonging to 19 species of the genus *Aphis* (Table 1). Among the 250 colonies, 51 displayed a positive infection to *S. symbiotica* (20%), belonging to 22 aphid species (38%) (Table 1 and Table S1). Within each infected aphid genus, the infection prevalence ranged from 13 to 100% (Table 1). The infection rate of *S. symbiotica* in *Aphis*, which is the most represented group in our sampling with 121 collected colonies, reached 29% (35/121) and 69% of infected aphid colonies belonged to the genus *Aphis* (35/51) (Table 1). More precisely, 27% (22/82) of *Aphis fabae* colonies, which is the most represented species of the genus *Aphis*, were found to be infected (Table S1). We also found a hight infection rate of *S. symbiotica* in the genera *Macrosiphum* (4/5), *Capitophorus* (1/1) and *Periphyllus* (1/1) (Table 1). In contrast, we observed that in the genera *Uroleucon* and *Hyperomyzus*, the infection rate was zero (0/49 and 0/10, respectively, Table 1).

### Environmental and ecological factors influencing *S. symbiotica* prevalence

We showed that among the 45 host plant genera collected (all containing aphids), 15 were associated with infected aphids (33%, Table 2). Of the 37 plants associated with infected aphids (all genera included), 7 were members of the genus *Cirsium* (18%) (Table 2). We asked whether the degree of plant specialization by aphids was associated with *S. symbiotica* infection in aphids. We found no significant differences in the proportion of aphids harboring *S. symbiotica* according to their degree of plant specialization (GLM, df = 3, χ^2^ = 6.35, p = 0.096, Figure S1). We also analyzed whether the infection of aphids with *S. symbiotica* that feed on multiple host plants was correlated with the species of plant on which they were collected. Sufficient data for analysis were available only for *A. fabae* that feed on different host plants (*Circium arvense*, *Circium vulgare*, *Daucus carota*, *Heraclum sphondylium*, *Rumex obtusifolium*, and *Sonchus asper*). We found that there were no significant differences in the proportion of *A. fabae* harboring *S. symbiotica* collected from different plants (N=49, df = 5, χ2 = 3.59, p = 0.61). The distribution of *S. symbiotica* in this sampling is uniform across collections of *A. fabae* feeding on different plant species.

During the sampling in Belgium from May through August 2018, the mean daily temperatures ranged from 12.9 °C to 25.8 °C and that maximum daily temperatures ranged from 15.9 °C to 32.9 °C (Table S1). To test the impact of temperature on *S. symbiotica* prevalence, we focused on the data sampled in July and August when most samples were collected (Table S1). During these months, the mean daily temperatures ranged from 19.5 °C to 25.8 °C and the maximum daily temperatures ranged from 22.9 °C to 32.9 °C (Table S1). To perform analyzes, temperature data were pooled into 6 balanced categories each (Table S1). We showed that *S. symbiotica* frequencies varied across Belgium aphid populations exposed to varying summer temperatures (Figure 1). We found that the proportion of aphids harboring *S. symbiotica* was significantly different between mean daily temperatures (GLM, df = 1, χ^2^ = 2.53, p = 0.011) and maximum daily temperatures in July and August (GLM, df = 1, χ^2^ = 2.08, p = 0.038) (Figure 1A-B). We also showed a significant positive correlation between the proportion of aphids harboring *S. symbiotica* and the maximum daily temperatures (Spearman’s rs = 1, p = 0.017, Figure 1B). *S. symbiotica* frequency in aphid populations was thus significantly higher when temperatures were higher at time of collection, ranging from 7% at 23 °C to 31% at 30 °C.

**Figure 1.**
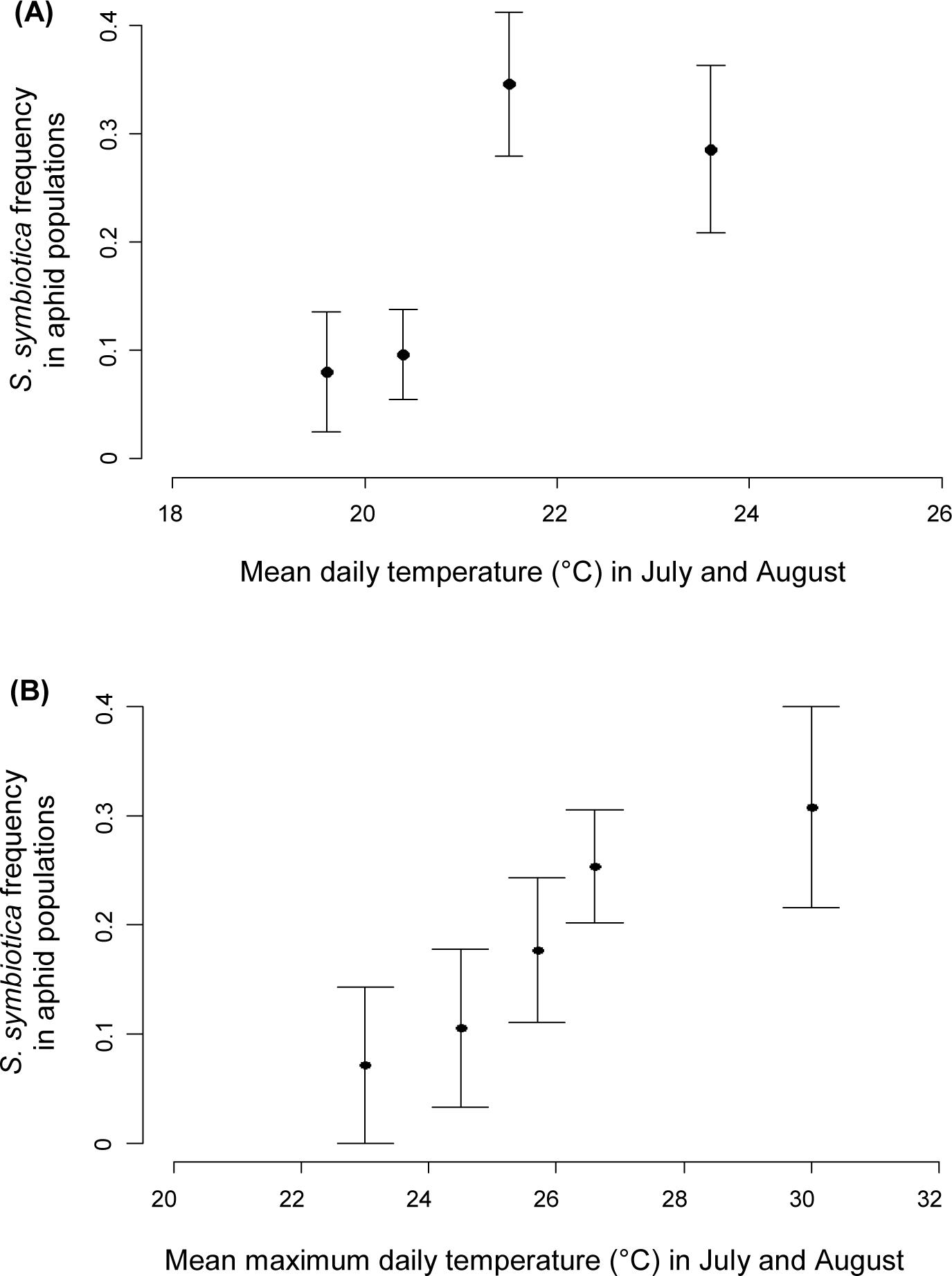
Relationships between *S. symbiotica* frequency across Belgian aphid populations and the mean daily temperature (**A**) or the maximum daily temperature (**B**) during sampling.

### Distribution of *S. symbiotica* in insect populations associated with aphid colonies

Other insects associated with aphid colonies were also collected to examine the presence of similar *S. symbiotica* strains in the trophic systems. Among 203 insects composed of 52 species, 25 displayed a positive infection to *S. symbiotica* (12%) (Table 3).

Ants were the most frequent insects found in aphid colonies and were represented by 10 different species (Table S1). Among the 98 tending ant samples, 10 exhibited a positive infection with *S. symbiotica*, representing an infection rate of about 10% (Table 3). These infected ants belonged to *Lasius niger, Linepithema humile* and *Myrmica rubra* species, with *L. niger* being the most represented species (7/10) (Table S1). We found that the proportion of tending ants harboring *S. symbiotica* was significantly higher when collected in infected aphid colonies (N = 8/27) compared to uninfected colonies (N = 2/71) (GLM, df = 1, χ^2^ = 13.55, p < 0.001, Figure 2A).

**Figure 2.**
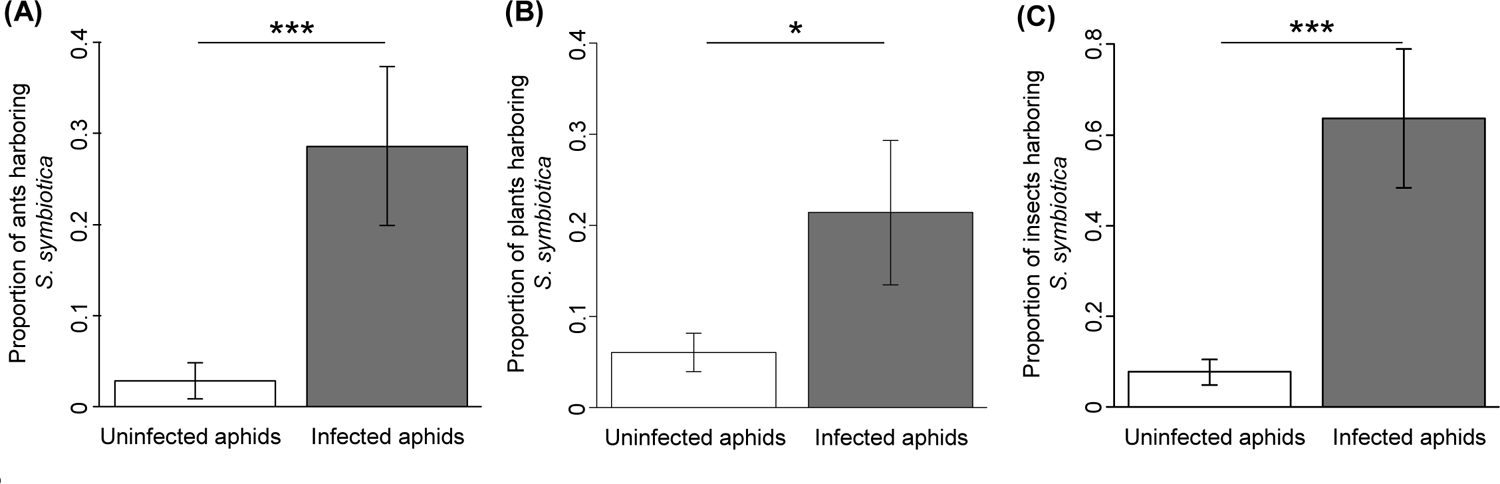
Prevalence of *S. symbiotica* in tending ants (A), plant samples (B) and associated insects (C) differ depending on the infection status of the aphid colonies. Columns represent the proportion of samples infected with *S. symbiotica* according to uninfected aphid colonies (white) or infected aphid colonies (dark grey). Error bars depict the standard error. Asterisks show significant differences (*: p<0.05, and ***: p<0.001).

Among the 15 bug individuals representing 10 species, two (*Dictyla humuli* and *Coreus marginatus*) were infected with *S. symbiotica* (Table S1). Twenty-two ladybugs were collected (larvae and adults), belonging to 6 species, and two adults were positive to *S. symbiotica* (*Harmonia axyridis* and *Coccinella septempunctata*; Table S1). Nineteen adult parasitoids were also found in aphid colonies, mainly collected when laying eggs into aphids (Table S1). Three individuals of three species (*Aphidius funebris, Lysiphlebus fabarum* and *Lysiphlebus hirticornis*) showed a positive infection (Table S1). These three infected individuals were not sampled from the infected aphid colonies. Of the 34 hoverfly larvae that were sampled, 4 were infected with *S. symbiotica* (*Episyrphus balteatus* and *Scaeva pyrastri*) (Table S1). Two aphid midge larvae (*Aphidoletes aphidimyza*) were infected out of 7, and one moth larva was infected out of 6 (*Eupithecia trisignaria*, Table S1). Moreover, one fly larva was collected and was positive for *S. symbiotica* (*Chamaemyiidae* sp.) whereas the only lacewing larva collected was not positive (Table 3 and Table S1). We found that the proportion of these associated insects harboring *S. symbiotica* was significantly higher when collected from infected aphid colonies (N = 7/11) compared to uninfected colonies (N = 7/90) (GLM, df = 1, χ^2^ = 19.86, p < 0.001, Figure 2C). We also examine if the insect feeding specialization (parasitoids, polyphagous or predators) can influence its symbiotic infection. We found that the feeding specialization did not significantly affect the proportion of these insects harboring *S. symbiotica* (GLM (*S. symbiotica*), df = 2, χ^2^ = 5.2, p = 0.074).

### Distribution of *S. symbiotica* in the natural aphid environment

To examine the presence of *S. symbiotica* in the surrounding environment of aphids, host plant associated with aphid colonies were collected. Among the 161 plant samples (11 leaves and 150 stems) collected from 52 plant species, 14 were positive for *S. symbiotica* (about 9%), belonging to 12 plant species (23%; Table S1). The genera *Sonchus* (N=32) and *Cirsium* (N=26) were the most collected groups (Table 2). The latter exhibited the highest infection rate with 29% of infection (4/14), followed by plants of the genus *Rumex* with 14% of infection (2/14, Table 2). We showed that among the 14 positive plant samples, only 6 were associated with positive aphid colonies (belonging to the genus *Aphis*) (Table 2). However, we found that the proportion of plants harboring *S. symbiotica* was significantly higher when associated with infected aphid colonies (N = 6/27) compared to uninfected colonies (N =8/132) (GLM, df = 1, χ^2^ = 5.8, p = 0.016, Figure 2B).

### Tissue tropism of *S. symbiotica* in insects

All insect samples that showed a positive infection to *S. symbiotica* were observed, using whole-mount fluorescence *in situ* hybridization. A total of 21 aphid colonies have been examined and observations revealed the existence of different patterns: 1) presence of *S. symbiotica* in the digestive tract, 2) in the sheath cells, and 3) in the secondary bacteriocytes (Figure 3).

**Figure 3.**
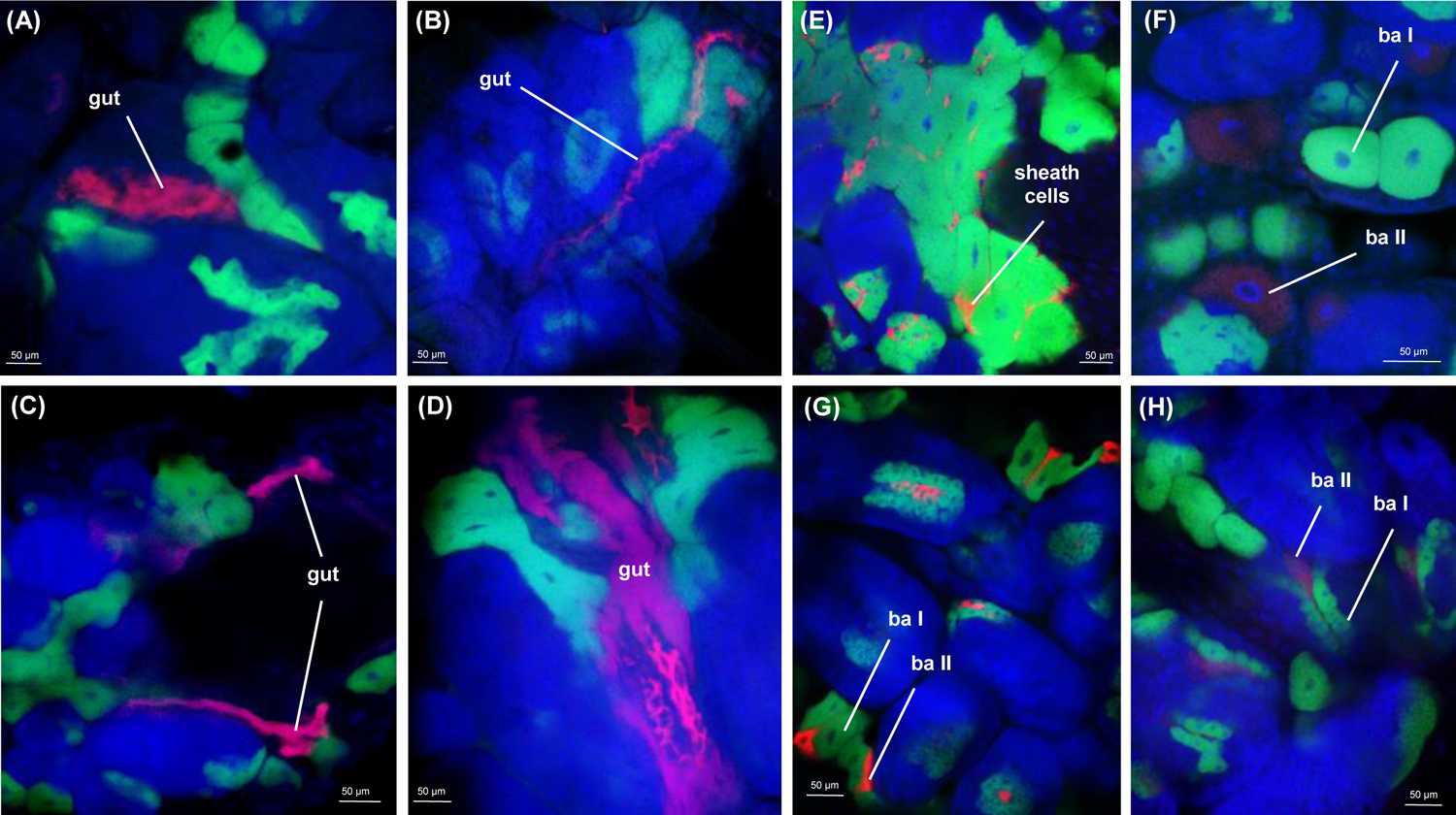
Whole-mount FISH of *S. symbiotica* in naturally infected aphids (ventral views). Red Cy3 signals are *S. symbiotica*, green Cy5 signals are *B. aphidicola*, and blue SYTOX Green signals are aphid tissues. **(A-D)** *S. symbiotica* found residing at the gut level of aphids. **(E)** *S. symbiotica* found residing at the sheath cells level of aphids. **(F-H)** *S. symbiotica* found residing at the bacteriocytes level of aphids (ba I: primary bacteriocyte and ba II: secondary bacteriocyte). **(A)** *Aphis grossulariae* (ID 491), **(B)** *Capitophorus elaeagni* (ID 605), **(C)** *Aphis fabae* (ID 517), **(D)** *Aphis pomi* (ID 511), **(E)** *Macrosiphum rosae* (ID 263), **(F)** *Macrosiphum mordvilkoi* (ID 135), **(G)** *Periphyllus testudinaceus* (ID 1), and **(H)** *Aphis fabae* (ID 380).

Gut-associated *S. symbiotica* were found either colonizing in the midgut or into the whole gut (Figure 3A-D). The digestive tract is localized in the middle of the aphid body between the primary bacteriocytes hosting the obligate symbiont *B. aphidicola* and is composed of multiple loops (Pons et al. 2019a). Eleven aphid colonies observed exhibited an infection in the gut (Table S1). Aphids that exhibited this pattern belonged to *Aphis sp.* (mostly *A. fabae*), except one *Capitophorus elaeagni* individual. Infection of sheath cells pattern was also found in two colonies of *Macrosiphum* species (Table S1). In this case, the symbionts were located around primary bacteriocytes enclosing *B. aphidicola* (Figure 3E). *S. symbiotica* was also found in secondary bacteriocytes flanking the primary bacteriocytes containing *B. aphidicola* (Figure 3F-H). This pattern was observed in 8 aphid colonies belonging to 4 aphid species (*Macrosiphoniella millefolii, Periphyllus testudinaceus, Macrosiphum mordvilkoi,* and *A. fabae*, Table S1).

Regarding insects associated to aphid colonies, *S. symbiotica* has already been detected among ants, in the proventriculus, a specialized organ involved in food filtration (Renoz et al. 2018). Here, only one observed hoverfly larva showed an infection to *S. symbiotica* in the prothorax, at the beginning of the digestive tract (Figure 4). Images clearly suggest that *S. symbiotica* is accumulate into the tentorial bar, in the junction of the pump chamber with the foregut, near the true mouth (Figure 4).

**Figure 4.**
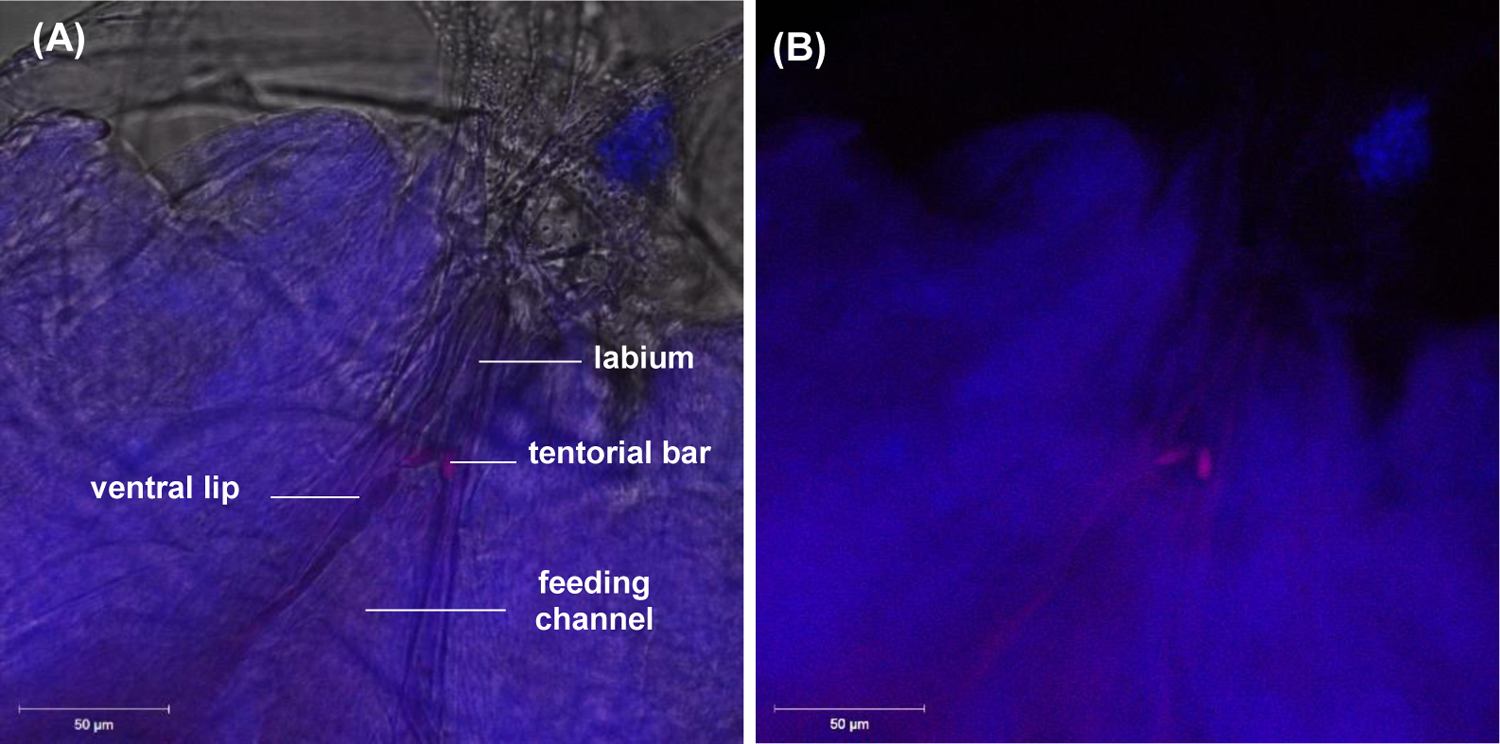
Whole-mount FISH of *S. symbiotica* in a naturally infected hoverfly larva (Prothorax, ventral view). Red Cy3 signals are *S. symbiotica*, and blue SYTOX Green signals are insect tissues. A is with bright field and B is without bright field. *S. symbiotica* is located at the level of the tentorial bar, near the true mouth (Reemer and Rotheray 2009).

### Diversity of *S. symbiotica* infections

Bacterial sequences of the *accD*, *gyrB*, *murE*, and *recJ* genes were taken from all 90 positive samples for *S. symbiotica* infection. Sequences were easily readable in 54 samples and for analyses, we excluded the remaining samples that had either polymorphic sequences, sequences that were difficult to read, or missing sequences (Table S2, Supplementary Information).

The phylogenetic relationship between these 54 *S. symbiotica* strains and other already described *S. symbiotica* strains was estimated using sequences obtained in this study, as well as sequences available in Genbank (Table S2). We also used *S. proteamaculans* 568, as outgroup. We found that the phylogeny of the *S. symbiotica* strains is strongly structured by the taxonomic identity of their host and the phylogenetic analyses established the existence of four distinct clades (Figure 5). Co-obligate *S. symbiotica* strains from *C. cedri* (SCc), *C. fornacula* (SCf), *C. strobi* (SCs), and *T. salignus* (STs Pazieg) aphids that were localized in secondary bacteriocytes form clade B (Burke and Moran 2011; Manzano-Marín and Latorre 2014; Manzano-Marín et al. 2017; Meseguer et al. 2017). The strains composing this clade are co-obligate symbionts of aphids and exhibit a long-term co-evolutionary history with their hosts. The clade A is composed of *S. symbiotica* strains that are generally considered as aphid facultative endosymbionts or recent co-obligates. This clade includes strains found in *A. pisum*: SAp Tucson strain localized in sheath cells (Burke and Moran 2011; Manzano-Marín et al. 2017) and SIS strain residing in secondary bacteriocytes and sheath cells (Nikoh et al. 2019). It also contains strains found in *C. tujafilina* (SCt VLC) (Burke and Moran 2011; Manzano-Marín et al. 2017) and *A. urticata* (AURT-53S) that are localized in secondary bacteriocytes, and in *M. carnosum* (MCAR-56S) localized in sheath cells (Monnin et al. 2020). It is also composed of strains found in *M. rosae* and *M. mordvilkoi* and localized in secondary bacteriocytes or sheath cells (Figure 3). The clade D forms a large monophyletic clade consisting of, among others, strains from aphids of the genus *Aphis*, and strains from the aphid species *M. absintii*, *M. millefolii*, *B. cardui* and *T. aurantia*. This clade also includes *S. symbiotica* strains from the host plants and insects associated with the aphid colonies. Interestingly, these strains fall into the same clade as *S. symbiotica* strains associated with aphids sampled in the corresponding colony. Also included in this clade, the three strains previously isolated (SCWBI-2.3^T^, SApa 8-A1, SAf 24.1) (Sabri et al. 2011; Grigorescu et al. 2017) and the strains previously detected in the aphid gut (FR65, FR28, FR35 and VP6; Table S4) (Renoz et al. 2018). All *S. symbiotica* strains localized in the aphid gut are grouped within this clade. However, some strains localized in secondary bacteriocytes (ID 33, 380, 369,197; Figure 3) are also included in this clade. We also observed a polytomy with a group containing strains from an aphid, an ant and a bug sampled in the same colony and another group containing the cultivable strain (SCWBI-2.3^T^) and two strains from the aphid gut. The clade C includes *S. symbiotica* strains asociated with aphids of the genus *Periphullus* and a strain associated with *U. sonchi.* These strains are considered as nutritional co-obligate (Monnin et al. 2020). In *P. testudinaceus*, *S. symbiotica* was clearly localized in bacteriocytes (Figure 3).

**Figure 5.**
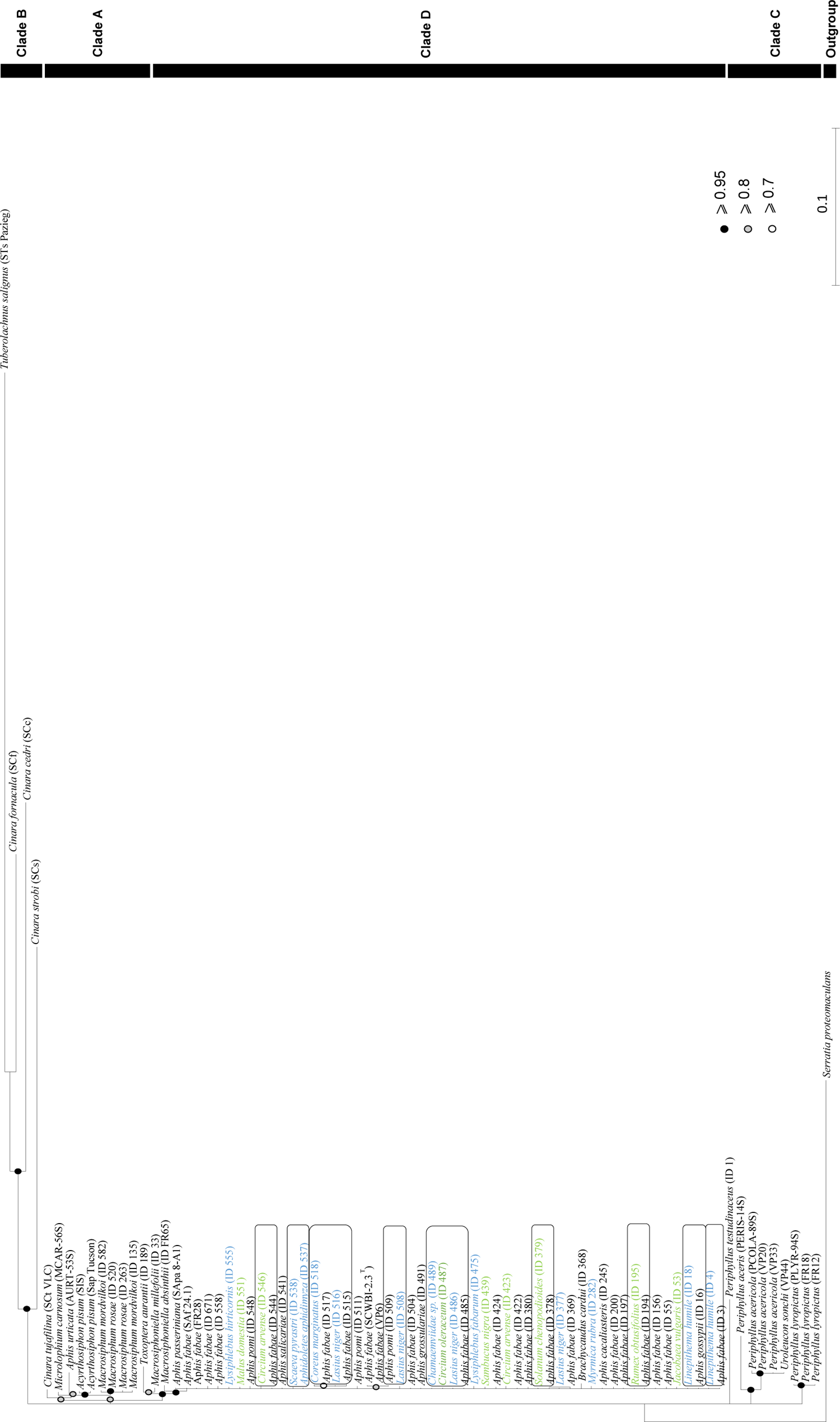
*Serratia symbiotica* phylogeny constructed using MrBayes analysis based on concatenated sequences of the *accD*, *gyrB*, *murE* and *recJ* genes. The names given for each terminal node reflect the taxonomic identity of the host from which the *S. symbiotica* strain was sequenced, followed by the names of the *S. symbiotica* strains. Circles on branches indicate Bayesian posterior probabilities: black circles for probabilities greater than or equal to 0.95, grey circles for probabilities between 0.94 and 0.8, and white circles for probabilities between 0.79 and 0.7. The black names indicate the aphid samples, the green names indicate the aphid host plant samples and the blue names the insect samples associated to the aphid colonies. Rectangles correspond to individuals belonging to the same colony. Clade D correspond to the clade D of the previous study (Renoz et al. 2018) grouping the strains isolated and exhibiting a free-living capacity in laboratory conditions as well as strains exhibiting aphid gut infection.

## DISCUSSION

In this study, we investigated the presence and the distribution of *S. symbiotica* in wild aphid populations, as well as in the different compartiments of the surrounding environment of these sap-feeding insects. Our results provide a comprehensive picture of the ubiquity of *S. symbiotica* in the natural aphid environment including ants, predators, parasitoids and plants. They first confirm that *S. symbiotica* exhibit different patterns of infection in aphids and that certain strains naturally transit through the digestive tract of these insects (Renoz et al. 2019). Our findings also demonstrate that the distribution of *S. symbiotica* is not limited to aphids but extends to other organisms that may interact with them and host plants. Finally, our results suggest that some *S. symbiotica* strains may be able to jump from one host to another, including plants, and could undergo frequent horizontal transfers. In the light of these results, we discuss the multi-faceted nature of *S. symbiotica* and the evolutionary scenarios of symbiont acquisitions in insects.

In a previous study, we experimentally demonstrated that a strain of *S. symbiotica* previously isolated from the aphid *Aphis fabae* (Sabri et al 2014) was capable to invade phloem sap of *Vicia faba* and that infected plants can then serve as reservoirs for horizontal transmission of *S. symbiotica* in aphids (Pons et al. 2019). Our field study suggests that *S. symbiotica* can naturally reside in plants. To our knowledge, it is the first time that this symbiont species was found in plants collected directly on the field. The presence of *S. symbiotica* in plants addresses major ecological issues including 1) the role of plants in the dissemination in insect populations of bacterial candidates, which may serve as environmental progenitors for the establishment of new mutualisms, and 2) the nature of the interaction that certain *S. symbiotica* strains have with plants. When *S. symbiotica* were detected in both aphids and the corresponding host plant, they are found in the same phylogenetic clades, indicating a close phylogenetic proximity whose suggests horizontal transfers between aphids and plants. The hypothesis of the existence of such transfers is strengthened by the fact that we found host plants harboring *S. symbiotica* significantly more frequently when aphids were also found infected on that plant. However, in some cases, we did not detect *S. symbiotica* in aphids when the host plant was positive. This may be because either aphids sampled had not yet been infected, we missed the infection, or some strains only infect plants and fail to infect aphids. Our results thus raise questions about the nature of the interaction between *S. symbiotica* and plants. The *Serratia* genus includes bacterial members that have the propancy to evolve in diverse environments including water, soil, plants, humans and invertebrates (Grimont and Grimont 2006), and many *Serratia* species can establish mutualistic or parasitic partnerships with plants (Petersen and Tisa 2013). In the context of our study, one assumption is that certain *S. symbiotica* strains are associated with plants, either in commensalistic associations, or in more intimate associations (pathogenic or mutualistic). Our previous study suggests that *S. symbiotica* can provide benefits to the infected plants, and that, in exchange, plants provide a suitable ecological niche mostly composed of sugar for *S. symbiotica* (Pons et al. 2019b). Ongoing studies should provide a better understanding of the nature of the interaction between certain bacterial strains and plants.

Another assumption is that *S. symbiotica* strains residing in plants could serve as an environmental symbiont pool from which new intimate associations are formed. This is supported by phylogenetic analyses showing that the *Cinara*-associated *Erwinia* co-obligate symbionts initially derived from plant associates (Manzano-Marín et al. 2020). Our field study thus suggests that *S. symbiotica* in aphids evolved from bacteria that originally inhabited plants, and highlights this bacterium as a valuable model for understanding how bacteria develop cross-kingdom host jumps and exploit multiple hosts before becoming, under still unclear circumstances, long-term mutualists in insects.

The broad sampling in our study highlights the great diversity of *S. symbiotica* strains in aphids with different tissue tropisms. We confirms the existence of strains that reside in the aphid gut, particularly in species of the genus *Aphis*, as already previously reported (Renoz et al. 2018). The existence of these strains that reside naturally within insect gut suggest that the feeding behavior is a likely source of new acquisition for these strains. Several studies support the idea that bacteria present in the aphid’s diet are a source of endosymbiotic bacteria (Harada et al. 1996; Manzano-Marín et al. 2020). Our results support this hypothesis and the following scenario: 1) acquisition of bacterial progenitors originally inhabiting the plant, 2) transition through the insect digestive tract, an intermediate step before, and 3) the possible establishment of more intimate associations. Experimental works have demonstrated that gut strains are unable to establish persistent maternal transmission (Pons et al. 2019, Perreau et al. 2020). A strict vertical transmission is, however, not always possible and in fact, not always necessary for the transmission of some gut symbionts. One of the most remarkable cases is the relationship between bean bug *Riptortus pedestris* and the beneficial gut symbiont *Burkholderia*. For successful inoculation of the next generation, the offspring must feed on inoculated or contaminated material from the adult’s gut flora, and the symbiont must be environmentally re-acquired with each generation through nymph feeding (Kikuchi et al. 2007; Kikuchi and Yumoto 2013). In the context of the interaction between aphid and these gut-associated *S. symbiotica*, a permanent environmental source of contamination is required to maintan the infection across aphid generations (Pons et al. 2019a). It is possible that infected plants provide such a source. These gut-associated strains were found to be grouped in a common phylogenetic clade (clade D), as previously shown in (Renoz et al. 2018). At the excpetion of the oleaster aphid *Capitophorus elaeagni*, aphid species harboring gut-associated *S. symbiotica* were members of the genus *Aphis*. If experimental studies have previously highlighted the moderate pathogenic effects of gut-associated *S. symbiotica* (CWBI-2.3^T^) (Pons et al. 2019, Perrau et al. 2020), their involvement in mutualistic associations with their insect host cannot be excluded. A recent study showed that the *S. symbiotica* strain (CWBI-2.3^T^) infecting the aphid gut is capable to produce proteases including metalloproteases, which may facilitate the digestion of plant proteins by helping to suppress plant defense (Skaljac et al. 2019). In light of this, it is possible that these gut-associated *S. symbiotica* contribute to improve the assimilation by aphids of some types of phloem sap or the elimination of toxins produced by host plants, allowing the aphid adaptation to feed on certain plants.

Our results offer the opportunity to investigate the different patterns of tissue tropism exhibited by *S. symbiotica* in aphids and beyond. It was previously established that intracellular *S. symbiotica* strains could be housed either in secondary bacteriocytes and in sheath cells in the case of facultative strains (Koga R. et al. 2003; Moran et al. 2005; Koga et al. 2012) and solely in bacteriocytes (subfamily Aphidinae subfamily) in the case of nutritional co-obligate strains (subfamilies Lachninae and Chaitophorinae subfamilies) (Manzano-Marín et al. 2016, 2017; Monnin et al. 2020). In parallel to these infection patterns, the aphid gut that evolve exclusively, to our knowledge, at the extracellular level (Renoz et al. 2018; Pons et al. 2019a; Perreau et al. 2020; Elston et al. 2021). All of these patterns could be observed from the sampling we carried out in this study. Intracellular *S. symbiotica* strains exclusively compartmentalized into sheath cells surrounding the primary bacteriocytes hosting *B. aphidicola* have only been observed in individuals belonging to *Macrosiphum* species. *S. symbiotica* was also observed in secondary bacteriocytes in different aphid species, including *Periphyllus testudinaceus, Macrosiphoniella millefolii, Aphis fabae, Macrosiphum mordvilkoi* and *Macrosiphum rosae*. So far, this tissue tropism of *S. symbiotica* has mostly been reported in aphids where the symbiont is a co-obligate nutritional partner (Manzano-Marín et al. 2016, 2017; Monnin et al. 2020). In our study, this is the case of *Periphyllus testudinaceus*, a member of the subfamily Chaitophorinae where *S. symbiotica* has evolved as a nutritional co-obligate partner (Monnin et al. 2020). This tissue tropism pattern has also been described in *A. pisum* but mainly after an artificial infection of the hemolymph (Fukatsu et al. 2000; Koga R. et al. 2003) and in *Aphis urticata* (Monnin et al. 2020). Our observations demonstrate that a specific microscopic approach is definitively needed to clarify the nature of the associations in which bacteria and insects are involved because the topology of the phylogenetic tree does not allow inferring the infection pattern displayed by *S. symbiotica*: the clade D remains diverse and includes strains that exhibit different tissue tropism patterns, suggesting a lack of resolution. It would thus be necessary to target additional housekeeping genes to increase the resolution of the phylogentic analyses, discriminate differences within clade D, and fully resolve the phylogeny of *S. symbiotica*.

While horizontal symbiont transfer is known to occur in the field, the rate at which it occurs is still unknown (Russell et al. 2003; Darby and Douglas 2003; Gehrer and Vorburger 2012; Jousselin et al. 2013; Henry et al. 2013; Guyomar et al. 2018). Here, we demonstrated that *S. symbiotica* infection can occur in non-aphid insect species found in close proximity to aphid colonies and involved in trophic relationships with them (12% of the screened insects were found positive to *S. symbiotica*). We found that these insects were significantly more likely to harbor *S. symbiotica* when aphid colonies were infected by the bacterium. In addition, we showed that *S. symbiotica* strains associated with the non-aphid insects were in the same clades as strains found in aphids of the corresponding sampled colonies. These results suggest that horizontal transfers of *S. symbiotica* can occur within natural insect communities and that *S. symbiotica* infections extend beyond aphids. This can be an adaptive characteristic allowing a wide dispersion of a fairly generalist bacterium although dependent on sugar rich diet. Ant-tending is a possible route of transmission of symbiotic bacteria from aphids to ants and vice versa, especially because ants feed on honeydew that can be infected by *S. symbiotica* (Pons et al 2019). With 10% of the sampled ants found infected by *S. symbiotica*, our results confirmed previous studies having reported the presence of the bacterium in these social insects (Sirvio and Pamilo 2010, Het et al. 2014, Renoz et al. 2019). This is an important issue because, even if *S. symbiotica* is only an occasional passenger in ants, the social and feeding behavior of these insects could promote the dissemination of certain *S. symbiotica* strains in the aphid colonies and increase the environmental symbiont pool from which potential new associations could originate. Another possible route of symbiont transmission from aphids to non-aphid insects is predation by entomophagous insects such as ladybugs, hoverflies, aphid midges and fly larvae. We showed that *S. symbiotica* infect these insects and we were able to localize the infection at the beginning of the digestive tract of a hoverfly larvae. This suggest that *S. symbiotica* can transit from one trophic level to another through predation and initiate an infection in non-aphid species. Further studies will be needed to determine the impact of *S. symbiotica* in these new hosts and the extent to which more systemic and persistent infection may occur. We also found that parasitoids could be infected, but in this case, the tissue tropism of *S. symbiotica* was not studied due to a lack of insect material to perform all analyses. A previous study showed the role of parasitoids as vectors of facultative bacterial symbionts in aphids (Gehrer and Vorburger 2012). These insects could promote the emergence of symbioses because their mode of parasitism could allow symbionts to have a direct access to the aphid hemolymph where the cells specialized in symbiont sheltering are found. Ongoing studies should clarify the nature of this interaction at the developmental level, but also at the ecological level with the cascading effects that *S. symbiotica* infections could induce on parasitoid communities. Indeed, *S. symbiotica*, whether facultative intracellular or gut-associated strains, can severely hamper the development of parasitoids in aphids (Pons et al. 2019a). *S. symbiotica* was also detected in bugs, caterpillars and moth larvae that are sap-sucking insects or herbivores that do not directly interact with aphids but can feed on the same host plants. These results suggest, once again, that plants could be reservoir at the origin of new interactions between *S. symbiotica* and insects. All these findings reveal the surprising ubiquity of *S. symbiotica,* with strains that are likely able to disperse through multiple insect species via horizontal transmission. Some bacterial genera show a high propensity to infect diverse hosts, including *Spiroplasma*, *Sodalis* and *Arsenophnus* symbionts (Fukatsu et al. 2001; Bressan et al. 2009; Clayton et al. 2012; Jousselin et al. 2013; Schwarz et al. 2014; Santos-Garcia et al. 2017; Ballinger et al. 2018; Masson et al. 2018). These exemples stresses that putative progenitors of host-beneficial symbionts circulate throughout the trophic system, with potential consequences not only for the ecology and evolution of their primary host, but also for those of transitive hosts (Moran and Yun 2015). In the case of *S. symbiotica* (especialy gut-associated strains), whether it is an established symbiont or whether a transient commensal resident in multiple insect hosts, is yet to be elucidated.

In our study, a total of 58 aphid species were sampled and the prevalence of *S. symbiotica* reaches 20% in aphids, which is fairly similar to what has been recorded in previous studies (Henry et al. 2015; Renoz et al. 2018). Our results confirmed the high presence of *S. symbiotica* infections in the genus *Aphis* with 29% of aphids being infected. In addition, *S. symbiotica* was identified in a significant proportion in other aphid genera, such as in *Macrosiphum sp.* and *Capitophorus sp.*, while no infection was detected in some aphid genera, such as in *Uroleuchon sp.*. The prevalence of *S. symbiotica* in aphid populations may depend on the aphid genus, but this assumption should be taken with caution. Indeed, one study reported no general effect of aphid phylogeny on the symbiont presence (Henry et al. 2015), although another study, which considered all aphid symbionts, suggests a genus-specific effect (Zytynska and Weisser 2016). The nature of the interaction between insects and symbiotic bacteria can also depend on abiotic and biotic factors including the presence of predators and parasitoids, the host plant and climatic factors (Henry et al. 2013; Oliver et al. 2014; Zytynska and Weisser 2016; McLean Ailsa H. C. et al. 2016). Our results suggest that the incidence of *S. symbiotica* infection increase with higher frequencies at higher seasonal temperatures. Some experimental studies have already shown that some strains of *S. symbiotica* can provide a protection against heat shock in aphids (Montllor et al. 2002; Russell and Moran 2006; Burke et al. 2009). These protective effects associated with *S. symbiotica* are also supported by the high prevalence of *S. symbiotica* reported in arid regions (Henry et al. 2013). All these findings underlined a possible link between temperature seasonality and the prevalence of *S. symbiotica* in aphid populations. The multifaceted nature of *S. symbiotica* must, however, be considered (a diversity that includes strains ranging from pathogens to strict mutualists). Previous studies reported the influence of host plants on the presence of facultative symbionts in aphids (Tsuchida et al. 2004; Brady and White 2013; Guidolin and Cônsoli 2017). In our sampling, the distribution of *S. symbiotica* is, however, uniform across collections of *A. fabae* regardless of host plants. More generally, we showed that the proportion of aphids (all species included) harboring *S. symbiotica* is not different according to their degree of plant specialization. This result is not consistent with another study, which showed that *S. symbiotica* is most commonly found in specialist aphids (Henry et al. 2015). The distribution of *S. symbiotica* in natural aphid populations thus seems difficult to explain by considering only one or some factors. In some cases, it appeared to be random and in other cases as a consequence of selection acting on specific associations.

To conclude, our study provides a comprehensive picture of the ubiquity of *S. symbiotica* in nature. It is now established that intracellular host-dependent symbionts of insects can evolve from originally free-living bacterial lineages (Husník et al. 2011; Clayton et al. 2012; Manzano-Marín et al. 2020). Genetic analyses indicate that these intimate relationships between insects and bacteria can evolve in a very dynamic fashion involving the recruitment of new bacterial partners and the repeated replacement of pre-existing intracellular symbionts (Koga and Moran 2014; Husnik and McCutcheon 2016; Meseguer et al. 2017; Monnin et al. 2020; Mao and Bennett 2020). Symbiont switching is an important evolutionary mechanism, which is not limited to insects (e.g., in reef organisms such as corals and sponges) (Webster and Reusch 2017), and by which maladaptive symbionts are replaced by new functional ones (Sudakaran et al. 2017). The presence of *S. symbiotica* at different trophic levels suggests the existence of an environmental pool of bacteria from which new intimate partnerships with insects may emerge. We hypothesize that the *S. symbiotica* diversity includes strains that exhibit a generalist lifestyle, still capable to develop cross-kingdom host jumps, before, under specific conditions, potentially transiting to a more specialized, and host-dependent lifestyle. A key aspect is understanding the factors that favor such a transition, in particular the conditions that promote the passage from the digestive tract to the hemolymph, a required step for the adoption of an intracellular life and for access to vertical transmission. Another burning question concerns the specific characteristics possessed by certain bacterial genera and species (e.g., members of the genera *Sodalis*, *Arsenophonus*, and members of *S. symbiotica*) to regularly engage in stable alliances with insects. The ecological versatility of *S. symbiotica* offers exciting avenues for answering these questions and refining our understanding of the evolution of bacterial mutualism.

## ACKNOWLEDGMENTS

We thank Charlotte Tinel, Aurélien Kaiser and Anastase Harelimana for some samples collection. We also thank Alain Vanderpoorten (University of Liège) for his help in phylogenetic analyses. This work was supported by the Fonds de la Recherche Scientifique (FNRS) through a Fonds pour la Formation à la Recherche dans l’Industrie et dans l’Agriculture (FRIA). This paper is publication BRC 348 of the Biodiversity Research Center (Universtité catholique de Louvain).

## REFERENCES

1. Aljanabi SM, Martinez I (1997) Universal and rapid salt-extraction of high quality genomic DNA for PCR-based techniques. Nucleic Acids Res 25:4692–4693. https://doi.org/10.1093/nar/25.22.4692

2. Ballinger MJ, Moore LD, Perlman SJ (2018) Evolution and Diversity of Inherited Spiroplasma Symbionts in Myrmica Ants. Appl Env Microbiol 84:e02299–17. https://doi.org/10.1128/AEM.02299-17

3. Baumann P, Moran NA, Baumann LC (2013) Bacteriocyte-Associated Endosymbionts of Insects. In: Rosenberg E, DeLong EF, Lory S, et al. (eds) The Prokaryotes: Prokaryotic Biology and Symbiotic Associations. Springer Berlin Heidelberg, Berlin, Heidelberg, pp 465–496

4. Brady CM, White JA (2013) Cowpea aphid (Aphis craccivora) associated with different host plants has different facultative endosymbionts. Ecol Entomol 38:433–437. https://doi.org/10.1111/een.12020

5. Bressan A, Sémétey O, Arneodo J, et al (2009) Vector Transmission of a Plant-Pathogenic Bacterium in the Arsenophonus Clade Sharing Ecological Traits with Facultative Insect Endosymbionts. Phytopathology 99:1289–1296. https://doi.org/10.1094/PHYTO-99-11-1289

6. Bright M, Bulgheresi S (2010) A complex journey: transmission of microbial symbionts. Nat Rev Microbiol 8:218–230. https://doi.org/10.1038/nrmicro2262

7. Burke G, Fiehn O, Moran N (2009) Effects of facultative symbionts and heat stress on the metabolome of pea aphids. ISME J 4:242–252. https://doi.org/10.1038/ismej.2009.114

8. Burke GR, Moran NA (2011) Massive Genomic Decay in Serratia symbiotica, a Recently Evolved Symbiont of Aphids. Genome Biol Evol 3:195–208. https://doi.org/10.1093/gbe/evr002

9. Caspi-Fluger Ayelet, Inbar Moshe, Mozes-Daube Netta, et al (2012) Horizontal transmission of the insect symbiont Rickettsia is plant-mediated. Proc R Soc B Biol Sci 279:1791–1796. https://doi.org/10.1098/rspb.2011.2095

10. Castresana J (2000) Selection of Conserved Blocks from Multiple Alignments for Their Use in Phylogenetic Analysis. Mol Biol Evol 17:540–552. https://doi.org/10.1093/oxfordjournals.molbev.a026334

11. Chari A, Oakeson KF, Enomoto S, et al (2015) Phenotypic characterization of Sodalis praecaptivus sp. nov., a close non-insect-associated member of the Sodalis-allied lineage of insect endosymbionts. Int J Syst Evol Microbiol 65:1400–1405. https://doi.org/10.1099/ijs.0.000091

12. Chong RA, Moran NA (2018) Evolutionary loss and replacement of Buchnera, the obligate endosymbiont of aphids. ISME J 12:898. https://doi.org/10.1038/s41396-017-0024-6

13. Clayton AL, Oakeson KF, Gutin M, et al (2012) A Novel Human-Infection-Derived Bacterium Provides Insights into the Evolutionary Origins of Mutualistic Insect–Bacterial Symbioses. PLOS Genet 8:e1002990. https://doi.org/10.1371/journal.pgen.1002990

14. D’acier AC, Cruaud Artige (2014) DNA Barcoding and the Associated PhylAphidB@se Website for the Identification of European Aphids (Insecta: Hemiptera: Aphididae). https://journals.plos.org/plosone/article?id=10.1371/journal.pone.0097620. Accessed 12 Apr 2019

15. Darby AC, Douglas AE (2003) Elucidation of the Transmission Patterns of an Insect-Borne Bacterium. Appl Environ Microbiol 69:4403–4407. https://doi.org/10.1128/AEM.69.8.4403-4407.2003

16. Douglas AE (1998) Nutritional Interactions in Insect-Microbial Symbioses: Aphids and Their Symbiotic Bacteria Buchnera. Annu Rev Entomol 43:17–37. https://doi.org/10.1146/annurev.ento.43.1.17

17. Douglas AE (2011) Lessons from Studying Insect Symbioses. Cell Host Microbe 10:359–367. https://doi.org/10.1016/j.chom.2011.09.001

18. Doyle J (1991) DNA Protocols for Plants. In: Molecular Techniques in Taxonomy. Springer, Berlin, Heidelberg, pp 283–293

19. Elston KM, Perreau J, Maeda GP, et al (2021) Engineering a Culturable Serratia symbiotica Strain for Aphid Paratransgenesis. Appl Environ Microbiol 87:. https://doi.org/10.1128/AEM.02245-20

20. Feldhaar H (2011) Bacterial symbionts as mediators of ecologically important traits of insect hosts. Ecol Entomol 36:533–543. https://doi.org/10.1111/j.1365-2311.2011.01318.x

21. Foray V, Grigorescu AS, Sabri A, et al (2014) Whole-Genome Sequence of Serratia symbiotica Strain CWBI-2.3T, a Free-Living Symbiont of the Black Bean Aphid Aphis fabae. Genome Announc 2:e00767–14. https://doi.org/10.1128/genomeA.00767-14

22. Fukatsu T, Nikoh N, Kawai R, Koga R (2000) The Secondary Endosymbiotic Bacterium of the Pea Aphid Acyrthosiphon pisum (Insecta: Homoptera). Appl Environ Microbiol 66:2748–2758. https://doi.org/10.1128/AEM.66.7.2748-2758.2000

23. Fukatsu T, Tsuchida T, Nikoh N, Koga R (2001) Spiroplasma Symbiont of the Pea Aphid, Acyrthosiphon pisum (Insecta: Homoptera). Appl Environ Microbiol 67:1284–1291. https://doi.org/10.1128/AEM.67.3.1284-1291.2001

24. Gehrer L, Vorburger C (2012) Parasitoids as vectors of facultative bacterial endosymbionts in aphids. Biol Lett rsbl20120144. https://doi.org/10.1098/rsbl.2012.0144

25. Gil R, Latorre A (2019) Unity Makes Strength: A Review on Mutualistic Symbiosis in Representative Insect Clades. Life 9:21. https://doi.org/10.3390/life9010021

26. Grigorescu AS, Renoz F, Sabri A, et al (2017) Accessing the Hidden Microbial Diversity of Aphids: an Illustration of How Culture-Dependent Methods Can Be Used to Decipher the Insect Microbiota. Microb Ecol 1–14. https://doi.org/10.1007/s00248-017-1092-x

27. Grimont F, Grimont PAD (2006) The Genus Serratia. In: Dworkin M, Falkow S, Rosenberg E, et al. (eds) The Prokaryotes. Springer New York, pp 219–244

28. Guidolin AS, Cônsoli FL (2017) Symbiont Diversity of Aphis (Toxoptera) citricidus (Hemiptera: Aphididae) as Influenced by Host Plants. Microb Ecol 73:201–210. https://doi.org/10.1007/s00248-016-0892-8

29. Guo J, Hatt S, He K, et al (2017) Nine facultative endosymbionts in aphids. A review. J Asia-Pac Entomol. https://doi.org/10.1016/j.aspen.2017.03.025

30. Guyomar C, Legeai F, Jousselin E, et al (2018) Multi-scale characterization of symbiont diversity in the pea aphid complex through metagenomic approaches. Microbiome. https://microbiomejournal.biomedcentral.com/articles/10.1186/s40168-018-0562-9. Accessed 25 Nov 2019

31. Harada H, Ishikawa H (1993) Gut microbe of aphid closely related to its intracellular symbiont. Biosystems 31:185–191. https://doi.org/10.1016/0303-2647(93)90048-H

32. Harada H, Oyaizu H, Ishikawa H (1996) A Consideration About the Origin of Aphid Intracellular Symbiont in Connection with Gut Bacterial Flora. J Gen Appl Microbiol 42:17–26. https://doi.org/10.2323/jgam.42.17

33. Harada H, Oyaizu H, Kosako Y, Ishikawa H (1997) Erwinia aphidicola, a new species isolated from pea aphid, Acyrthosiphon pisum. J Gen Appl Microbiol 43:349–354. https://doi.org/10.2323/jgam.43.349

34. Henry LM, Maiden MCJ, Ferrari J, Godfray HCJ (2015) Insect life history and the evolution of bacterial mutualism. Ecol Lett 18:516–525. https://doi.org/10.1111/ele.12425

35. Henry LM, Peccoud J, Simon J-C, et al (2013) Horizontally Transmitted Symbionts and Host Colonization of Ecological Niches. Curr Biol 23:1713–1717. https://doi.org/10.1016/j.cub.2013.07.029

36. Heyworth ER, Ferrari J (2015) A facultative endosymbiont in aphids can provide diverse ecological benefits. J Evol Biol 28:1753–1760. https://doi.org/10.1111/jeb.12705

37. Hosokawa T, Ishii Y, Nikoh N, et al (2016) Obligate bacterial mutualists evolving from environmental bacteria in natural insect populations. Nat Microbiol 1:15011. https://doi.org/10.1038/nmicrobiol.2015.11

38. Husník F, Chrudimský T, Hypša V (2011) Multiple origins of endosymbiosis within the Enterobacteriaceae (γ-Proteobacteria): convergence of complex phylogenetic approaches. BMC Biol 9:87. https://doi.org/10.1186/1741-7007-9-87

39. Husnik F, McCutcheon JP (2016) Repeated replacement of an intrabacterial symbiont in the tripartite nested mealybug symbiosis. Proc Natl Acad Sci 113:E5416–E5424. https://doi.org/10.1073/pnas.1603910113

40. Jousselin E, Cœur d’Acier A, Vanlerberghe-Masutti F, Duron O (2013) Evolution and diversity of Arsenophonus endosymbionts in aphids. Mol Ecol 22:260–270. https://doi.org/10.1111/mec.12092

41. Kearse M, Moir R, Wilson A, et al (2012) Geneious Basic: An integrated and extendable desktop software platform for the organization and analysis of sequence data. Bioinformatics 28:1647–1649. https://doi.org/10.1093/bioinformatics/bts199

42. Kikuchi Y, Hosokawa T, Fukatsu T (2007) Insect-Microbe Mutualism without Vertical Transmission: a Stinkbug Acquires a Beneficial Gut Symbiont from the Environment Every Generation. Appl Environ Microbiol 73:4308–4316. https://doi.org/10.1128/AEM.00067-07

43. Kikuchi Y, Yumoto I (2013) Efficient Colonization of the Bean Bug Riptortus pedestris by an Environmentally Transmitted Burkholderia Symbiont. Appl Environ Microbiol 79:2088–2091. https://doi.org/10.1128/AEM.03299-12

44. Koga R, Meng X-Y, Tsuchida T, Fukatsu T (2012) Cellular mechanism for selective vertical transmission of an obligate insect symbiont at the bacteriocyte–embryo interface. Proc Natl Acad Sci 109:E1230–E1237. https://doi.org/10.1073/pnas.1119212109

45. Koga R, Moran NA (2014) Swapping symbionts in spittlebugs: evolutionary replacement of a reduced genome symbiont. ISME J 8:1237–1246. https://doi.org/10.1038/ismej.2013.235

46. Koga R., Tsuchida T., Fukatsu T. (2003) Changing partners in an obligate symbiosis: a facultative endosymbiont can compensate for loss of the essential endosymbiont Buchnera in an aphid. Proc R Soc Lond B Biol Sci 270:2543–2550. https://doi.org/10.1098/rspb.2003.2537

47. Koga R, Tsuchida T, Fukatsu T (2009) Quenching autofluorescence of insect tissues for in situ detection of endosymbionts. Appl Entomol Zool 44:281–291. https://doi.org/10.1303/aez.2009.281

48. Lamelas A, Gosalbes MJ, Manzano-Marín A, et al (2011) Serratia symbiotica from the Aphid Cinara cedri: A Missing Link from Facultative to Obligate Insect Endosymbiont. PLoS Genet 7:e1002357. https://doi.org/10.1371/journal.pgen.1002357

49. Lanfear R, Calcott B, Ho SYW, Guindon S (2012) PartitionFinder: Combined Selection of Partitioning Schemes and Substitution Models for Phylogenetic Analyses. Mol Biol Evol 29:1695–1701. https://doi.org/10.1093/molbev/mss020

50. Latorre A, Manzano-Marín A (2017) Dissecting genome reduction and trait loss in insect endosymbionts. Ann N Y Acad Sci 1389:52–75. https://doi.org/10.1111/nyas.13222

51. Łukasik P, Guo H, Asch M van, et al (2015) Horizontal transfer of facultative endosymbionts is limited by host relatedness. Evolution 69:2757–2766. https://doi.org/10.1111/evo.12767

52. Manzano-Marín A, Coeur d’acier A, Clamens A-L, et al (2018) A Freeloader? The Highly Eroded Yet Large Genome of the Serratia symbiotica Symbiont of Cinara strobi. Genome Biol Evol 10:2178–2189. https://doi.org/10.1093/gbe/evy173

53. Manzano-Marín A, Latorre A (2016) Snapshots of a shrinking partner: Genome reduction in *Serratia symbiotica*. Sci Rep 6:32590. https://doi.org/10.1038/srep32590

54. Manzano-Marín A, Latorre A (2014) Settling Down: The Genome of Serratia symbiotica from the Aphid Cinara tujafilina Zooms in on the Process of Accommodation to a Cooperative Intracellular Life. Genome Biol Evol 6:1683–1698. https://doi.org/10.1093/gbe/evu133

55. Manzano-Marín A, Simon J-C, Latorre A (2016) Reinventing the Wheel and Making It Round Again: Evolutionary Convergence in Buchnera–Serratia Symbiotic Consortia between the Distantly Related Lachninae Aphids Tuberolachnus salignus and Cinara cedri. Genome Biol Evol 8:1440–1458. https://doi.org/10.1093/gbe/evw085

56. Manzano-Marín A, Szabó G, Simon J-C, et al (2017) Happens in the best of subfamilies: establishment and repeated replacements of co-obligate secondary endosymbionts within Lachninae aphids. Environ Microbiol 19:393–408. https://doi.org/10.1111/1462-2920.13633

57. Manzano-Marın A, Coeur D’acier A, Clamens A-L, et al (2020) Serial horizontal transfer of vitamin-biosynthetic genes enables the establishment of new nutritional symbionts in aphids’ di-symbiotic systems. ISME J 14:259–273. https://doi.org/10.1038/s41396-019-0533-6

58. Mao M, Bennett GM (2020) Symbiont replacements reset the co-evolutionary relationship between insects and their heritable bacteria. ISME J 14:1384–1395. https://doi.org/10.1038/s41396-020-0616-4

59. Masson F, Copete SC, Schüpfer F, et al (2018) In Vitro Culture of the Insect Endosymbiont Spiroplasma poulsonii Highlights Bacterial Genes Involved in Host-Symbiont Interaction. mBio 9:. https://doi.org/10.1128/mBio.00024-18

60. Matsuura Y, Moriyama M, Łukasik P, et al (2018) Recurrent symbiont recruitment from fungal parasites in cicadas. Proc Natl Acad Sci 115:E5970–E5979. https://doi.org/10.1073/pnas.1803245115

61. McFall-Ngai M, Hadfield MG, Bosch TCG, et al (2013) Animals in a bacterial world, a new imperative for the life sciences. Proc Natl Acad Sci 110:3229–3236. https://doi.org/10.1073/pnas.1218525110

62. McLean Ailsa H. C., Parker Benjamin J., Hrček Jan, et al (2016) Insect symbionts in food webs. Philos Trans R Soc B Biol Sci 371:20150325. https://doi.org/10.1098/rstb.2015.0325

63. Meseguer AS, Manzano-Marín A, d’Acier AC, et al (2017) Buchnera has changed flatmate but the repeated replacement of co-obligate symbionts is not associated with the ecological expansions of their aphid hosts. Mol Ecol 26:2363–2378. https://doi.org/10.1111/mec.13910

64. Monnin D, Jackson R, Kiers ET, et al (2020) Parallel Evolution in the Integration of a Co-obligate Aphid Symbiosis. Curr Biol 30:1949–1957.e6. https://doi.org/10.1016/j.cub.2020.03.011

65. Montllor CB, Maxmen A, Purcell AH (2002) Facultative bacterial endosymbionts benefit pea aphids Acyrthosiphon pisum under heat stress. Ecol Entomol 27:189–195. https://doi.org/10.1046/j.1365-2311.2002.00393.x

66. Moran NA (2007) Symbiosis as an adaptive process and source of phenotypic complexity. Proc Natl Acad Sci 104:8627–8633. https://doi.org/10.1073/pnas.0611659104

67. Moran NA, Baumann P (2000) Bacterial endosymbionts in animals. Curr Opin Microbiol 3:270–275. https://doi.org/10.1016/S1369-5274(00)00088-6

68. Moran NA, Russell JA, Koga R, Fukatsu T (2005) Evolutionary Relationships of Three New Species of Enterobacteriaceae Living as Symbionts of Aphids and Other Insects. Appl Environ Microbiol 71:3302–3310. https://doi.org/10.1128/AEM.71.6.3302-3310.2005

69. Moran NA, Yun Y (2015) Experimental replacement of an obligate insect symbiont. Proc Natl Acad Sci 112:2093–2096. https://doi.org/10.1073/pnas.1420037112

70. Nadarasah G, Stavrinides J (2011) Insects as alternative hosts for phytopathogenic bacteria. FEMS Microbiol Rev 35:555–575. https://doi.org/10.1111/j.1574-6976.2011.00264.x

71. Nikoh N, Koga R, Oshima K, et al (2019) Genome Sequence of “Candidatus Serratia symbiotica” Strain IS, a Facultative Bacterial Symbiont of the Pea Aphid Acyrthosiphon pisum. Microbiol Resour Announc 8:. https://doi.org/10.1128/MRA.00272-19

72. Nylander JAA, Wilgenbusch JC, Warren DL, Swofford DL (2008) AWTY (are we there yet?): a system for graphical exploration of MCMC convergence in Bayesian phylogenetics. Bioinformatics 24:581–583. https://doi.org/10.1093/bioinformatics/btm388

73. Oliver KM, Degnan PH, Burke GR, Moran NA (2010) Facultative Symbionts in Aphids and the Horizontal Transfer of Ecologically Important Traits. Annu Rev Entomol 55:247–266. https://doi.org/10.1146/annurev-ento-112408-085305

74. Oliver KM, Russell JA, Moran NA, Hunter MS (2003) Facultative bacterial symbionts in aphids confer resistance to parasitic wasps. Proc Natl Acad Sci 100:1803–1807. https://doi.org/10.1073/pnas.0335320100

75. Oliver KM, Smith AH, Russell JA (2014) Defensive symbiosis in the real world – advancing ecological studies of heritable, protective bacteria in aphids and beyond. Funct Ecol 28:341–355. https://doi.org/10.1111/1365-2435.12133

76. Perreau J, Patel DJ, Anderson H, et al (2020) Vertical transmission at the pathogen-symbiont interface: Serratia symbiotica and aphids. bioRxiv 2020.09.01.279018. https://doi.org/10.1101/2020.09.01.279018

77. Petersen LM, Tisa LS (2013) Friend or foe? A review of the mechanisms that drive Serratia towards diverse lifestyles. Can J Microbiol 59:627–640. https://doi.org/10.1139/cjm-2013-0343

78. Pons I, Renoz F, Noël C, Hance T (2019a) New Insights into the Nature of Symbiotic Associations in Aphids: Infection Process, Biological Effects and Transmission Mode of Cultivable Serratia symbiotica Bacteria. Appl Env Microbiol AEM.02445–18. https://doi.org/10.1128/AEM.02445-18

79. Pons I, Renoz F, Noël C, Hance T (2019b) Circulation of the Cultivable Symbiont Serratia symbiotica in Aphids Is Mediated by Plants. Front. Microbiol.

80. Reemer M, Rotheray G (2009) Pollen feeding larvae in the presumed predatory syrphine genus Toxomerus Macquart (Diptera, Syrphidae). J. Nat. Hist.

81. Renoz F, Ambroise J, Bearzatto B, et al (2020) Draft Genome Sequences of Two Cultivable Strains of the Bacterial Symbiont Serratia symbiotica. Microbiol Resour Announc 9:. https://doi.org/10.1128/MRA.01579-19

82. Renoz F, Pons I, Vanderpoorten A, et al (2018) Evidence for Gut-Associated Serratia symbiotica in Wild Aphids and Ants Provides New Perspectives on the Evolution of Bacterial Mutualism in Insects. Microb Ecol. https://doi.org/10.1007/s00248-018-1265-2

83. Ronquist F, Teslenko M, van der Mark P, et al (2012) MrBayes 3.2: Efficient Bayesian Phylogenetic Inference and Model Choice Across a Large Model Space. Syst Biol 61:539–542. https://doi.org/10.1093/sysbio/sys029

84. Russell JA, Latorre A, Sabater-Muñoz B, et al (2003) Side-stepping secondary symbionts: widespread horizontal transfer across and beyond the Aphidoidea. Mol Ecol 12:1061–1075

85. Russell JA, Moran NA (2006) Costs and benefits of symbiont infection in aphids: variation among symbionts and across temperatures. Proc Biol Sci 273:603–610. https://doi.org/10.1098/rspb.2005.3348

86. Sabri A, Leroy P, Haubruge E, et al (2011) Isolation, pure culture and characterization of Serratia symbiotica sp. nov., the R-type of secondary endosymbiont of the black bean aphid Aphis fabae. Int J Syst Evol Microbiol 61:2081–2088. https://doi.org/10.1099/ijs.0.024133-0

87. Salem H, Florez L, Gerardo N, Kaltenpoth M (2015) An out-of-body experience: the extracellular dimension for the transmission of mutualistic bacteria in insects. Proc R Soc Lond B Biol Sci 282: 20142957. https://doi.org/10.1098/rspb.2014.2957

88. Santos-Garcia D, Silva FJ, Morin S, et al (2017) The All-Rounder Sodalis: A New Bacteriome-Associated Endosymbiont of the Lygaeoid Bug Henestaris halophilus (Heteroptera: Henestarinae) and a Critical Examination of Its Evolution. Genome Biol Evol 9:2893–2910. https://doi.org/10.1093/gbe/evx202

89. Schwarz RS, Teixeira ÉW, Tauber JP, et al (2014) Honey bee colonies act as reservoirs for two Spiroplasma facultative symbionts and incur complex, multiyear infection dynamics. MicrobiologyOpen 3:341–355. https://doi.org/10.1002/mbo3.172

90. Skaljac M, Vogel H, Wielsch N, et al (2019) Transmission of a Protease-Secreting Bacterial Symbiont Among Pea Aphids via Host Plants. Front Physiol 10:. https://doi.org/10.3389/fphys.2019.00438

91. Su Q, Oliver KM, Pan H, et al (2013) Facultative Symbiont Hamiltonella Confers Benefits to Bemisia tabaci (Hemiptera: Aleyrodidae), an Invasive Agricultural Pest Worldwide. Environ Entomol 42:1265–1271. https://doi.org/10.1603/EN13182

92. Sudakaran S, Kost C, Kaltenpoth M (2017) Symbiont Acquisition and Replacement as a Source of Ecological Innovation. Trends Microbiol 25:375–390. https://doi.org/10.1016/j.tim.2017.02.014

93. Takeshita K, Kikuchi Y (2017) Riptortus pedestris and Burkholderia symbiont: an ideal model system for insect–microbe symbiotic associations. Res Microbiol 168:175–187. https://doi.org/10.1016/j.resmic.2016.11.005

94. Tsuchida T, Koga R, Fukatsu T (2004) Host plant specialization governed by facultative symbiont. Science 303:1989. https://doi.org/10.1126/science.1094611

95. Vorburger C, Gouskov A (2011) Only helpful when required: a longevity cost of harbouring defensive symbionts. J Evol Biol 24:1611–1617. https://doi.org/10.1111/j.1420-9101.2011.02292.x

96. Webster NS, Reusch TBH (2017) Microbial contributions to the persistence of coral reefs. ISME J 11:2167– 2174. https://doi.org/10.1038/ismej.2017.66

97. Wernegreen JJ (2017) In it for the long haul: evolutionary consequences of persistent endosymbiosis. Curr Opin Genet Dev 47:83–90. https://doi.org/10.1016/j.gde.2017.08.006

98. Zytynska SE, Tighiouart K, Frago E (2021) The benefits and costs of hosting facultative symbionts in plant-sucking insects: a meta-analysis. Mol Ecol n/a: https://doi.org/10.1111/mec.15897

99. Zytynska SE, Weisser WW (2016) The natural occurrence of secondary bacterial symbionts in aphids. Ecol Entomol 41:13–26. https://doi.org/10.1111/een.12281

