## Supporting information for "Ubiquity of the Symbiont *Serratia symbiotica* in the Aphid Natural Environment: Distribution, Diversity and Evolution at a Multitrophic Level"

**Host**

**Alternating**

**Polyphagous**

**Restricted**

**Specialized**

Proportion of aphids harboring *S. symbiotica*

0

0.1

0.2

0.3

0.4

0.5

a

a

a

a

**Figure S1.** Prevalence of *S. symbiotica* in aphid species that differ in degrees of feeding specialization: host alternation (N=31), polyphagous (N=105), restricted (N=91), and specialized (N=13). Error bars depict the standard error. The lowercase letter show no significant differences.

**Table S1.** Detailed results of the screen of aphid species, the associated insects, and plant samples for the presence of *S. symbiotica*. Stars correspond to samples that were observed using fluorescence in situ hybridization. Tmean corresponds to the mean daily temperatures and Tmax corresponds to the maximum daily temperatures. Mtmean and Mtmax correspond to temperature data pooled into 6 categories.

| **Colony ID** | **Sampling ID** | **Samples** | **Host plant species** | **Insect species** | **Feeding specialization** | **Subfamily** | **Tribe** | **Sampling date** | **Collection site** | **Mtmean** | **Tmean** | **Mtmax** | **Tmax** | ***Serratia* infection** |
| --- | --- | --- | --- | --- | --- | --- | --- | --- | --- | --- | --- | --- | --- | --- |
| 1 | ID 1 * | Aphids | *Acer platanoides* | *Periphyllus testudinaceus* | restricted | Chaitophorinae | Chaitophorini | 14/05/2018 | Louvain-la-Neuve, Belgium | 13.9 | 12.9 | 18.2 | 17.7 | + |
| 1 | ID 2 | Ants | *Acer platanoides* | *Lasius niger* | / | Formicinae | Lasiini | 14/05/2018 | Louvain-la-Neuve, Belgium | 13.9 | 12.10 | 18.2 | 17.8 | - |
| 2 | ID 3 * | Aphids | *Rumex obtusifolius* | *Aphis fabae* | polyphagous | Aphidinae | Aphidini | 27/05/2018 | Levanto, Italy | NA | NA | NA | NA | + |
| 2 | ID 4 | Ants | *Rumex obtusifolius* | *Linepithema humile* | / | Dolichoderinae | Dolichoderini | 27/05/2018 | Levanto, Italy | NA | NA | NA | NA | + |
| 3 | ID 5 | Aphids | *Centranthus ruber* | *Aphis fabae* | polyphagous | Aphidinae | Aphidini | 28/05/2018 | Monteroso, Italy | NA | NA | NA | NA | - |
| 3 | ID 6 | Ants | *Centranthus ruber* | *Camponotus lateralis* | / | Formicinae | Camponotini | 28/05/2018 | Monteroso, Italy | NA | NA | NA | NA | - |
| 4 | ID 7 | Aphids | *Cirsium arvense* | *Aphis fabae* | polyphagous | Aphidinae | Aphidini | 28/05/2018 | Monteroso, Italy | NA | NA | NA | NA | - |
| 4 | ID 8 | Ants | *Cirsium arvense* | *Camponotus sylvaticus* | / | Formicinae | Camponotini | 28/05/2018 | Monteroso, Italy | NA | NA | NA | NA | - |
| 5 | ID 9 | Aphids | *Lathyrus odoratus* | *Megoura viciae* | restricted | Aphidinae | Macrosiphini | 28/05/2018 | Vernazza,Italy | NA | NA | NA | NA | - |
| 6 | ID 10 | Aphids | *Cucurbita pepo* | *Uroleucon jaceae* | restricted | Aphidinae | Macrosiphini | 28/05/2018 | Vernazza,Italy | NA | NA | NA | NA | - |
| 7 | ID 11 | Aphids | *Picris hieracioides* | *Uroleucon sonchi* | restricted | Aphidinae | Macrosiphini | 28/05/2018 | Corniglia, Italy | NA | NA | NA | NA | - |
| 7 | ID 12 | Ants | *Picris hieracioides* | *Lasius niger* | / | Formicinae | Lasiini | 28/05/2018 | Corniglia, Italy | NA | NA | NA | NA | - |
| 8 | ID 13 | Aphids | *Senecio vulgaris* | *Sitobion avenae* | polyphagous | Aphidinae | Macrosiphini | 28/05/2018 | Corniglia, Italy | NA | NA | NA | NA | - |
| 9 | ID 14 | Aphids | *Cirsium arvense* | *Aphis fabae* | polyphagous | Aphidinae | Aphidini | 28/05/2018 | Corniglia, Italy | NA | NA | NA | NA | - |
| 9 | ID 15 | Ants | *Cirsium arvense* | *Camponotus lateralis* | / | Formicinae | Camponotini | 28/05/2018 | Corniglia, Italy | NA | NA | NA | NA | - |
| 10 | ID 16 | Aphids | *Malva sylvestris* | *Aphis gossypii* | polyphagous | Aphidinae | Aphidini | 28/05/2018 | Levanto, Italy | NA | NA | NA | NA | + |
| 10 | ID 17 | Aphids | *Malva sylvestris* | *Acyrthosiphon malvae* | polyphagous | Aphidinae | Macrosiphini | 28/05/2018 | Levanto, Italy | NA | NA | NA | NA | - |
| 10 | ID 18 | Ants | *Malva sylvestris* | *Linepithema humile* | / | Dolichoderinae | Dolichoderini | 28/05/2018 | Levanto, Italy | NA | NA | NA | NA | + |
| 11 | ID 19 | Aphids | *Nerium oleander* | *Aphis nerri* | polyphagous | Aphidinae | Aphidini | 30/05/2018 | Pisa, Italy | NA | NA | NA | NA | - |
| 12 | ID 20 | Aphids | *Rumex obtusifolius* | *Aphis rumicis* | restricted | Aphidinae | Aphidini | 14/06/2018 | Havelange, Belgium | 13.9 | 15.5 | 18.2 | 19.2 | - |
| 12 | ID 21 | Ants | *Rumex obtusifolius* | *Lasius niger* | / | Formicinae | Lasiini | 14/06/2018 | Havelange, Belgium | 13.9 | 15.5 | 18.2 | 19.2 | - |
| 12 | ID 22 | Leaves | *Rumex obtusifolius* | / | / | / | / | 14/06/2018 | Havelange, Belgium | 13.9 | 15.5 | 18.2 | 19.2 | - |
| 12 | ID 23 | Stems | *Rumex obtusifolius* | / | / | / | / | 14/06/2018 | Havelange, Belgium | 13.9 | 15.5 | 18.2 | 19.2 | - |
| 12 | ID 24 | Hoverfly Larvae | *Rumex obtusifolius* | *Episyrphus balteatus* | / | Syrphinae | Syrphini | 14/06/2018 | Havelange, Belgium | 13.9 | 15.5 | 18.2 | 19.2 | - |
| 13 | ID 26 | Aphids | *Cirsium arvense* | *Aphis fabae* | polyphagous | Aphidinae | Aphidini | 14/06/2018 | Havelange, Belgium | 13.9 | 15.5 | 18.2 | 19.2 | - |
| 13 | ID 27 | Ants | *Cirsium arvense* | *Myrmica rubra* | / | Myrmicinae | Myrmicini | 14/06/2018 | Havelange, Belgium | 13.9 | 15.5 | 18.2 | 19.2 | - |
| 13 | ID 28 | Stems | *Cirsium arvense* | / | / | / | / | 14/06/2018 | Havelange, Belgium | 13.9 | 15.5 | 18.2 | 19.2 | - |
| 14 | ID 30 | Aphids | *Crepis capillaris* | *Uroleucon hypochoeridis* | restricted | Aphidinae | Macrosiphini | 14/06/2018 | Havelange, Belgium | 13.9 | 15.5 | 18.2 | 19.2 | - |
| 14 | ID 31 | Stems | *Crepis capillaris* | / | / | / | / | 14/06/2018 | Havelange, Belgium | 13.9 | 15.5 | 18.2 | 19.2 | - |
| 15 | ID 33 * | Aphids | *Achillea millefolium* | *Macrosiphoniella millefolii* | specialized | Aphidinae | Macrosiphini | 14/06/2018 | Havelange, Belgium | 13.9 | 15.5 | 18.2 | 19.2 | + |
| 15 | ID 34 | Stems | *Achillea millefolium* | / | / | / | / | 14/06/2018 | Havelange, Belgium | 13.9 | 15.5 | 18.2 | 19.2 | - |
| 16 | ID 36 | Aphids | *Triticum aestivum* | *Sitobion avenae* | polyphagous | Aphidinae | Macrosiphini | 14/06/2018 | Havelange, Belgium | 13.9 | 15.5 | 18.2 | 19.2 | - |
| 17 | ID 37 | Aphids | *Triticum aestivum* | *Metopolophium dirhodum* | host alternating | Aphidinae | Macrosiphini | 14/06/2018 | Havelange, Belgium | 13.9 | 15.5 | 18.2 | 19.2 | - |
| 18 | ID 38 | Aphids | *Triticum aestivum* | *Rhopalosiphum padi* | host alternating | Aphidinae | Aphidini | 14/06/2018 | Havelange, Belgium | 13.9 | 15.5 | 18.2 | 19.2 | - |
| 19 | ID 39 | Aphids | *Vicia faba* | *Aphis fabae* | polyphagous | Aphidinae | Aphidini | 14/06/2018 | Havelange, Belgium | 13.9 | 15.5 | 18.2 | 19.2 | - |
| 19 | ID 40 | Ants | *Vicia faba* | *Lasius niger* | / | Formicinae | Lasiini | 14/06/2018 | Havelange, Belgium | 13.9 | 15.5 | 18.2 | 19.2 | - |
| 19 | ID 41 | Stems | *Vicia faba* | / | / | / | / | 14/06/2018 | Havelange, Belgium | 13.9 | 15.5 | 18.2 | 19.2 | - |
| 20 | ID 43 | Aphids | *Centaurea cyanus* | *Uroleucon jaceae* | restricted | Aphidinae | Macrosiphini | 14/06/2018 | Havelange, Belgium | 13.9 | 15.5 | 18.2 | 19.2 | - |
| 20 | ID 44 | Ants | *Centaurea cyanus* | *Lasius niger* | / | Formicinae | Lasiini | 14/06/2018 | Havelange, Belgium | 13.9 | 15.5 | 18.2 | 19.2 | - |
| 21 | ID 46 | Aphids | *Triticum aestivum* | *Metopolophium dirhodum* | host alternating | Aphidinae | Macrosiphini | 14/06/2018 | Havelange, Belgium | 13.9 | 15.5 | 18.2 | 19.2 | - |
| 22 | ID 47 | Ants | *Rumex obtusifolius* | *Lasius niger* | / | Formicinae | Lasiini | 14/06/2018 | Louvain-la-Neuve, Belgium | 13.9 | 15.5 | 18.2 | 19.2 | - |
| 22 | ID 48 | Aphids | *Rumex obtusifolius* | *Aphis fabae* | polyphagous | Aphidinae | Aphidini | 14/06/2018 | Louvain-la-Neuve, Belgium | 13.9 | 15.5 | 18.2 | 19.2 | - |
| 22 | ID 49 | Stems | *Rumex obtusifolius* | / | / | / | / | 14/06/2018 | Louvain-la-Neuve, Belgium | 13.9 | 15.5 | 18.2 | 19.2 | - |
| 23 | ID 51 | Aphids | *Jacobaea vulgaris* | *Aphis fabae* | polyphagous | Aphidinae | Aphidini | 14/06/2018 | Louvain-la-Neuve, Belgium | 13.9 | 15.5 | 18.2 | 19.2 | - |
| 23 | ID 52 | Ants | *Jacobaea vulgaris* | *Lasius niger* | / | Formicinae | Lasiini | 14/06/2018 | Louvain-la-Neuve, Belgium | 13.9 | 15.5 | 18.2 | 19.2 | - |
| 23 | ID 53 | Stems | *Jacobaea vulgaris* | / | / | / | / | 14/06/2018 | Louvain-la-Neuve, Belgium | 13.9 | 15.5 | 18.2 | 19.2 | + |
| 24 | ID 55 | Aphids | *Leucanthemum vulgare* | *Aphis fabae* | polyphagous | Aphidinae | Aphidini | 14/06/2018 | Louvain-la-Neuve, Belgium | 13.9 | 15.5 | 18.2 | 19.2 | + |
| 24 | ID 56 | Ants | *Leucanthemum vulgare* | *Lasius niger* | / | Formicinae | Lasiini | 14/06/2018 | Louvain-la-Neuve, Belgium | 13.9 | 15.5 | 18.2 | 19.2 | - |
| 24 | ID 57 | Stems | *Leucanthemum vulgare* | / | / | / | / | 14/06/2018 | Louvain-la-Neuve, Belgium | 13.9 | 15.5 | 18.2 | 19.2 | - |
| 25 | ID 59 | Aphids | *Cirsium vulgare* | *Aphis fabae* | polyphagous | Aphidinae | Aphidini | 14/06/2018 | Louvain-la-Neuve, Belgium | 13.9 | 15.5 | 18.2 | 19.2 | - |
| 25 | ID 60 | Ants | *Cirsium vulgare* | *Lasius niger* | / | Formicinae | Lasiini | 14/06/2018 | Louvain-la-Neuve, Belgium | 13.9 | 15.5 | 18.2 | 19.2 | - |
| 25 | ID 61 | Stems | *Cirsium vulgare* | / | / | / | / | 14/06/2018 | Louvain-la-Neuve, Belgium | 13.9 | 15.5 | 18.2 | 19.2 | + |
| 26 | ID 63 | Aphids | *Cirsium vulgare* | *Aphis fabae* | polyphagous | Aphidinae | Aphidini | 14/06/2018 | Louvain-la-Neuve, Belgium | 13.9 | 15.5 | 18.2 | 19.2 | - |
| 26 | ID 64 | Aphids | *Cirsium vulgare* | NA | NA | NA | NA | 14/06/2018 | Louvain-la-Neuve, Belgium | 13.9 | 15.5 | 18.2 | 19.2 | - |
| 26 | ID 65 | Ants | *Cirsium vulgare* | *Myrmica scabrinodis* | / | Myrmicinae | Myrmicini | 14/06/2018 | Louvain-la-Neuve, Belgium | 13.9 | 15.5 | 18.2 | 19.2 | - |
| 26 | ID 66 | Stems | *Cirsium vulgare* | / | / | / | / | 14/06/2018 | Louvain-la-Neuve, Belgium | 13.9 | 15.5 | 18.2 | 19.2 | - |
| 27 | ID 68 | Aphids | *Leucanthemum vulgare* | *Aphis fabae* | polyphagous | Aphidinae | Aphidini | 15/06/2018 | Louvain-la-Neuve, Belgium | 17.9 | 17.7 | 23 | 23.6 | - |
| 27 | ID 69 | Ants | *Leucanthemum vulgare* | *Lasius niger* | / | Formicinae | Lasiini | 15/06/2018 | Louvain-la-Neuve, Belgium | 17.9 | 17.7 | 23 | 23.6 | - |
| 27 | ID 70 | Stems | *Leucanthemum vulgare* | / | / | / | / | 15/06/2018 | Louvain-la-Neuve, Belgium | 17.9 | 17.7 | 23 | 23.6 | - |
| 28 | ID 72 | Aphids | *Rumex obtusifolius* | *Aphis fabae* | polyphagous | Aphidinae | Aphidini | 15/06/2018 | Louvain-la-Neuve, Belgium | 17.9 | 17.7 | 23 | 23.6 | - |
| 28 | ID 73 | Ants | *Rumex obtusifolius* | *Lasius niger* | / | Formicinae | Lasiini | 15/06/2018 | Louvain-la-Neuve, Belgium | 17.9 | 17.7 | 23 | 23.6 | - |
| 28 | ID 74 | Stems | *Rumex obtusifolius* | / | / | / | / | 15/06/2018 | Louvain-la-Neuve, Belgium | 17.9 | 17.7 | 23 | 23.6 | - |
| 29 | ID 76 | Aphids | *Epipactis helleborine* | *Aphis fabae* | polyphagous | Aphidinae | Aphidini | 15/06/2018 | Louvain-la-Neuve, Belgium | 17.9 | 17.7 | 23 | 23.6 | - |
| 29 | ID 77 | Ants | *Epipactis helleborine* | *Lasius niger* | / | Formicinae | Lasiini | 15/06/2018 | Louvain-la-Neuve, Belgium | 17.9 | 17.7 | 23 | 23.6 | - |
| 29 | ID 78 | Stems | *Epipactis helleborine* | / | / | / | / | 15/06/2018 | Louvain-la-Neuve, Belgium | 17.9 | 17.7 | 23 | 23.6 | - |
| 30 | ID 80 | Aphids | *Campanula rapunculus* | *Macrosiphoniella sp.* | NA | Aphidinae | Macrosiphini | 15/06/2018 | Louvain-la-Neuve, Belgium | 17.9 | 17.7 | 23 | 23.6 | - |
| 30 | ID 81 | Stems | *Campanula rapunculus* | / | / | / | / | 15/06/2018 | Louvain-la-Neuve, Belgium | 17.9 | 17.7 | 23 | 23.6 | - |
| 31 | ID 83 | Aphids | *Artemisia annua* | *Macrosiphoniella artemisiae* | specialized | Aphidinae | Macrosiphini | 15/06/2018 | Louvain-la-Neuve, Belgium | 17.9 | 17.7 | 23 | 23.6 | - |
| 31 | ID 84 | Stems | *Artemisia annua* | / | / | / | / | 15/06/2018 | Louvain-la-Neuve, Belgium | 17.9 | 17.7 | 23 | 23.6 | - |
| 32 | ID 86 | Ants | *Crataegus monogyna* | *Lasius niger* | / | Formicinae | Lasiini | 15/06/2018 | Louvain-la-Neuve, Belgium | 17.9 | 17.7 | 23 | 23.6 | - |
| 32 | ID 87 | Aphids | *Crataegus monogyna* | *Aphis fabae* | polyphagous | Aphidinae | Aphidini | 15/06/2018 | Louvain-la-Neuve, Belgium | 17.9 | 17.7 | 23 | 23.6 | - |
| 32 | ID 88 | Stems | *Crataegus monogyna* | / | / | / | / | 15/06/2018 | Louvain-la-Neuve, Belgium | 17.9 | 17.7 | 23 | 23.6 | - |
| 33 | ID 89 | Aphids | *Cirsium vulgare* | *Aphis fabae* | polyphagous | Aphidinae | Aphidini | 15/06/2018 | Louvain-la-Neuve, Belgium | 17.9 | 17.7 | 23 | 23.6 | - |
| 33 | ID 90 | Ants | *Cirsium vulgare* | *Lasius niger* | / | Formicinae | Lasiini | 15/06/2018 | Louvain-la-Neuve, Belgium | 17.9 | 17.7 | 23 | 23.6 | - |
| 33 | ID 91 | Stems | *Cirsium vulgare* | / | / | / | / | 15/06/2018 | Louvain-la-Neuve, Belgium | 17.9 | 17.7 | 23 | 23.6 | - |
| 34 | ID 92 | Aphids | *Cirsium vulgare* | *Brachycaudus cardui* | restricted | Aphidinae | Macrosiphini | 15/06/2018 | Louvain-la-Neuve, Belgium | 17.9 | 17.7 | 23 | 23.6 | - |
| 34 | ID 93 | Stems | *Cirsium vulgare* | / | / | / | / | 15/06/2018 | Louvain-la-Neuve, Belgium | 17.9 | 17.7 | 23 | 23.6 | - |
| 35 | ID 94 | Aphids | *Tanacetum vulgare* | *Aphis fabae* | polyphagous | Aphidinae | Aphidini | 15/06/2018 | Louvain-la-Neuve, Belgium | 17.9 | 17.7 | 23 | 23.6 | - |
| 35 | ID 95 | Ants | *Tanacetum vulgare* | *Lasius niger* | / | Formicinae | Lasiini | 15/06/2018 | Louvain-la-Neuve, Belgium | 17.9 | 17.7 | 23 | 23.6 | - |
| 35 | ID 96 | Stems | *Tanacetum vulgare* | / | / | / | / | 15/06/2018 | Louvain-la-Neuve, Belgium | 17.9 | 17.7 | 23 | 23.6 | - |
| 35 | ID 97 | Ladybug larvae | *Tanacetum vulgare* | *Harmonia axyridis* | / | Coccinellinae | Coccinellini | 15/06/2018 | Louvain-la-Neuve, Belgium | 17.9 | 17.7 | 23 | 23.6 | - |
| 36 | ID 99 | Aphids | *Lythrum salicaria* | *Myzus lythri* | host alternating | Aphidinae | Macrosiphini | 15/06/2018 | Louvain-la-Neuve, Belgium | 17.9 | 17.7 | 23 | 23.6 | - |
| 36 | ID 100 | Stems | *Lythrum salicaria* | / | / | / | / | 15/06/2018 | Louvain-la-Neuve, Belgium | 17.9 | 17.7 | 23 | 23.6 | - |
| 37 | ID 102 | Aphids | *Artemisia annua* | *Macrosiphoniella artemisiae* | specialized | Aphidinae | Macrosiphini | 15/06/2018 | Louvain-la-Neuve, Belgium | 17.9 | 17.7 | 23 | 23.6 | - |
| 37 | ID 103 | Ladybugs | *Artemisia annua* | *Harmonia axyridis* | / | Coccinellinae | Coccinellini | 15/06/2018 | Louvain-la-Neuve, Belgium | 17.9 | 17.7 | 23 | 23.6 | - |
| 37 | ID 104 | Stems | *Artemisia annua* | / | / | / | / | 15/06/2018 | Louvain-la-Neuve, Belgium | 17.9 | 17.7 | 23 | 23.6 | - |
| 38 | ID 106 | Aphids | *Cirsium arvense* | *Uroleucon cirsii* | restricted | Aphidinae | Macrosiphini | 15/06/2018 | Louvain-la-Neuve, Belgium | 17.9 | 17.7 | 23 | 23.6 | - |
| 38 | ID 107 | Ants | *Cirsium arvense* | *Lasius niger* | / | Formicinae | Lasiini | 15/06/2018 | Louvain-la-Neuve, Belgium | 17.9 | 17.7 | 23 | 23.6 | - |
| 38 | ID 108 | Stems | *Cirsium arvense* | / | / | / | / | 15/06/2018 | Louvain-la-Neuve, Belgium | 17.9 | 17.7 | 23 | 23.6 | - |
| 39 | ID 110 | Aphids | *Cirsium arvense* | *Aphis fabae* | polyphagous | Aphidinae | Aphidini | 15/06/2018 | Louvain-la-Neuve, Belgium | 17.9 | 17.7 | 23 | 23.6 | - |
| 39 | ID 111 | Ants | *Cirsium arvense* | *Lasius niger* | / | Formicinae | Lasiini | 15/06/2018 | Louvain-la-Neuve, Belgium | 17.9 | 17.7 | 23 | 23.6 | - |
| 39 | ID 112 | Parasitoids | *Cirsium arvense* | *Lysiphlebus fabarum* | / | Aphidiinae | Aphidiini | 15/06/2018 | Louvain-la-Neuve, Belgium | 17.9 | 17.7 | 23 | 23.6 | - |
| 39 | ID 113 | Stems | *Cirsium arvense* | / | / | / | / | 15/06/2018 | Louvain-la-Neuve, Belgium | 17.9 | 17.7 | 23 | 23.6 | - |
| 40 | ID 115 | Aphids | *Tanacetum vulgare* | *Uroleucon achilleae* | restricted | Aphidinae | Macrosiphini | 15/06/2018 | Louvain-la-Neuve, Belgium | 17.9 | 17.7 | 23 | 23.6 | - |
| 40 | ID 116 | Ladybug larvae | *Tanacetum vulgare* | *Rhyzobius litura* | / | Coccidulinae | Coccidulini | 15/06/2018 | Louvain-la-Neuve, Belgium | 17.9 | 17.7 | 23 | 23.6 | - |
| 40 | ID 117 | Stems | *Tanacetum vulgare* | / | / | / | / | 15/06/2018 | Louvain-la-Neuve, Belgium | 17.9 | 17.7 | 23 | 23.6 | - |
| 41 | ID 119 | Aphids | *Ranunculus repens* | *Aphis fabae* | polyphagous | Aphidinae | Aphidini | 15/06/2018 | Mont St-Guibert, Belgium | 17.9 | 17.7 | 23 | 23.6 | - |
| 41 | ID 120 | Ants | *Ranunculus repens* | *Myrmica rubra* | / | Myrmicinae | Myrmicini | 15/06/2018 | Mont St-Guibert, Belgium | 17.9 | 17.7 | 23 | 23.6 | - |
| 41 | ID 121 | Stems | *Ranunculus repens* | / | / | / | / | 15/06/2018 | Mont St-Guibert, Belgium | 17.9 | 17.7 | 23 | 23.6 | + |
| 42 | ID 122 | Aphids | *Rumex obtusifolius* | *Aphis fabae* | polyphagous | Aphidinae | Aphidini | 15/06/2018 | Mont St-Guibert, Belgium | 17.9 | 17.7 | 23 | 23.6 | - |
| 42 | ID 123 | Ants | *Rumex obtusifolius* | *Lasius emarginatus* | / | Formicinae | Lasiini | 15/06/2018 | Mont St-Guibert, Belgium | 17.9 | 17.7 | 23 | 23.6 | - |
| 42 | ID 124 | Bugs | *Rumex obtusifolius* | *Coreus marginatus* | / | Coreinae | Coreini | 15/06/2018 | Mont St-Guibert, Belgium | 17.9 | 17.7 | 23 | 23.6 | - |
| 42 | ID 125 | Stems | *Rumex obtusifolius* | / | / | / | / | 15/06/2018 | Mont St-Guibert, Belgium | 17.9 | 17.7 | 23 | 23.6 | + |
| 43 | ID 127 | Aphids | *Gaillet gratteron* | *Aphis fabae* | polyphagous | Aphidinae | Aphidini | 15/06/2018 | Mont St-Guibert, Belgium | 17.9 | 17.7 | 23 | 23.6 | - |
| 43 | ID 128 | Ants | *Gaillet gratteron* | *Lasius niger* | / | Formicinae | Lasiini | 15/06/2018 | Mont St-Guibert, Belgium | 17.9 | 17.7 | 23 | 23.6 | - |
| 43 | ID 129 | Stems | *Gaillet gratteron* | / | / | / | / | 15/06/2018 | Mont St-Guibert, Belgium | 17.9 | 17.7 | 23 | 23.6 | - |
| 44 | ID 130 | Aphids | *Lactuca virosa* | *Macrosiphum euphorbiae* | polyphagous | Aphidinae | Macrosiphini | 15/06/2018 | Mont St-Guibert, Belgium | 17.9 | 17.7 | 23 | 23.6 | - |
| 44 | ID 131 | Stems | *Lactuca virosa* | / | / | / | / | 15/06/2018 | Mont St-Guibert, Belgium | 17.9 | 17.7 | 23 | 23.6 | - |
| 45 | ID 132 | Ants | *Cirsium vulgare* | *Myrmica rubra* | / | Myrmicinae | Myrmicini | 15/06/2018 | Mont St-Guibert, Belgium | 17.9 | 17.7 | 23 | 23.6 | - |
| 45 | ID 133 | Ladybugs | *Cirsium vulgare* | *Scymnus interruptus* | / | Scymninae | Scymnini | 15/06/2018 | Mont St-Guibert, Belgium | 17.9 | 17.7 | 23 | 23.6 | - |
| 45 | ID 134 | Stems | *Cirsium vulgare* | / | / | / | / | 15/06/2018 | Mont St-Guibert, Belgium | 17.9 | 17.7 | 23 | 23.6 | - |
| 45 | ID 135 * | Aphids | *Cirsium vulgare* | *Macrosiphum mordvilkoi* | specialized | Aphidinae | Macrosiphini | 15/06/2018 | Mont St-Guibert, Belgium | 17.9 | 17.7 | 23 | 23.6 | + |
| 46 | ID 136 | Aphids | *Rosa sp.* | *Aphis fabae* | polyphagous | Aphidinae | Aphidini | 15/06/2018 | Mont St-Guibert, Belgium | 17.9 | 17.7 | 23 | 23.6 | - |
| 46 | ID 137 | Stems | *Rosa sp.* | / | / | / | / | 15/06/2018 | Mont St-Guibert, Belgium | 17.9 | 17.7 | 23 | 23.6 | - |
| 47 | ID 138 | Aphids | *Cirsium vulgare* | *Aphis fabae* | polyphagous | Aphidinae | Aphidini | 16/06/2018 | Erezée, Belgium | 13.9 | 16.8 | 23 | 21.6 | + |
| 48 | ID 139 | Aphids | *Rubus sp.* | *Aphis fabae* | polyphagous | Aphidinae | Aphidini | 16/06/2018 | Erezée, Belgium | 13.9 | 16.8 | 23 | 21.6 | - |
| 49 | ID 140 | Aphids | *Daucus carota* | *Aphis fabae* | polyphagous | Aphidinae | Aphidini | 16/06/2018 | Mormont, Belgium | 13.9 | 16.8 | 23 | 21.6 | - |
| 49 | ID 141 | Ants | *Daucus carota* | *Lasius niger* | / | Formicinae | Lasiini | 16/06/2018 | Mormont, Belgium | 13.9 | 16.8 | 23 | 21.6 | - |
| 50 | ID 142 | Aphids | *Heracleum sphondylium* | *Cavariella pastinacae* | host alternating | Aphidinae | Macrosiphini | 16/06/2018 | Mormont, Belgium | 13.9 | 16.8 | 23 | 21.6 | + |
| 51 | ID 143 | Aphids | *Centaurea cyanus* | *Uroleucon jaceae* | restricted | Aphidinae | Macrosiphini | 16/06/2018 | Mormont, Belgium | 13.9 | 16.8 | 23 | 21.6 | - |
| 52 | ID 144 | Aphids | *Cirsium vulgare* | *Aphis fabae* | polyphagous | Aphidinae | Aphidini | 16/06/2018 | Mormont, Belgium | 13.9 | 16.8 | 23 | 21.6 | + |
| 52 | ID 145 | Ladybugs | *Cirsium vulgare* | *Harmonia axyridis* | / | Coccinellinae | Coccinellini | 16/06/2018 | Mormont, Belgium | 13.9 | 16.8 | 23 | 21.6 | + |
| 53 | ID 146 | Aphids | *Limoniastrum monopetalum* | *Staticobium sp.* | NA | Aphidinae | Macrosiphini | 24/06/2018 | Maspalomas, Spain | NA | NA | NA | NA | - |
| 54 | ID 147 | Aphids | *Hibiscus sabdariffa* | *Aphis gossypii* | polyphagous | Aphidinae | Aphidini | 24/06/2018 | Maspalomas, Spain | NA | NA | NA | NA | - |
| 55 | ID 148 | Aphids | *Hibiscus sabdariffa* | *Aphis gossypii* | polyphagous | Aphidinae | Aphidini | 25/06/2018 | Puerto Rico, Spain | NA | NA | NA | NA | + |
| 56 | ID 149 | Aphids | *Solanum nigrum* | *Aphis fabae* | polyphagous | Aphidinae | Aphidini | 3/07/2018 | Louvain-la-Neuve, Belgium | 23.6 | 23 | 30 | 29.7 | - |
| 56 | ID 150 | Ants | *Solanum nigrum* | *Lasius niger* | / | Formicinae | Lasiini | 3/07/2018 | Louvain-la-Neuve, Belgium | 23.6 | 23 | 30 | 29.7 | - |
| 56 | ID 152 | Stems | *Solanum nigrum* | / | / | / | / | 3/07/2018 | Louvain-la-Neuve, Belgium | 23.6 | 23 | 30 | 29.7 | - |
| 57 | ID 153 | Aphids | *Clematis vitalba* | *Aphis vitalbae* | specialized | Aphidinae | Aphidini | 3/07/2018 | Louvain-la-Neuve, Belgium | 23.6 | 23 | 30 | 29.7 | + |
| 57 | ID 154 | Ants | *Clematis vitalba* | *Lasius fuliginosus* | / | Formicinae | Lasiini | 3/07/2018 | Louvain-la-Neuve, Belgium | 23.6 | 23 | 30 | 29.7 | - |
| 57 | ID 155 | Stems | *Clematis vitalba* | / | / | / | / | 3/07/2018 | Louvain-la-Neuve, Belgium | 23.6 | 23 | 30 | 29.7 | - |
| 58 | ID 156 * | Aphids | *Hedera helix* | *Aphis fabae* | polyphagous | Aphidinae | Aphidini | 3/07/2018 | Louvain-la-Neuve, Belgium | 23.6 | 23 | 30 | 29.7 | + |
| 58 | ID 157 | Ants | *Hedera helix* | *Lasius fuliginosus* | / | Formicinae | Lasiini | 3/07/2018 | Louvain-la-Neuve, Belgium | 23.6 | 23 | 30 | 29.7 | - |
| 58 | ID 158 | Stems | *Hedera helix* | / | / | / | / | 3/07/2018 | Louvain-la-Neuve, Belgium | 23.6 | 23 | 30 | 29.7 | - |
| 59 | ID 159 | Aphids | *Plantago lanceolata* | *Dysaphis plantaginea* | host alternating | Aphidinae | Macrosiphini | 3/07/2018 | Louvain-la-Neuve, Belgium | 23.6 | 23 | 30 | 29.7 | - |
| 60 | ID 160 | Aphids | *Epipactis helleborine* | *Aphis fabae* | polyphagous | Aphidinae | Aphidini | 4/07/2018 | Louvain-la-Neuve, Belgium | 23.6 | 22.8 | 30 | 28.5 | - |
| 60 | ID 161 | Bugs | *Epipactis helleborine* | NA | / | NA | NA | 4/07/2018 | Louvain-la-Neuve, Belgium | 23.6 | 22.8 | 30 | 28.5 | - |
| 60 | ID 163 | Ants | *Epipactis helleborine* | *Lasius niger* | / | Formicinae | Lasiini | 4/07/2018 | Louvain-la-Neuve, Belgium | 23.6 | 22.8 | 30 | 28.5 | - |
| 61 | ID 164 | Aphids | *Holcus lanatus* | *Schizaphis graminum* | restricted | Aphidinae | Aphidini | 10/05/2018 | Buzin, Belgium | 13.9 | 12.9 | 18.2 | 15.9 | + |
| 62 | ID 165 | Aphids | *Centaurea jacea* | *Uroleucon jaceae* | restricted | Aphidinae | Macrosiphini | 10/05/2018 | Failon, Belgium | 13.9 | 12.9 | 18.2 | 15.9 | - |
| 62 | ID 166 | Parasitoids | *Centaurea jacea* | *Aphidius funebris* | / | Aphidiinae | Aphidiini | 10/05/2018 | Failon, Belgium | 13.9 | 12.9 | 18.2 | 15.9 | + |
| 63 | ID 167 | Aphids | *Holcus lanatus* | *Uroleucon sp.* | NA | Aphidinae | Macrosiphini | 10/05/2018 | Failon, Belgium | 13.9 | 12.9 | 18.2 | 15.9 | - |
| 64 | ID 168 | Aphids | *Triticum aestivum* | *Sitobion fragariae* | host alternating | Aphidinae | Macrosiphini | 11/05/2018 | Failon, Belgium | 13.9 | 13.2 | 18.2 | 19.9 | - |
| 65 | ID 169 | Aphids | *Triticum aestivum* | *Sitobion avenae* | polyphagous | Aphidinae | Macrosiphini | 21/05/2018 | Buzin, Belgium | 17.9 | 18.1 | 23 | 23.8 | - |
| 66 | ID 170 | Aphids | *Holcus lanatus* | *Schizaphis graminum* | restricted | Aphidinae | Aphidini | 21/05/2018 | Buzin, Belgium | 17.9 | 18.1 | 23 | 23.8 | - |
| 67 | ID 171 | Aphids | *Dactylis glomerata* | *Hyalopteroides humilis* | restricted | Aphidinae | Macrosiphini | 25/05/2018 | Buzin, Belgium | 19.6 | 19.5 | 25.7 | 25.5 | - |
| 67 | ID 172 | Parasitoids | *Dactylis glomerata* | *Aphidius rhopalosiphi* | / | Aphidiinae | Aphidiini | 25/05/2018 | Buzin, Belgium | 19.6 | 19.5 | 25.7 | 25.5 | - |
| 68 | ID 173 | Aphids | *Centaurea jacea* | *Hyalopteroides humilis* | restricted | Aphidinae | Macrosiphini | 25/05/2018 | Buzin, Belgium | 19.6 | 19.5 | 25.7 | 25.5 | - |
| 69 | ID 174 | Aphids | *Holcus lanatus* | *Schizaphis graminum* | restricted | Aphidinae | Aphidini | 25/05/2018 | Failon, Belgium | 19.6 | 19.5 | 25.7 | 25.5 | + |
| 70 | ID 175 | Aphids | *Dactylis glomerata* | *Hyalopteroides humilis* | restricted | Aphidinae | Macrosiphini | 25/05/2018 | Failon, Belgium | 19.6 | 19.5 | 25.7 | 25.5 | - |
| 71 | ID 176 | Aphids | *Triticum aestivum* | *Hyalopteroides humilis* | restricted | Aphidinae | Macrosiphini | 25/05/2018 | Failon, Belgium | 19.6 | 19.5 | 25.7 | 25.5 | + |
| 72 | ID 177 | Aphids | *Lotus corniculatus* | *Acyrthosiphon pisum* | restricted | Aphidinae | Macrosiphini | 25/05/2018 | Failon, Belgium | 19.6 | 19.5 | 25.7 | 25.5 | - |
| 73 | ID 178 | Aphids | *Centaurea jacea* | *Uroleucon jaceae* | restricted | Aphidinae | Macrosiphini | 25/05/2018 | Failon, Belgium | 19.6 | 19.5 | 25.7 | 25.5 | - |
| 74 | ID 179 | Aphids | *Rumex obtusifolius* | *Aphis rumicis* | polyphagous | Aphidinae | Aphidini | 25/05/2018 | Failon, Belgium | 19.6 | 19.5 | 25.7 | 25.5 | - |
| 75 | ID 180 | Aphids | *Triticum aestivum* | *Sitobion avenae* | polyphagous | Aphidinae | Macrosiphini | 30/05/2018 | Nodebais, Belgium | 21.5 | 21.1 | 25.7 | 25.9 | - |
| 76 | ID 181 | Aphids | *Triticum aestivum* | *Sitobion avenae* | polyphagous | Aphidinae | Macrosiphini | 30/05/2018 | Buzin, Belgium | 21.5 | 21.1 | 25.7 | 25.9 | - |
| 77 | ID 182 | Aphids | *Triticum aestivum* | *Metopolophium dirhodum* | host alternating | Aphidinae | Macrosiphini | 30/05/2018 | Buzin, Belgium | 21.5 | 21.1 | 25.7 | 25.9 | - |
| 78 | ID 183 | Aphids | *Triticum aestivum* | *Metopolophium dirhodum* | polyphagous | Aphidinae | Macrosiphini | 30/05/2018 | Failon, Belgium | 21.5 | 21.1 | 25.7 | 25.9 | - |
| 79 | ID 184 | Aphids | *Coffea arabica* | *Toxoptera aurantii* | polyphagous | Aphidinae | Aphidini | 10/07/2017 | Rwanda, Africa | NA | NA | NA | NA | - |
| 80 | ID 185 | Aphids | *Coffea arabica* | *Toxoptera aurantii* | polyphagous | Aphidinae | Aphidini | 10/07/2017 | Rwanda, Africa | NA | NA | NA | NA | - |
| 81 | ID 186 | Aphids | *Coffea arabica* | *Toxoptera aurantii* | polyphagous | Aphidinae | Aphidini | 10/07/2017 | Rwanda, Africa | NA | NA | NA | NA | - |
| 82 | ID 187 | Aphids | *Coffea arabica* | *Toxoptera aurantii* | polyphagous | Aphidinae | Aphidini | 10/07/2017 | Rwanda, Africa | NA | NA | NA | NA | - |
| 83 | ID 188 | Aphids | *Coffea arabica* | *Toxoptera aurantii* | polyphagous | Aphidinae | Aphidini | 10/07/2017 | Rwanda, Africa | NA | NA | NA | NA | + |
| 84 | ID 189 | Aphids | *Coffea arabica* | *Toxoptera aurantii* | polyphagous | Aphidinae | Aphidini | 10/07/2017 | Rwanda, Africa | NA | NA | NA | NA | + |
| 85 | ID 190 | Aphids | *Salvia farinacea* | *Aphis fabae* | polyphagous | Aphidinae | Aphidini | 7/07/2018 | Louvain-la-Neuve, Belgium | 21.5 | 22 | 26.6 | 26.8 | - |
| 85 | ID 191 | Ants | *Salvia farinacea* | *Lasius niger* | / | Formicinae | Lasiini | 7/07/2018 | Louvain-la-Neuve, Belgium | 21.5 | 22 | 26.6 | 26.8 | - |
| 85 | ID 192 | Stems | *Salvia farinacea* | / | / | / | / | 7/07/2018 | Louvain-la-Neuve, Belgium | 21.5 | 22 | 26.6 | 26.8 | - |
| 86 | ID 194 * | Aphids | *Rumex obtusifolius* | *Aphis fabae* | polyphagous | Aphidinae | Aphidini | 7/07/2018 | Louvain-la-Neuve, Belgium | 21.5 | 22 | 26.6 | 26.8 | + |
| 86 | ID 195 | Stems | *Rumex obtusifolius* | / | / | / | / | 7/07/2018 | Louvain-la-Neuve, Belgium | 21.5 | 22 | 26.6 | 26.8 | + |
| 87 | ID 197 * | Aphids | *Cirsium vulgare* | *Aphis fabae* | polyphagous | Aphidinae | Aphidini | 7/07/2018 | Louvain-la-Neuve, Belgium | 21.5 | 22 | 26.6 | 26.8 | + |
| 87 | ID 198 | Stems | *Cirsium vulgare* | / | / | / | / | 7/07/2018 | Louvain-la-Neuve, Belgium | 21.5 | 22 | 26.6 | 26.8 | - |
| 88 | ID 200 | Aphids | *Cirsium oleraceum* | *Aphis fabae* | polyphagous | Aphidinae | Aphidini | 7/07/2018 | Louvain-la-Neuve, Belgium | 21.5 | 22 | 26.6 | 26.8 | + |
| 88 | ID 201 | Ants | *Cirsium oleraceum* | *Lasius platythorax* | / | Formicinae | Lasiini | 7/07/2018 | Louvain-la-Neuve, Belgium | 21.5 | 22 | 26.6 | 26.8 | - |
| 88 | ID 202 | Stems | *Cirsium oleraceum* | / | / | / | / | 7/07/2018 | Louvain-la-Neuve, Belgium | 21.5 | 22 | 26.6 | 26.8 | - |
| 89 | ID 203 | Aphids | *Epilobium ciliatum* | *Aphis frangulae* | polyphagous | Aphidinae | Aphidini | 7/07/2018 | Louvain-la-Neuve, Belgium | 21.5 | 22 | 26.6 | 26.8 | + |
| 89 | ID 204 | Stems | *Epilobium ciliatum* | / | / | / | / | 7/07/2018 | Louvain-la-Neuve, Belgium | 21.5 | 22 | 26.6 | 26.8 | - |
| 90 | ID 206 | Aphids | *Triticum aestivum* | *Sitobion avenae* | polyphagous | Aphidinae | Macrosiphini | 6/07/2018 | Sart-Risbart, Belgium | 21.5 | 22.1 | 26.6 | 26.9 | + |
| 91 | ID 207 | Aphids | *Triticum aestivum* | *Sitobion avenae* | polyphagous | Aphidinae | Macrosiphini | 6/07/2018 | Sart-Risbart, Belgium | 21.5 | 22.1 | 26.6 | 26.9 | - |
| 92 | ID 208 | Aphids | *Achillea millefolium* | *Macrosiphoniella millefolii* | specialized | Aphidinae | Macrosiphini | 6/07/2018 | Munich, Germany | NA | NA | NA | NA | - |
| 93 | ID 209 | Aphids | *Cirsium arvense* | *Uroleucon aeneum* | restricted | Aphidinae | Macrosiphini | 9/07/2018 | Corroy-le-Grand, Belgium | 19.6 | 19.8 | 24.5 | 24.1 | - |
| 93 | ID 210 | Ladybugs | *Cirsium arvense* | *Coccinella septempunctata* | / | Coccinellinae | Coccinellini | 9/07/2018 | Corroy-le-Grand, Belgium | 19.6 | 19.8 | 24.5 | 24.1 | - |
| 93 | ID 211 | Stems | *Cirsium arvense* | / | / | / | / | 9/07/2018 | Corroy-le-Grand, Belgium | 19.6 | 19.8 | 24.5 | 24.1 | - |
| 94 | ID 213 | Aphids | *Cirsium arvense* | *Aphis fabae* | polyphagous | Aphidinae | Aphidini | 9/07/2018 | Corroy-le-Grand, Belgium | 19.6 | 19.8 | 24.5 | 24.1 | - |
| 94 | ID 214 | Aphids | *Cirsium arvense* | NA | NA | NA | NA | 9/07/2018 | Corroy-le-Grand, Belgium | 19.6 | 19.8 | 24.5 | 24.1 | - |
| 94 | ID 215 | Ants | *Cirsium arvense* | *Lasius niger* | / | Formicinae | Lasiini | 9/07/2018 | Corroy-le-Grand, Belgium | 19.6 | 19.8 | 24.5 | 24.1 | - |
| 94 | ID 216 | Stems | *Cirsium arvense* | / | / | / | / | 9/07/2018 | Corroy-le-Grand, Belgium | 19.6 | 19.8 | 24.5 | 24.1 | - |
| 95 | ID 218 | Aphids | *Cirsium arvense* | *Uroleucon aeneum* | restricted | Aphidinae | Macrosiphini | 9/07/2018 | Corroy-le-Grand, Belgium | 19.6 | 19.8 | 24.5 | 24.1 | - |
| 95 | ID 219 | Stems | *Cirsium arvense* | / | / | / | / | 9/07/2018 | Corroy-le-Grand, Belgium | 19.6 | 19.8 | 24.5 | 24.1 | - |
| 96 | ID 221 | Aphids | *Jacobaea vulgaris* | *Aphis jacobaeae* | specialized | Aphidinae | Aphidini | 9/07/2018 | Louvain-la-Neuve, Belgium | 19.6 | 19.8 | 24.5 | 24.1 | + |
| 96 | ID 222 | Ants | *Jacobaea vulgaris* | *Lasius niger* | / | Formicinae | Lasiini | 9/07/2018 | Louvain-la-Neuve, Belgium | 19.6 | 19.8 | 24.5 | 24.1 | - |
| 96 | ID 223 | Stems | *Jacobaea vulgaris* | / | / | / | / | 9/07/2018 | Louvain-la-Neuve, Belgium | 19.6 | 19.8 | 24.5 | 24.1 | - |
| 97 | ID 225 | Aphids | *Tanacetum sp.* | *Uroleucon achilleae* | restricted | Aphidinae | Macrosiphini | 9/07/2018 | Louvain-la-Neuve, Belgium | 19.6 | 19.8 | 24.5 | 24.1 | - |
| 97 | ID 226 | Leaves | *Tanacetum sp.* | / | / | / | / | 9/07/2018 | Louvain-la-Neuve, Belgium | 19.6 | 19.8 | 24.5 | 24.1 | - |
| 98 | ID 227 | Aphids | *Chaenomeles japonica* | *Aphis fabae* | polyphagous | Aphidinae | Aphidini | 9/07/2018 | Louvain-la-Neuve, Belgium | 19.6 | 19.8 | 24.5 | 24.1 | - |
| 98 | ID 228 | Ants | *Chaenomeles japonica* | *Lasius niger* | / | Formicinae | Lasiini | 9/07/2018 | Louvain-la-Neuve, Belgium | 19.6 | 19.8 | 24.5 | 24.1 | - |
| 98 | ID 229 | Stems | *Chaenomeles japonica* | / | / | / | / | 9/07/2018 | Louvain-la-Neuve, Belgium | 19.6 | 19.8 | 24.5 | 24.1 | - |
| 99 | ID 231 | Aphids | *Spiraea japonica* | *Aphis fabae* | polyphagous | Aphidinae | Aphidini | 9/07/2018 | Louvain-la-Neuve, Belgium | 19.6 | 19.8 | 24.5 | 24.1 | - |
| 99 | ID 232 | Aphids | *Spiraea japonica* | *Aphis spiraecola* | polyphagous | Aphidinae | Aphidini | 9/07/2018 | Louvain-la-Neuve, Belgium | 19.6 | 19.8 | 24.5 | 24.1 | - |
| 99 | ID 233 | Ants | *Spiraea japonica* | *Lasius niger* | / | Formicinae | Lasiini | 9/07/2018 | Louvain-la-Neuve, Belgium | 19.6 | 19.8 | 24.5 | 24.1 | - |
| 99 | ID 234 | Stems | *Spiraea japonica* | / | / | / | / | 9/07/2018 | Louvain-la-Neuve, Belgium | 19.6 | 19.8 | 24.5 | 24.1 | - |
| 100 | ID 236 | Aphids | *Cirsium arvense* | *Aphis fabae* | polyphagous | Aphidinae | Aphidini | 9/07/2018 | Louvain-la-Neuve, Belgium | 19.6 | 19.8 | 24.5 | 24.1 | - |
| 100 | ID 237 | Ants | *Cirsium arvense* | *Lasius niger* | / | Formicinae | Lasiini | 9/07/2018 | Louvain-la-Neuve, Belgium | 19.6 | 19.8 | 24.5 | 24.1 | - |
| 100 | ID 238 | Stems | *Cirsium arvense* | / | / | / | / | 9/07/2018 | Louvain-la-Neuve, Belgium | 19.6 | 19.8 | 24.5 | 24.1 | - |
| 101 | ID 240 | Ants | *Jacobaea vulgaris* | *Lasius niger* | / | Formicinae | Lasiini | 16/07/2018 | Maransart, Belgium | 23.6 | 22.8 | 30 | 29.9 | - |
| 101 | ID 241 | Aphids | *Jacobaea vulgaris* | *Aphis jacobaeae* | specialized | Aphidinae | Aphidini | 16/07/2018 | Maransart, Belgium | 23.6 | 22.8 | 30 | 29.9 | - |
| 101 | ID 242 | Stems | *Jacobaea vulgaris* | / | / | / | / | 16/07/2018 | Maransart, Belgium | 23.6 | 22.8 | 30 | 29.9 | - |
| 102 | ID 244 | Ants | *Jacobaea vulgaris* | NA | / | NA | NA | 16/07/2018 | Maransart, Belgium | 23.6 | 22.8 | 30 | 29.9 | - |
| 102 | ID 245 | Aphids | *Jacobaea vulgaris* | *Aphis cacaliasteris* | polyphagous | Aphidinae | Aphidini | 16/07/2018 | Maransart, Belgium | 23.6 | 22.8 | 30 | 29.9 | + |
| 102 | ID 246 | Aphid Midge Larvae | *Jacobaea vulgaris* | *Aphidoletes aphidimyza* | / | Cecidomyiinae | Aphidoletini | 16/07/2018 | Maransart, Belgium | 23.6 | 22.8 | 30 | 29.9 | + |
| 102 | ID 247 | Stems | *Jacobaea vulgaris* | / | / | / | / | 16/07/2018 | Maransart, Belgium | 23.6 | 22.8 | 30 | 29.9 | - |
| 103 | ID 249 | Aphids | *Vesca sp.* | *Aphis craccae* | specialized | Aphidinae | Aphidini | 16/07/2018 | Maransart, Belgium | 23.6 | 22.8 | 30 | 29.9 | - |
| 103 | ID 250 | Stems | *Vesca sp.* | / | / | / | / | 16/07/2018 | Maransart, Belgium | 23.6 | 22.8 | 30 | 29.9 | - |
| 104 | ID 252 | Ants | *Jacobaea vulgaris* | *Lasius niger* | / | Formicinae | Lasiini | 16/07/2018 | Maransart, Belgium | 23.6 | 22.8 | 30 | 29.9 | - |
| 104 | ID 253 | Aphids | *Jacobaea vulgaris* | *Aphis jacobaeae* | specialized | Aphidinae | Aphidini | 16/07/2018 | Maransart, Belgium | 23.6 | 22.8 | 30 | 29.9 | - |
| 104 | ID 254 | Stems | *Jacobaea vulgaris* | / | / | / | / | 16/07/2018 | Maransart, Belgium | 23.6 | 22.8 | 30 | 29.9 | - |
| 105 | ID 256 | Ants | *Jacobaea vulgaris* | NA | / | NA | NA | 16/07/2018 | Lasne, Belgium | 23.6 | 22.8 | 30 | 29.9 | - |
| 105 | ID 257 | Aphids | *Jacobaea vulgaris* | *Brachycaudus cardui* | restricted | Aphidinae | Macrosiphini | 16/07/2018 | Lasne, Belgium | 23.6 | 22.8 | 30 | 29.9 | - |
| 105 | ID 258 | Stems | *Jacobaea vulgaris* | / | / | / | / | 16/07/2018 | Lasne, Belgium | 23.6 | 22.8 | 30 | 29.9 | - |
| 106 | ID 260 | Aphids | *Cirsium arvense* | *Uroleucon cirsii* | restricted | Aphidinae | Macrosiphini | 16/07/2018 | Lasne, Belgium | 23.6 | 22.8 | 30 | 29.9 | - |
| 106 | ID 261 | Stems | *Cirsium arvense* | / | / | / | / | 16/07/2018 | Lasne, Belgium | 23.6 | 22.8 | 30 | 29.9 | - |
| 107 | ID 263 * | Aphids | *Rosa sp.* | *Macrosiphum rosae* | restricted | Aphidinae | Macrosiphini | 16/07/2018 | Lasne, Belgium | 23.6 | 22.8 | 30 | 29.9 | + |
| 107 | ID 264 | Stems | *Rosa sp.* | / | / | / | / | 16/07/2018 | Lasne, Belgium | 23.6 | 22.8 | 30 | 29.9 | - |
| 108 | ID 265 | Ants | *Tanacetum vulgare* | *Lasius niger* | / | Formicinae | Lasiini | 16/07/2018 | Lasne, Belgium | 23.6 | 22.8 | 30 | 29.9 | - |
| 108 | ID 266 | Aphids | *Tanacetum vulgare* | *Metopeurum fuscoviride* | restricted | Aphidinae | Macrosiphini | 16/07/2018 | Lasne, Belgium | 23.6 | 22.8 | 30 | 29.9 | - |
| 108 | ID 267 | Stems | *Tanacetum vulgare* | / | / | / | / | 16/07/2018 | Lasne, Belgium | 23.6 | 22.8 | 30 | 29.9 | - |
| 109 | ID 269 | Aphids | *Centaurea jacea* | *Uroleucon jaceae* | restricted | Aphidinae | Macrosiphini | 16/07/2018 | Lasne, Belgium | 23.6 | 22.8 | 30 | 29.9 | - |
| 109 | ID 270 | Stems | *Centaurea jacea* | / | / | / | / | 16/07/2018 | Lasne, Belgium | 23.6 | 22.8 | 30 | 29.9 | - |
| 110 | ID 272 | Aphids | *Centaurea jacea* | *Uroleucon jaceae* | restricted | Aphidinae | Macrosiphini | 16/07/2018 | Lasne, Belgium | 23.6 | 22.8 | 30 | 29.9 | - |
| 110 | ID 273 | Stems | *Centaurea jacea* | / | / | / | / | 16/07/2018 | Lasne, Belgium | 23.6 | 22.8 | 30 | 29.9 | - |
| 111 | ID 275 | Aphids | *Artemisia vulgaris* | *Macrosiphoniella artemisiae* | specialized | Aphidinae | Macrosiphini | 17/07/2018 | Vieux-Sart, Belgium | 20.4 | 20.5 | 25.7 | 25.8 | - |
| 111 | ID 276 | Stems | *Artemisia vulgaris* | / | / | / | / | 17/07/2018 | Vieux-Sart, Belgium | 20.4 | 20.5 | 25.7 | 25.8 | - |
| 112 | ID 278 | Ants | *Chenopodium album* | NA | / | NA | NA | 17/07/2018 | Vieux-Sart, Belgium | 20.4 | 20.5 | 25.7 | 25.8 | - |
| 112 | ID 279 | Aphids | *Chenopodium album* | NA | NA | NA | NA | 17/07/2018 | Vieux-Sart, Belgium | 20.4 | 20.5 | 25.7 | 25.8 | - |
| 112 | ID 280 | Stems | *Chenopodium album* | / | / | / | / | 17/07/2018 | Vieux-Sart, Belgium | 20.4 | 20.5 | 25.7 | 25.8 | - |
| 113 | ID 282 | Ants | *Epipactis helleborine* | *Myrmica rubra* | / | Myrmicinae | Myrmicini | 17/07/2018 | Vieux-Sart, Belgium | 20.4 | 20.5 | 25.7 | 25.8 | + |
| 113 | ID 283 | Aphids | *Epipactis helleborine* | *Aphis fabae* | polyphagous | Aphidinae | Aphidini | 17/07/2018 | Vieux-Sart, Belgium | 20.4 | 20.5 | 25.7 | 25.8 | - |
| 113 | ID 284 | Stems | *Epipactis helleborine* | / | / | / | / | 17/07/2018 | Vieux-Sart, Belgium | 20.4 | 20.5 | 25.7 | 25.8 | - |
| 114 | ID 286 | Aphids | *Artemisia vulgaris* | *Macrosiphoniella artemisiae* | specialized | Aphidinae | Macrosiphini | 17/07/2018 | Vieux-Sart, Belgium | 20.4 | 20.5 | 25.7 | 25.8 | - |
| 114 | ID 287 | Aphid Midge Larvae | *Artemisia vulgaris* | *Aphidoletes aphidimyza* | / | Cecidomyiinae | Aphidoletini | 17/07/2018 | Vieux-Sart, Belgium | 20.4 | 20.5 | 25.7 | 25.8 | - |
| 114 | ID 288 | Stems | *Artemisia vulgaris* | / | / | / | / | 17/07/2018 | Vieux-Sart, Belgium | 20.4 | 20.5 | 25.7 | 25.8 | - |
| 115 | ID 290 | Aphids | *Cirsium vulgare* | *Aphis fabae* | polyphagous | Aphidinae | Aphidini | 17/07/2018 | Vieux-Sart, Belgium | 20.4 | 20.5 | 25.7 | 25.8 | - |
| 115 | ID 291 | Parasitoids | *Cirsium vulgare* | *Lysiphlebus fabarum* | / | Aphidiinae | Aphidiini | 17/07/2018 | Vieux-Sart, Belgium | 20.4 | 20.5 | 25.7 | 25.8 | - |
| 115 | ID 292 | Ladybug Larvae | *Cirsium vulgare* | *Harmonia axyridis* | / | Coccinellinae | Coccinellini | 17/07/2018 | Vieux-Sart, Belgium | 20.4 | 20.5 | 25.7 | 25.8 | - |
| 115 | ID 293 | Stems | *Cirsium vulgare* | / | / | / | / | 17/07/2018 | Vieux-Sart, Belgium | 20.4 | 20.5 | 25.7 | 25.8 | - |
| 116 | ID 294 | Aphids | *Phragmites australis* | *Hyalopterus pruni* | host alternating | Aphidinae | Aphidini | 17/07/2018 | Vieux-Sart, Belgium | 20.4 | 20.5 | 25.7 | 25.8 | - |
| 116 | ID 295 | Ladybug Larvae | *Phragmites australis* | *Propylea quatuordecimpunctata* | / | Coccinellinae | Coccinellini | 17/07/2018 | Vieux-Sart, Belgium | 20.4 | 20.5 | 25.7 | 25.8 | - |
| 116 | ID 296 | Ladybug Larvae | *Phragmites australis* | *Harmonia axyridis* | / | Coccinellinae | Coccinellini | 17/07/2018 | Vieux-Sart, Belgium | 20.4 | 20.5 | 25.7 | 25.8 | - |
| 116 | ID 297 | Leaves | *Phragmites australis* | / | / | / | / | 17/07/2018 | Vieux-Sart, Belgium | 20.4 | 20.5 | 25.7 | 25.8 | - |
| 117 | ID 298 | Aphids | *Sonchus asper* | *Uroleucon sonchi* | restricted | Aphidinae | Macrosiphini | 17/07/2018 | Corbais, Belgium | 20.4 | 20.5 | 25.7 | 25.8 | - |
| 117 | ID 299 | Ladybugs | *Sonchus asper* | *Coccinella septempunctata* | / | Coccinellinae | Coccinellini | 17/07/2018 | Corbais, Belgium | 20.4 | 20.5 | 25.7 | 25.8 | - |
| 117 | ID 300 | Stems | *Sonchus asper* | / | / | / | / | 17/07/2018 | Corbais, Belgium | 20.4 | 20.5 | 25.7 | 25.8 | - |
| 118 | ID 302 | Aphids | *Daucus carota* | *Aphis fabae* | polyphagous | Aphidinae | Aphidini | 17/07/2018 | Corroy-le-Grand, Belgium | 20.4 | 20.5 | 25.7 | 25.8 | - |
| 118 | ID 303 | Ladybugs | *Daucus carota* | *Coccinella septempunctata* | / | Coccinellinae | Coccinellini | 17/07/2018 | Corroy-le-Grand, Belgium | 20.4 | 20.5 | 25.7 | 25.8 | - |
| 118 | ID 304 | Hoverfly Larvae | *Daucus carota* | *Scaeva pyrastri* | / | Syrphinae | Syrphini | 17/07/2018 | Corroy-le-Grand, Belgium | 20.4 | 20.5 | 25.7 | 25.8 | - |
| 118 | ID 305 | Stems | *Daucus carota* | / | / | / | / | 17/07/2018 | Corroy-le-Grand, Belgium | 20.4 | 20.5 | 25.7 | 25.8 | - |
| 119 | ID 306 | Aphids | *Populus tremula* | *Chaitophorus populeti* | restricted | Chaitophorinae | Chaitophorini | 17/07/2018 | Corroy-le-Grand, Belgium | 20.4 | 20.5 | 25.7 | 25.8 | - |
| 119 | ID 307 | Ants | *Populus tremula* | *Lasius niger* | / | Formicinae | Lasiini | 17/07/2018 | Corroy-le-Grand, Belgium | 20.4 | 20.5 | 25.7 | 25.8 | - |
| 119 | ID 308 | Stems | *Populus tremula* | / | / | / | / | 17/07/2018 | Corroy-le-Grand, Belgium | 20.4 | 20.5 | 25.7 | 25.8 | - |
| 120 | ID 310 | Aphids | *Daucus carota* | *Aphis fabae* | polyphagous | Aphidinae | Aphidini | 17/07/2018 | Corroy-le-Grand, Belgium | 20.4 | 20.5 | 25.7 | 25.8 | - |
| 120 | ID 311 | Ladybugs | *Daucus carota* | [*Coccinella septempunctata*](https://blast.ncbi.nlm.nih.gov/Blast.cgi#alnHdr_1214171732) | / | Coccinellinae | Coccinellini | 17/07/2018 | Corroy-le-Grand, Belgium | 20.4 | 20.5 | 25.7 | 25.8 | - |
| 120 | ID 312 | Lacewing Larvae | *Daucus carota* | *Chrysoperla nipponensis* | / | Chrysopinae | Chrysopini | 17/07/2018 | Corroy-le-Grand, Belgium | 20.4 | 20.5 | 25.7 | 25.8 | - |
| 120 | ID 313 | Hoverfly Larvae | *Daucus carota* | *Episyrphus balteatus* | / | Syrphinae | Syrphini | 17/07/2018 | Corroy-le-Grand, Belgium | 20.4 | 20.5 | 25.7 | 25.8 | - |
| 120 | ID 314 | Stems | *Daucus carota* | / | / | / | / | 17/07/2018 | Corroy-le-Grand, Belgium | 20.4 | 20.5 | 25.7 | 25.8 | - |
| 121 | ID 316 | Ants | *Epilobium angustifolium* | *Lasius niger* | / | Formicinae | Lasiini | 17/07/2018 | Corroy-le-Grand, Belgium | 20.4 | 20.5 | 25.7 | 25.8 | - |
| 121 | ID 317 | Aphids | *Epilobium angustifolium* | *Aphis salicariae* | host alternating | Aphidinae | Aphidini | 17/07/2018 | Corroy-le-Grand, Belgium | 20.4 | 20.5 | 25.7 | 25.8 | - |
| 121 | ID 318 | Stems | *Epilobium angustifolium* | / | / | / | / | 17/07/2018 | Corroy-le-Grand, Belgium | 20.4 | 20.5 | 25.7 | 25.8 | - |
| 122 | ID 319 | Aphids | *Artemisia vulgaris* | *Macrosiphoniella artemisiae* | specialized | Aphidinae | Macrosiphini | 17/07/2018 | Corroy-le-Grand, Belgium | 20.4 | 20.5 | 25.7 | 25.8 | - |
| 122 | ID 320 | Stems | *Artemisia vulgaris* | / | / | / | / | 17/07/2018 | Corroy-le-Grand, Belgium | 20.4 | 20.5 | 25.7 | 25.8 | - |
| 123 | ID 321 | Aphids | *Phragmites australis* | *Hyalopterus pruni* | host alternating | Aphidinae | Aphidini | 18/07/2018 | Gistoux, Belgium | 19.6 | 19.6 | 25.7 | 25.4 | - |
| 123 | ID 322 | Hoverfly Larvae | *Phragmites australis* | *Syrphus ribesii* | / | Syrphinae | Syrphini | 18/07/2018 | Gistoux, Belgium | 19.6 | 19.6 | 25.7 | 25.4 | - |
| 123 | ID 323 | Leaves | *Phragmites australis* | / | / | / | / | 18/07/2018 | Gistoux, Belgium | 19.6 | 19.6 | 25.7 | 25.4 | - |
| 124 | ID 324 | Ants | *Daucus carota* | *Myrmica rubra* | / | Myrmicinae | Myrmicini | 18/07/2018 | Gistoux, Belgium | 19.6 | 19.6 | 25.7 | 25.4 | - |
| 124 | ID 325 | Aphids | *Daucus carota* | *Aphis fabae* | polyphagous | Aphidinae | Aphidini | 18/07/2018 | Gistoux, Belgium | 19.6 | 19.6 | 25.7 | 25.4 | - |
| 124 | ID 326 | Stems | *Daucus carota* | / | / | / | / | 18/07/2018 | Gistoux, Belgium | 19.6 | 19.6 | 25.7 | 25.4 | - |
| 125 | ID 328 | Aphids | *Conyza canadensis* | *Uroleucon erigeronensis* | restricted | Aphidinae | Macrosiphini | 18/07/2018 | Gistoux, Belgium | 19.6 | 19.6 | 25.7 | 25.4 | - |
| 125 | ID 329 | Hoverfly Larvae | *Conyza canadensis* | *Syrphus ribesii* | / | Syrphinae | Syrphini | 18/07/2018 | Gistoux, Belgium | 19.6 | 19.6 | 25.7 | 25.4 | - |
| 125 | ID 330 | Parasitoids | *Conyza canadensis* | *Tetrastichinae sp.* | / | Tetrastichinae | NA | 18/07/2018 | Gistoux, Belgium | 19.6 | 19.6 | 25.7 | 25.4 | - |
| 125 | ID 331 | Stems | *Conyza canadensis* | / | / | / | / | 18/07/2018 | Gistoux, Belgium | 19.6 | 19.6 | 25.7 | 25.4 | - |
| 126 | ID 332 | Aphids | *Sonchus asper* | *Uroleucon sonchi* | restricted | Aphidinae | Macrosiphini | 18/07/2018 | Gistoux, Belgium | 19.6 | 19.6 | 25.7 | 25.4 | - |
| 126 | ID 333 | Parasitoids | *Sonchus asper* | *Aphelinus abdominalis* | / | Aphelininae | Aphelinini | 18/07/2018 | Gistoux, Belgium | 19.6 | 19.6 | 25.7 | 25.4 | - |
| 126 | ID 334 | Stems | *Sonchus asper* | / | / | / | / | 18/07/2018 | Gistoux, Belgium | 19.6 | 19.6 | 25.7 | 25.4 | - |
| 127 | ID 335 | Ants | *Cirsium arvense* | *Myrmica rubra* | / | Myrmicinae | Myrmicini | 18/07/2018 | Gistoux, Belgium | 19.6 | 19.6 | 25.7 | 25.4 | - |
| 127 | ID 336 | Aphids | *Cirsium arvense* | *Aphis fabae* | polyphagous | Aphidinae | Aphidini | 18/07/2018 | Gistoux, Belgium | 19.6 | 19.6 | 25.7 | 25.4 | - |
| 127 | ID 337 | Bugs | *Cirsium arvense* | *Dolycoris baccarum* | / | Pentatominae | Carpocorini | 18/07/2018 | Gistoux, Belgium | 19.6 | 19.6 | 25.7 | 25.4 | - |
| 127 | ID 338 | Stems | *Cirsium arvense* | / | / | / | / | 18/07/2018 | Gistoux, Belgium | 19.6 | 19.6 | 25.7 | 25.4 | - |
| 128 | ID 340 | Aphids | *Sonchus asper* | *Uroleucon sonchi* | restricted | Aphidinae | Macrosiphini | 18/07/2018 | Gistoux, Belgium | 19.6 | 19.6 | 25.7 | 25.4 | - |
| 128 | ID 341 | Stems | *Sonchus asper* | / | / | / | / | 18/07/2018 | Gistoux, Belgium | 19.6 | 19.6 | 25.7 | 25.4 | - |
| 129 | ID 342 | Aphids | *Epilobium ciliatum* | *Aphis grossulariae* | restricted | Aphidinae | Aphidini | 18/07/2018 | Gistoux, Belgium | 19.6 | 19.6 | 25.7 | 25.4 | - |
| 129 | ID 343 | Hoverfly Larvae | *Epilobium ciliatum* | *Platycheirus scutatus* | / | Syrphinae | Bacchini | 18/07/2018 | Gistoux, Belgium | 19.6 | 19.6 | 25.7 | 25.4 | - |
| 129 | ID 344 | Hoverfly Larvae | *Epilobium ciliatum* | *Platycheirus scutatus* | / | Syrphinae | Bacchini | 18/07/2018 | Gistoux, Belgium | 19.6 | 19.6 | 25.7 | 25.4 | - |
| 129 | ID 345 | Stems | *Epilobium ciliatum* | / | / | / | / | 18/07/2018 | Gistoux, Belgium | 19.6 | 19.6 | 25.7 | 25.4 | - |
| 130 | ID 347 | Aphids | *Cirsium arvense* | *Uroleucon cirsii* | restricted | Aphidinae | Macrosiphini | 18/07/2018 | Gistoux, Belgium | 19.6 | 19.6 | 25.7 | 25.4 | - |
| 130 | ID 348 | Ladybugs | *Cirsium arvense* | *Coccinella septempunctata* | / | Coccinellinae | Coccinellini | 18/07/2018 | Gistoux, Belgium | 19.6 | 19.6 | 25.7 | 25.4 | - |
| 130 | ID 349 | Hoverfly Larvae | *Cirsium arvense* | *Scaeva pyrastri* | / | Syrphinae | Syrphini | 18/07/2018 | Gistoux, Belgium | 19.6 | 19.6 | 25.7 | 25.4 | - |
| 130 | ID 350 | Stems | *Cirsium arvense* | / | / | / | / | 18/07/2018 | Gistoux, Belgium | 19.6 | 19.6 | 25.7 | 25.4 | - |
| 131 | ID 352 | Aphids | *Daucus carota* | *Aphis fabae* | polyphagous | Aphidinae | Aphidini | 18/07/2018 | Gistoux, Belgium | 19.6 | 19.6 | 25.7 | 25.4 | - |
| 131 | ID 353 | Moth Larvae | *Daucus carota* | *Eupithecia tripunctaria* | / | Larentiinae | Eupitheciini | 18/07/2018 | Gistoux, Belgium | 19.6 | 19.6 | 25.7 | 25.4 | - |
| 131 | ID 354 | Stems | *Daucus carota* | / | / | / | / | 18/07/2018 | Gistoux, Belgium | 19.6 | 19.6 | 25.7 | 25.4 | - |
| 132 | ID 356 | Aphids | *Phragmites australis* | *Hyalopterus pruni* | host alternating | Aphidinae | Aphidini | 18/07/2018 | Gistoux, Belgium | 19.6 | 19.6 | 25.7 | 25.4 | - |
| 132 | ID 357 | Hoverfly Larvae | *Phragmites australis* | *Syrphus ribesii* | / | Syrphinae | Syrphini | 18/07/2018 | Gistoux, Belgium | 19.6 | 19.6 | 25.7 | 25.4 | - |
| 132 | ID 358 | Leaves | *Phragmites australis* | / | / | / | / | 18/07/2018 | Gistoux, Belgium | 19.6 | 19.6 | 25.7 | 25.4 | - |
| 133 | ID 359 | Aphids | *Epilobium hirsutum* | *Aphis epilobii* | restricted | Aphidinae | Aphidini | 18/07/2018 | Gistoux, Belgium | 19.6 | 19.6 | 25.7 | 25.4 | - |
| 133 | ID 360 | Moth Larvae | *Epilobium hirsutum* | *Mompha epilobiella* | / | NA | NA | 18/07/2018 | Gistoux, Belgium | 19.6 | 19.6 | 25.7 | 25.4 | - |
| 133 | ID 361 | Stems | *Epilobium hirsutum* | / | / | / | / | 18/07/2018 | Gistoux, Belgium | 19.6 | 19.6 | 25.7 | 25.4 | - |
| 134 | ID 363 | Aphids | *Sonchus asper* | *Uroleucon sonchi* | restricted | Aphidinae | Macrosiphini | 19/07/2018 | Limal, Belgium | 21.5 | 21 | 26.6 | 27 | - |
| 134 | ID 364 | Stems | *Sonchus asper* | / | / | / | / | 19/07/2018 | Limal, Belgium | 21.5 | 21 | 26.6 | 27 | - |
| 135 | ID 365 | Aphids | *Lactuca serriola* | *Acyrthosiphon lactucae* | restricted | Aphidinae | Macrosiphini | 19/07/2018 | Limal, Belgium | 21.5 | 21 | 26.6 | 27 | - |
| 135 | ID 366 | Stems | *Lactuca serriola* | / | / | / | / | 19/07/2018 | Limal, Belgium | 21.5 | 21 | 26.6 | 27 | - |
| 136 | ID 367 | Ants | *Laphangium luteoalbum* | *Lasius niger* | / | Formicinae | Lasiini | 19/07/2018 | Limal, Belgium | 21.5 | 21 | 26.6 | 27 | - |
| 136 | ID 368 | Aphids | *Laphangium luteoalbum* | *Brachycaudus cardui* | restricted | Aphidinae | Macrosiphini | 19/07/2018 | Limal, Belgium | 21.5 | 21 | 26.6 | 27 | + |
| 136 | ID 369 * | Aphids | *Laphangium luteoalbum* | *Aphis fabae* | polyphagous | Aphidinae | Aphidini | 19/07/2018 | Limal, Belgium | 21.5 | 21 | 26.6 | 27 | + |
| 136 | ID 370 | Stems | *Laphangium luteoalbum* | / | / | / | / | 19/07/2018 | Limal, Belgium | 21.5 | 21 | 26.6 | 27 | - |
| 137 | ID 371 | Ants | *Laphangium luteoalbum* | *Formica lemani* | / | Formicinae | Formicini | 19/07/2018 | Limal, Belgium | 21.5 | 21 | 26.6 | 27 | - |
| 137 | ID 372 | Aphids | *Laphangium luteoalbum* | *Aphis fabae* | polyphagous | Aphidinae | Aphidini | 19/07/2018 | Limal, Belgium | 21.5 | 21 | 26.6 | 27 | - |
| 137 | ID 373 | Stems | *Laphangium luteoalbum* | / | / | / | / | 19/07/2018 | Limal, Belgium | 21.5 | 21 | 26.6 | 27 | - |
| 138 | ID 374 | Ants | *Sonchus asper* | *Lasius niger* | / | Formicinae | Lasiini | 19/07/2018 | Limal, Belgium | 21.5 | 21 | 26.6 | 27 | - |
| 138 | ID 375 | Aphids | *Sonchus asper* | *Aphis fabae* | polyphagous | Aphidinae | Aphidini | 19/07/2018 | Limal, Belgium | 21.5 | 21 | 26.6 | 27 | - |
| 138 | ID 376 | Stems | *Sonchus asper* | / | / | / | / | 19/07/2018 | Limal, Belgium | 21.5 | 21 | 26.6 | 27 | - |
| 139 | ID 377 | Ants | *Solanum chenopodioides* | *Lasius niger* | / | Formicinae | Lasiini | 19/07/2018 | Limal, Belgium | 21.5 | 21 | 26.6 | 27 | + |
| 139 | ID 378 * | Aphids | *Solanum chenopodioides* | *Aphis fabae* | polyphagous | Aphidinae | Aphidini | 19/07/2018 | Limal, Belgium | 21.5 | 21 | 26.6 | 27 | + |
| 139 | ID 379 | Stems | *Solanum chenopodioides* | / | / | / | / | 19/07/2018 | Limal, Belgium | 21.5 | 21 | 26.6 | 27 | + |
| 140 | ID 380 * | Aphids | *Daucus carota* | *Aphis fabae* | polyphagous | Aphidinae | Aphidini | 19/07/2018 | Limal, Belgium | 21.5 | 21 | 26.6 | 27 | + |
| 140 | ID 381 | Moth Larvae | *Daucus carota* | *Eupithecia trisignaria* | / | Larentiinae | Eupitheciini | 19/07/2018 | Limal, Belgium | 21.5 | 21 | 26.6 | 27 | + |
| 140 | ID 382 | Hoverfly Larvae | *Daucus carota* | *Episyrphus balteatus* | / | Syrphinae | Syrphini | 19/07/2018 | Limal, Belgium | 21.5 | 21 | 26.6 | 27 | + |
| 140 | ID 383 | Stems | *Daucus carota* | / | / | / | / | 19/07/2018 | Limal, Belgium | 21.5 | 21 | 26.6 | 27 | - |
| 141 | ID 385 | Ants | *Rumex obtusifolius* | *Lasius niger* | / | Formicinae | Lasiini | 19/07/2018 | Limal, Belgium | 21.5 | 21 | 26.6 | 27 | - |
| 141 | ID 386 | Aphids | *Rumex obtusifolius* | *Aphis fabae* | polyphagous | Aphidinae | Aphidini | 19/07/2018 | Limal, Belgium | 21.5 | 21 | 26.6 | 27 | + |
| 141 | ID 387 | Stems | *Rumex obtusifolius* | / | / | / | / | 19/07/2018 | Limal, Belgium | 21.5 | 21 | 26.6 | 27 | - |
| 142 | ID 389 | Aphids | *Sonchus asper* | *Hyperomyzus lactucae* | host alternating | Aphidinae | Macrosiphini | 19/07/2018 | Limal, Belgium | 21.5 | 21 | 26.6 | 27 | - |
| 142 | ID 390 | Hoverfly Larvae | *Sonchus asper* | *Paragus cooverti* | / | Syrphinae | Paragini | 19/07/2018 | Limal, Belgium | 21.5 | 21 | 26.6 | 27 | - |
| 142 | ID 391 | Stems | *Sonchus asper* | / | / | / | / | 19/07/2018 | Limal, Belgium | 21.5 | 21 | 26.6 | 27 | - |
| 143 | ID 392 | Aphids | *Sonchus asper* | *Uroleucon sonchi* | restricted | Aphidinae | Macrosiphini | 19/07/2018 | Limal, Belgium | 21.5 | 21 | 26.6 | 27 | - |
| 143 | ID 393 | Stems | *Sonchus asper* | / | / | / | / | 19/07/2018 | Limal, Belgium | 21.5 | 21 | 26.6 | 27 | - |
| 144 | ID 394 | Aphids | *Glebionis segetum* | *Aphis fabae* | polyphagous | Aphidinae | Aphidini | 19/07/2018 | Limal, Belgium | 21.5 | 21 | 26.6 | 27 | - |
| 144 | ID 395 | Stems | *Glebionis segetum* | / | / | / | / | 19/07/2018 | Limal, Belgium | 21.5 | 21 | 26.6 | 27 | - |
| 145 | ID 396 | Aphids | *Robinia pseudoacacia* | *Aphis glycines* | host alternating | Aphidinae | Aphidini | 19/07/2018 | Limal, Belgium | 21.5 | 21 | 26.6 | 27 | - |
| 145 | ID 397 | Stems | *Robinia pseudoacacia* | / | / | / | / | 19/07/2018 | Limal, Belgium | 21.5 | 21 | 26.6 | 27 | - |
| 146 | ID 399 | Ants | *Chenopodium album* | *Lasius niger* | / | Formicinae | Lasiini | 19/07/2018 | Limal, Belgium | 21.5 | 21 | 26.6 | 27 | - |
| 146 | ID 400 | Aphids | *Chenopodium album* | *Aphis fabae* | polyphagous | Aphidinae | Aphidini | 19/07/2018 | Limal, Belgium | 21.5 | 21 | 26.6 | 27 | - |
| 146 | ID 401 | Parasitoids | *Chenopodium album* | *Lysiphlebus fabarum* | / | Aphidiinae | Aphidiini | 19/07/2018 | Limal, Belgium | 21.5 | 21 | 26.6 | 27 | - |
| 146 | ID 402 | Stems | *Chenopodium album* | / | / | / | / | 19/07/2018 | Limal, Belgium | 21.5 | 21 | 26.6 | 27 | - |
| 147 | ID 403 | Ants | *Cirsium arvense* | *Lasius niger* | / | Formicinae | Lasiini | 19/07/2018 | Limal, Belgium | 21.5 | 21 | 26.6 | 27 | - |
| 147 | ID 404 | Aphids | *Cirsium arvense* | *Aphis fabae* | polyphagous | Aphidinae | Aphidini | 19/07/2018 | Limal, Belgium | 21.5 | 21 | 26.6 | 27 | - |
| 147 | ID 405 | Stems | *Cirsium arvense* | / | / | / | / | 19/07/2018 | Limal, Belgium | 21.5 | 21 | 26.6 | 27 | - |
| 148 | ID 406 | Ants | *Sonchus asper* | *Lasius niger* | / | Formicinae | Lasiini | 19/07/2018 | Limal, Belgium | 21.5 | 21 | 26.6 | 27 | - |
| 148 | ID 407 | Aphids | *Sonchus asper* | *Aphis fabae* | polyphagous | Aphidinae | Aphidini | 19/07/2018 | Limal, Belgium | 21.5 | 21 | 26.6 | 27 | - |
| 148 | ID 408 | Stems | *Sonchus asper* | / | / | / | / | 19/07/2018 | Limal, Belgium | 21.5 | 21 | 26.6 | 27 | - |
| 149 | ID 409 | Aphids | *Sonchus asper* | *Uroleucon sonchi* | restricted | Aphidinae | Macrosiphini | 19/07/2018 | Limal, Belgium | 21.5 | 21 | 26.6 | 27 | - |
| 149 | ID 410 | Hoverfly Larvae | *Sonchus asper* | *Scaeva pyrastri* | / | Syrphinae | Syrphini | 19/07/2018 | Limal, Belgium | 21.5 | 21 | 26.6 | 27 | - |
| 149 | ID 411 | Stems | *Sonchus asper* | / | / | / | / | 19/07/2018 | Limal, Belgium | 21.5 | 21 | 26.6 | 27 | - |
| 150 | ID 412 | Aphids | *Sonchus asper* | *Uroleucon sonchi* | restricted | Aphidinae | Macrosiphini | 19/07/2018 | Limal, Belgium | 21.5 | 21 | 26.6 | 27 | - |
| 150 | ID 413 | Hoverfly Larvae | *Sonchus asper* | *Scaeva pyrastri* | / | Syrphinae | Syrphini | 19/07/2018 | Limal, Belgium | 21.5 | 21 | 26.6 | 27 | - |
| 150 | ID 414 | Stems | *Sonchus asper* | / | / | / | / | 19/07/2018 | Limal, Belgium | 21.5 | 21 | 26.6 | 27 | - |
| 151 | ID 415 | Aphids | *Sonchus asper* | *Hyperomyzus lactucae* | host alternating | Aphidinae | Macrosiphini | 20/07/2018 | Grez-Doiceau, Belgium | 20.4 | 20.6 | 26.6 | 26 | - |
| 151 | ID 416 | Hoverfly Larvae | *Sonchus asper* | *Episyrphus balteatus* | / | Syrphinae | Syrphini | 20/07/2018 | Grez-Doiceau, Belgium | 20.4 | 20.6 | 26.6 | 26 | - |
| 151 | ID 417 | Stems | *Sonchus asper* | / | / | / | / | 20/07/2018 | Grez-Doiceau, Belgium | 20.4 | 20.6 | 26.6 | 26 | - |
| 152 | ID 418 | Aphids | *Daucus carota* | *Aphis fabae* | polyphagous | Aphidinae | Aphidini | 20/07/2018 | Grez-Doiceau, Belgium | 20.4 | 20.6 | 26.6 | 26 | - |
| 152 | ID 419 | Hoverfly Larvae | *Daucus carota* | *Episyrphus balteatus* | / | Syrphinae | Syrphini | 20/07/2018 | Grez-Doiceau, Belgium | 20.4 | 20.6 | 26.6 | 26 | - |
| 152 | ID 420 | Stems | *Daucus carota* | / | / | / | / | 20/07/2018 | Grez-Doiceau, Belgium | 20.4 | 20.6 | 26.6 | 26 | - |
| 153 | ID 421 | Ants | *Cirsium arvense* | *Lasius niger* | / | Formicinae | Lasiini | 20/07/2018 | Grez-Doiceau, Belgium | 20.4 | 20.6 | 26.6 | 26 | - |
| 153 | ID 422 | Aphids | *Cirsium arvense* | *Aphis fabae* | polyphagous | Aphidinae | Aphidini | 20/07/2018 | Grez-Doiceau, Belgium | 20.4 | 20.6 | 26.6 | 26 | + |
| 153 | ID 423 | Stems | *Cirsium arvense* | / | / | / | / | 20/07/2018 | Grez-Doiceau, Belgium | 20.4 | 20.6 | 26.6 | 26 | + |
| 154 | ID 424 | Aphids | *Daucus carota* | *Aphis fabae* | polyphagous | Aphidinae | Aphidini | 20/07/2018 | Grez-Doiceau, Belgium | 20.4 | 20.6 | 26.6 | 26 | + |
| 154 | ID 425 | Stems | *Daucus carota* | / | / | / | / | 20/07/2018 | Grez-Doiceau, Belgium | 20.4 | 20.6 | 26.6 | 26 | - |
| 155 | ID 426 | Aphids | *Epilobium hirsutum* | *Aphis frangulae* | polyphagous | Aphidinae | Aphidini | 20/07/2018 | Grez-Doiceau, Belgium | 20.4 | 20.6 | 26.6 | 26 | - |
| 155 | ID 427 | Ladybugs | *Epilobium hirsutum* | *Coccinella septempunctata* | / | Coccinellinae | Coccinellini | 20/07/2018 | Grez-Doiceau, Belgium | 20.4 | 20.6 | 26.6 | 26 | - |
| 155 | ID 428 * | Hoverfly Larvae | *Epilobium hirsutum* | *Scaeva pyrastri* | / | Syrphinae | Syrphini | 20/07/2018 | Grez-Doiceau, Belgium | 20.4 | 20.6 | 26.6 | 26 | + |
| 155 | ID 429 | Stems | *Epilobium hirsutum* | / | / | / | / | 20/07/2018 | Grez-Doiceau, Belgium | 20.4 | 20.6 | 26.6 | 26 | - |
| 156 | ID 431 | Aphids | *Epilobium hirsutum* | *Aphis epilobii* | restricted | Aphidinae | Aphidini | 20/07/2018 | Grez-Doiceau, Belgium | 20.4 | 20.6 | 26.6 | 26 | - |
| 156 | ID 432 | Parasitoids | *Epilobium hirsutum* | *Eulophinae sp.* | / | Eulophinae | NA | 20/07/2018 | Grez-Doiceau, Belgium | 20.4 | 20.6 | 26.6 | 26 | - |
| 156 | ID 433 | Stems | *Epilobium hirsutum* | / | / | / | / | 20/07/2018 | Grez-Doiceau, Belgium | 20.4 | 20.6 | 26.6 | 26 | - |
| 157 | ID 434 | Aphids | *Sonchus asper* | *Uroleucon sonchi* | restricted | Aphidinae | Macrosiphini | 20/07/2018 | Grez-Doiceau, Belgium | 20.4 | 20.6 | 26.6 | 26 | - |
| 157 | ID 435 | Aphid Midge Larvae | *Sonchus asper* | *Aphidoletes aphidimyza* | / | Cecidomyiinae | Aphidoletini | 20/07/2018 | Grez-Doiceau, Belgium | 20.4 | 20.6 | 26.6 | 26 | - |
| 157 | ID 436 | Hoverfly Larvae | *Sonchus asper* | *Scaeva pyrastri* | / | Syrphinae | Syrphini | 20/07/2018 | Grez-Doiceau, Belgium | 20.4 | 20.6 | 26.6 | 26 | - |
| 157 | ID 437 | Stems | *Sonchus asper* | / | / | / | / | 20/07/2018 | Grez-Doiceau, Belgium | 20.4 | 20.6 | 26.6 | 26 | + |
| 158 | ID 438 | Aphids | *Sambucus nigra* | *Aphis sambuci* | host alternating | Aphidinae | Aphidini | 20/07/2018 | Grez-Doiceau, Belgium | 20.4 | 20.6 | 26.6 | 26 | + |
| 158 | ID 439 | Stems | *Sambucus nigra* | / | / | / | / | 20/07/2018 | Grez-Doiceau, Belgium | 20.4 | 20.6 | 26.6 | 26 | + |
| 159 | ID 440 | Hoverfly Larvae | *Sonchus asper* | [*Paragus cooverti*](https://blast.ncbi.nlm.nih.gov/Blast.cgi#alnHdr_1389462909) | / | Syrphinae | Paragini | 20/07/2018 | Grez-Doiceau, Belgium | 20.4 | 20.6 | 26.6 | 26 | - |
| 159 | ID 441 | Stems | *Sonchus asper* | / | / | / | / | 20/07/2018 | Grez-Doiceau, Belgium | 20.4 | 20.6 | 26.6 | 26 | - |
| 159 | ID 443 | Aphids | *Sonchus asper* | *Uroleucon sonchi* | restricted | Aphidinae | Macrosiphini | 20/07/2018 | Grez-Doiceau, Belgium | 20.4 | 20.6 | 26.6 | 26 | - |
| 160 | ID 444 | Stems | *Epilobium hirsutum* | / | / | / | / | 20/07/2018 | Grez-Doiceau, Belgium | 20.4 | 20.6 | 26.6 | 26 | - |
| 160 | ID 445 | Aphids | *Epilobium hirsutum* | *Aphis epilobii* | restricted | Aphidinae | Aphidini | 21/07/2018 | Grez-Doiceau, Belgium | 21.5 | 21.4 | 25.7 | 25.8 | - |
| 161 | ID 446 | Aphids | *Robinia pseudoacacia* | *Aphis glycines* | host alternating | Aphidinae | Aphidini | 30/07/2018 | Louvain-la-neuve, Belgium | 23.6 | 23.7 | 30 | 28.8 | - |
| 161 | ID 447 | Bugs | *Robinia pseudoacacia* | *Pinalitus viscicola* | / | Mirinae | Mirini | 30/07/2018 | Louvain-la-neuve, Belgium | 23.6 | 23.7 | 30 | 28.8 | - |
| 161 | ID 448 | Ladybugs | *Robinia pseudoacacia* | *Harmonia axyridis* | / | Coccinellinae | Coccinellini | 30/07/2018 | Louvain-la-neuve, Belgium | 23.6 | 23.7 | 30 | 28.8 | - |
| 161 | ID 449 | Ladybug Larvae | *Robinia pseudoacacia* | *Harmonia axyridis* | / | Coccinellinae | Coccinellini | 30/07/2018 | Louvain-la-neuve, Belgium | 23.6 | 23.7 | 30 | 28.8 | - |
| 161 | ID 450 | Aphids | *Robinia pseudoacacia* | *Tinocallis viridis* | NA | Calaphidinae | Panaphidini | 30/07/2018 | Louvain-la-neuve, Belgium | 23.6 | 23.7 | 30 | 28.8 | - |
| 161 | ID 451 | Stems | *Robinia pseudoacacia* | / | / | / | / | 30/07/2018 | Louvain-la-neuve, Belgium | 23.6 | 23.7 | 30 | 28.8 | - |
| 162 | ID 452 | Aphids | *Cirsium arvense* | *Aphis fabae* | polyphagous | Aphidinae | Aphidini | 31/07/2018 | Braine-l'Alleud, Belgium | 23.6 | 22.7 | 26.6 | 26.8 | - |
| 162 | ID 453 | Ants | *Cirsium arvense* | *Lasius niger* | / | Formicinae | Lasiini | 31/07/2018 | Braine-l'Alleud, Belgium | 23.6 | 22.7 | 26.6 | 26.8 | - |
| 162 | ID 454 | Stems | *Cirsium arvense* | / | / | / | / | 31/07/2018 | Braine-l'Alleud, Belgium | 23.6 | 22.7 | 26.6 | 26.8 | - |
| 163 | ID 456 | Aphids | *Corylus avellana* | *Pterocallis maculata* | specialized | Calaphidinae | Panaphidini | 31/07/2018 | Braine-l'Alleud, Belgium | 23.6 | 22.7 | 26.6 | 26.8 | - |
| 163 | ID 457 | Ants | *Corylus avellana* | *Lasius niger* | / | Formicinae | Lasiini | 31/07/2018 | Braine-l'Alleud, Belgium | 23.6 | 22.7 | 26.6 | 26.8 | - |
| 163 | ID 458 | Leaves | *Corylus avellana* | / | / | / | / | 31/07/2018 | Braine-l'Alleud, Belgium | 23.6 | 22.7 | 26.6 | 26.8 | - |
| 164 | ID 459 | Aphids | *Epilobium hirsutum* | *Aphis epilobii* | restricted | Aphidinae | Aphidini | 31/07/2018 | Braine-l'Alleud, Belgium | 23.6 | 22.7 | 26.6 | 26.8 | - |
| 164 | ID 460 | Parasitoids | *Epilobium hirsutum* | *Pnigalio sp.* | / | Eulophinae | Eulophini | 31/07/2018 | Braine-l'Alleud, Belgium | 23.6 | 22.7 | 26.6 | 26.8 | - |
| 164 | ID 461 | Bugs | *Epilobium hirsutum* | *Dicyphus bolivari* | / | Bryocorinae | Dicyphini | 31/07/2018 | Braine-l'Alleud, Belgium | 23.6 | 22.7 | 26.6 | 26.8 | - |
| 164 | ID 462 | Stems | *Epilobium hirsutum* | / | / | / | / | 31/07/2018 | Braine-l'Alleud, Belgium | 23.6 | 22.7 | 26.6 | 26.8 | - |
| 165 | ID 463 | Aphids | *Daucus carota* | *Cavariella theobaldi* | host alternating | Aphidinae | Macrosiphini | 31/07/2018 | Braine-l'Alleud, Belgium | 23.6 | 22.7 | 26.6 | 26.8 | - |
| 165 | ID 464 | Ladybug Larvae | *Daucus carota* | *Hippodamia variegata* | / | Coccinellinae | Coccinellini | 31/07/2018 | Braine-l'Alleud, Belgium | 23.6 | 22.7 | 26.6 | 26.8 | - |
| 165 | ID 465 | Aphids | *Daucus carota* | *Aphis fabae* | polyphagous | Aphidinae | Aphidini | 31/07/2018 | Braine-l'Alleud, Belgium | 23.6 | 22.7 | 26.6 | 26.8 | - |
| 165 | ID 466 | Moth Larvae | *Daucus carota* | *Eupithecia tripunctaria* | / | Larentiinae | Eupitheciini | 31/07/2018 | Braine-l'Alleud, Belgium | 23.6 | 22.7 | 26.6 | 26.8 | - |
| 165 | ID 467 | Stems | *Daucus carota* | / | / | / | / | 31/07/2018 | Braine-l'Alleud, Belgium | 23.6 | 22.7 | 26.6 | 26.8 | - |
| 166 | ID 469 | Bugs | *Corylus avellana* | *Malacocoris chlorizans* | / | Orthotylinae | Orthotylini | 31/07/2018 | Genval, Belgium | 23.6 | 22.7 | 26.6 | 26.8 | - |
| 166 | ID 470 | Aphids | *Corylus avellana* | *Corylobium avellanae* | specialized | Aphidinae | Macrosiphini | 31/07/2018 | Genval, Belgium | 23.6 | 22.7 | 26.6 | 26.8 | - |
| 166 | ID 471 | Bugs | *Corylus avellana* | *Deraeocoris lutescens* | / | Deraeocorinae | Deraeocorini | 31/07/2018 | Genval, Belgium | 23.6 | 22.7 | 26.6 | 26.8 | - |
| 166 | ID 472 | Leaves | *Corylus avellana* | / | / | / | / | 31/07/2018 | Genval, Belgium | 23.6 | 22.7 | 26.6 | 26.8 | - |
| 167 | ID 473 | Aphids | *Rumex obtusifolius* | *Aphis fabae* | polyphagous | Aphidinae | Aphidini | 31/07/2018 | Genval, Belgium | 23.6 | 22.7 | 26.6 | 26.8 | - |
| 167 | ID 474 | Ants | *Rumex obtusifolius* | *Lasius niger* | / | Formicinae | Lasiini | 31/07/2018 | Genval, Belgium | 23.6 | 22.7 | 26.6 | 26.8 | - |
| 167 | ID 475 | Parasitoids | *Rumex obtusifolius* | *Lysiphlebus fabarum* | / | Aphidiinae | Aphidiini | 31/07/2018 | Genval, Belgium | 23.6 | 22.7 | 26.6 | 26.8 | + |
| 167 | ID 476 | Leaves | *Rumex obtusifolius* | / | / | / | / | 31/07/2018 | Genval, Belgium | 23.6 | 22.7 | 26.6 | 26.8 | - |
| 168 | ID 477 | Bugs | *Symphytum tuberrosum* | *Dictyla humuli* | / | Tinginae | Tingini | 31/07/2018 | Genval, Belgium | 23.6 | 22.7 | 26.6 | 26.8 | - |
| 168 | ID 478 | Bugs | *Symphytum tuberrosum* | *Dictyla humuli* | / | Tinginae | Tingini | 31/07/2018 | Genval, Belgium | 23.6 | 22.7 | 26.6 | 26.8 | + |
| 168 | ID 479 | Leaves | *Symphytum tuberrosum* | / | / | / | / | 31/07/2018 | Genval, Belgium | 23.6 | 22.7 | 26.6 | 26.8 | - |
| 169 | ID 480 | Aphids | *Eupatorium cannabinum* | *Aphis frangulae* | polyphagous | Aphidinae | Aphidini | 31/07/2018 | Genval, Belgium | 23.6 | 22.7 | 26.6 | 26.8 | + |
| 169 | ID 481 | Ants | *Eupatorium cannabinum* | *Lasius niger* | / | Formicinae | Lasiini | 31/07/2018 | Genval, Belgium | 23.6 | 22.7 | 26.6 | 26.8 | - |
| 169 | ID 482 | Stems | *Eupatorium cannabinum* | / | / | / | / | 31/07/2018 | Genval, Belgium | 23.6 | 22.7 | 26.6 | 26.8 | - |
| 169 | ID 483 | Leaves | *Eupatorium cannabinum* | / | / | / | / | 31/07/2018 | Genval, Belgium | 23.6 | 22.7 | 26.6 | 26.8 | - |
| 170 | ID 485 * | Aphids | *Cirsium oleraceum* | *Aphis fabae* | polyphagous | Aphidinae | Aphidini | 31/07/2018 | Genval, Belgium | 23.6 | 22.7 | 26.6 | 26.8 | + |
| 170 | ID 486 | Ants | *Cirsium oleraceum* | *Lasius niger* | / | Formicinae | Lasiini | 31/07/2018 | Genval, Belgium | 23.6 | 22.7 | 26.6 | 26.8 | + |
| 170 | ID 487 | Stems | *Cirsium oleraceum* | / | / | / | / | 31/07/2018 | Genval, Belgium | 23.6 | 22.7 | 26.6 | 26.8 | + |
| 170 | ID 489 | Fly Larvae | *Cirsium oleraceum* | *Chamaemyiidae sp.* | / | NA | NA | 31/07/2018 | Genval, Belgium | 23.6 | 22.7 | 26.6 | 26.8 | + |
| 171 | ID 490 | Ants | *Ribes odoratum* | *Lasius niger* | / | Formicinae | Lasiini | 25/06/2018 | Lautenne, Belgium | 17.9 | 17.8 | 23 | 23.1 | - |
| 171 | ID 491 * | Aphids | *Ribes odoratum* | *Aphis grossulariae* | restricted | Aphidinae | Aphidini | 25/06/2018 | Lautenne, Belgium | 17.9 | 17.8 | 23 | 23.1 | + |
| 172 | ID 492 | Ants | *Spiraea sp.* | *Lasius niger* | / | Formicinae | Lasiini | 25/06/2018 | Lautenne, Belgium | 17.9 | 17.8 | 23 | 23.1 | - |
| 172 | ID 493 | Aphids | *Spiraea sp.* | *Aphis spiraecola* | polyphagous | Aphidinae | Aphidini | 25/06/2018 | Lautenne, Belgium | 17.9 | 17.8 | 23 | 23.1 | - |
| 173 | ID 494 | Ants | *Rubus idaeus* | *Lasius niger* | / | Formicinae | Lasiini | 25/06/2018 | Lautenne, Belgium | 17.9 | 17.8 | 23 | 23.1 | - |
| 173 | ID 495 | Aphids | *Rubus idaeus* | *Aphis idaei* | restricted | Aphidinae | Aphidini | 25/06/2018 | Lautenne, Belgium | 17.9 | 17.8 | 23 | 23.1 | - |
| 174 | ID 496 | Ants | *Spiraea salicifolia* | *Lasius niger* | / | Formicinae | Lasiini | 25/06/2018 | Lautenne, Belgium | 17.9 | 17.8 | 23 | 23.1 | - |
| 174 | ID 497 | Aphids | *Spiraea salicifolia* | *Aphis spiraecola* | polyphagous | Aphidinae | Aphidini | 25/06/2018 | Lautenne, Belgium | 17.9 | 17.8 | 23 | 23.1 | - |
| 175 | ID 498 | Aphids | *Crepis capillaris* | *Uroleucon hypochoeridis* | restricted | Aphidinae | Macrosiphini | 5/07/2018 | Soignies, Belgium | 21.5 | 21.1 | 25.7 | 25.8 | - |
| 175 | ID 499 | Aphids | *Crepis capillaris* | *Uroleucon hypochoeridis* | restricted | Aphidinae | Macrosiphini | 5/07/2018 | Soignies, Belgium | 21.5 | 21.1 | 25.7 | 25.8 | - |
| 175 | ID 500 | Moth Larvae | *Crepis capillaris* | *Hecatera bicolorata* | / | Hadeninae | NA | 5/07/2018 | Soignies, Belgium | 21.5 | 21.1 | 25.7 | 25.8 | - |
| 176 | ID 501 | Ants | *Cirsium vulgare* | *Lasius niger* | / | Formicinae | Lasiini | 5/07/2018 | Soignies, Belgium | 21.5 | 21.1 | 25.7 | 25.8 | - |
| 176 | ID 502 | Aphids | *Cirsium vulgare* | *Aphis fabae* | polyphagous | Aphidinae | Aphidini | 5/07/2018 | Soignies, Belgium | 21.5 | 21.1 | 25.7 | 25.8 | - |
| 177 | ID 503 | Ants | *Daucus carota* | *Lasius niger* | / | Formicinae | Lasiini | 5/07/2018 | Soignies, Belgium | 21.5 | 21.1 | 25.7 | 25.8 | - |
| 177 | ID 504 * | Aphids | *Daucus carota* | *Aphis fabae* | polyphagous | Aphidinae | Aphidini | 5/07/2018 | Soignies, Belgium | 21.5 | 21.1 | 25.7 | 25.8 | + |
| 178 | ID 505 | Ants | *Cornus mas* | *Lasius niger* | / | Formicinae | Lasiini | 5/07/2018 | Soignies, Belgium | 21.5 | 21.1 | 25.7 | 25.8 | - |
| 178 | ID 506 | Parasitoids | *Cornus mas* | *Lysiphlebus fabarum* | / | Aphidiinae | Aphidiini | 5/07/2018 | Soignies, Belgium | 21.5 | 21.1 | 25.7 | 25.8 | - |
| 178 | ID 507 | Aphids | *Cornus mas* | *Aphis fabae* | polyphagous | Aphidinae | Aphidini | 5/07/2018 | Soignies, Belgium | 21.5 | 21.1 | 25.7 | 25.8 | - |
| 179 | ID 508 | Ants | *Crataegus monogyna* | *Lasius niger* | / | Formicinae | Lasiini | 5/07/2018 | Soignies, Belgium | 21.5 | 21.1 | 25.7 | 25.8 | + |
| 179 | ID 509 * | Aphids | *Crataegus monogyna* | *Aphis pomi* | restricted | Aphidinae | Aphidini | 5/07/2018 | Soignies, Belgium | 21.5 | 21.1 | 25.7 | 25.8 | + |
| 180 | ID 510 | Ants | *Crataegus monogyna* | *Lasius niger* | / | Formicinae | Lasiini | 5/07/2018 | Soignies, Belgium | 21.5 | 21.1 | 25.7 | 25.8 | - |
| 180 | ID 511 * | Aphids | *Crataegus monogyna* | *Aphis pomi* | restricted | Aphidinae | Aphidini | 5/07/2018 | Soignies, Belgium | 21.5 | 21.1 | 25.7 | 25.8 | + |
| 181 | ID 512 | Ants | *Cornus mas* | *Lasius niger* | / | Formicinae | Lasiini | 5/07/2018 | Soignies, Belgium | 21.5 | 21.1 | 25.7 | 25.8 | + |
| 181 | ID 513 | Aphids | *Cornus mas* | *Aphis fabae* | polyphagous | Aphidinae | Aphidini | 5/07/2018 | Soignies, Belgium | 21.5 | 21.1 | 25.7 | 25.8 | + |
| 182 | ID 514 | Ants | *Sonchus asper* | *Lasius niger* | / | Formicinae | Lasiini | 5/07/2018 | Soignies, Belgium | 21.5 | 21.1 | 25.7 | 25.8 | - |
| 182 | ID 515 | Aphids | *Sonchus asper* | *Aphis fabae* | polyphagous | Aphidinae | Aphidini | 5/07/2018 | Soignies, Belgium | 21.5 | 21.1 | 25.7 | 25.8 | + |
| 183 | ID 516 | Ants | *Rumex obtusifolius* | *Lasius niger* | / | Formicinae | Lasiini | 5/07/2018 | Soignies, Belgium | 21.5 | 21.1 | 25.7 | 25.8 | + |
| 183 | ID 517 * | Aphids | *Rumex obtusifolius* | *Aphis fabae* | polyphagous | Aphidinae | Aphidini | 5/07/2018 | Soignies, Belgium | 21.5 | 21.1 | 25.7 | 25.8 | + |
| 183 | ID 518 | Bugs | *Rumex obtusifolius* | *Coreus marginatus* | / | Coreinae | Coreini | 5/07/2018 | Soignies, Belgium | 21.5 | 21.1 | 25.7 | 25.8 | + |
| 184 | ID 519 | Aphids | *Calendula officinalis* | *Aphis fabae* | polyphagous | Aphidinae | Aphidini | 12/07/2018 | Lautenne, Belgium | 19.6 | 19.5 | 24.5 | 24.4 | - |
| 185 | ID 520 | Aphids | *Rosa sp.* | *Macrosiphum rosae* | restricted | Aphidinae | Macrosiphini | 12/07/2018 | Lautenne, Belgium | 19.6 | 19.5 | 24.5 | 24.4 | + |
| 185 | ID 521 | Hoverfly Larvae | *Rosa sp.* | *Syrphus ribesii* | / | Syrphinae | Syrphini | 12/07/2018 | Lautenne, Belgium | 19.6 | 19.5 | 24.5 | 24.4 | - |
| 186 | ID 522 | Aphids | *Tropaeolum majus* | *Aphis fabae* | polyphagous | Aphidinae | Aphidini | 12/07/2018 | Lautenne, Belgium | 19.6 | 19.5 | 24.5 | 24.4 | - |
| 186 | ID 523 | Parasitoids | *Tropaeolum majus* | *Diglyphus isaea* | / | Eulophinae | NA | 12/07/2018 | Lautenne, Belgium | 19.6 | 19.5 | 24.5 | 24.4 | - |
| 186 | ID 524 | Moth Larvae | *Tropaeolum majus* | *Pieris brassicae* | / | Pierinae | Pierini | 12/07/2018 | Lautenne, Belgium | 19.6 | 19.5 | 24.5 | 24.4 | - |
| 187 | ID 525 | Ants | *Yucca filamentosa* | *Lasius niger* | / | Formicinae | Lasiini | 12/07/2018 | Lautenne, Belgium | 19.6 | 19.5 | 24.5 | 24.4 | + |
| 187 | ID 526 | Aphids | *Yucca filamentosa* | *Uroleucon sonchi* | restricted | Aphidinae | Macrosiphini | 12/07/2018 | Lautenne, Belgium | 19.6 | 19.5 | 24.5 | 24.4 | - |
| 188 | ID 527 | Aphids | *Sonchus asper* | *Uroleucon sonchi* | restricted | Aphidinae | Macrosiphini | 3/08/2018 | Villers-la-ville, Belgium | 23.6 | 25.8 | 30 | 32.9 | - |
| 188 | ID 528 | Ladybugs | *Sonchus asper* | *Propylea quatuordecimpunctata* | / | Coccinellinae | Coccinellini | 3/08/2018 | Villers-la-ville, Belgium | 23.6 | 25.8 | 30 | 32.9 | - |
| 188 | ID 529 | Stems | *Sonchus asper* | / | / | / | / | 3/08/2018 | Villers-la-ville, Belgium | 23.6 | 25.8 | 30 | 32.9 | - |
| 189 | ID 531 | Aphids | *Sonchus asper* | *Uroleucon sonchi* | restricted | Aphidinae | Macrosiphini | 3/08/2018 | Villers-la-ville, Belgium | 23.6 | 25.8 | 30 | 32.9 | - |
| 189 | ID 532 | Ladybugs | *Sonchus asper* | *Coccinella septempunctata* | / | Coccinellinae | Coccinellini | 3/08/2018 | Villers-la-ville, Belgium | 23.6 | 25.8 | 30 | 32.9 | - |
| 189 | ID 533 | Stems | *Sonchus asper* | / | / | / | / | 3/08/2018 | Villers-la-ville, Belgium | 23.6 | 25.8 | 30 | 32.9 | - |
| 190 | ID 535 | Aphids | *Sonchus asper* | *Uroleucon sonchi* | restricted | Aphidinae | Macrosiphini | 3/08/2018 | Villers-la-ville, Belgium | 23.6 | 25.8 | 30 | 32.9 | - |
| 190 | ID 536 | Ladybugs | *Sonchus asper* | *Coccinella septempunctata* | / | Coccinellinae | Coccinellini | 3/08/2018 | Villers-la-ville, Belgium | 23.6 | 25.8 | 30 | 32.9 | + |
| 190 | ID 537 | Aphid Midge Larvae | *Sonchus asper* | *Aphidoletes aphidimyza* | / | Cecidomyiinae | Aphidoletini | 3/08/2018 | Villers-la-ville, Belgium | 23.6 | 25.8 | 30 | 32.9 | + |
| 190 | ID 538 | Hoverfly Larvae | *Sonchus asper* | *Scaeva pyrastri* | / | Syrphinae | Syrphini | 3/08/2018 | Villers-la-ville, Belgium | 23.6 | 25.8 | 30 | 32.9 | + |
| 190 | ID 539 | Stems | *Sonchus asper* | / | / | / | / | 3/08/2018 | Villers-la-ville, Belgium | 23.6 | 25.8 | 30 | 32.9 | - |
| 191 | ID 541 | Aphids | *Epilobium hirsutum* | *Aphis salicariae* | host alternating | Aphidinae | Aphidini | 3/08/2018 | Villers-la-ville, Belgium | 23.6 | 25.8 | 30 | 32.9 | + |
| 191 | ID 542 | Stems | *Epilobium hirsutum* | / | / | / | / | 3/08/2018 | Villers-la-ville, Belgium | 23.6 | 25.8 | 30 | 32.9 | - |
| 192 | ID 544 | Aphids | *Cirsium arvense* | *Aphis fabae* | polyphagous | Aphidinae | Aphidini | 3/08/2018 | Villers-la-ville, Belgium | 23.6 | 25.8 | 30 | 32.9 | + |
| 192 | ID 545 | Ants | *Cirsium arvense* | *Lasius niger* | / | Formicinae | Lasiini | 3/08/2018 | Villers-la-ville, Belgium | 23.6 | 25.8 | 30 | 32.9 | - |
| 192 | ID 546 | Stems | *Cirsium arvense* | / | / | / | / | 3/08/2018 | Villers-la-ville, Belgium | 23.6 | 25.8 | 30 | 32.9 | + |
| 193 | ID 548 | Aphids | *Papaver somniferum* | *Aphis pomi* | restricted | Aphidinae | Aphidini | 3/08/2018 | Villers-la-ville, Belgium | 23.6 | 25.8 | 30 | 32.9 | + |
| 194 | ID 549 | Aphids | *Malus domestica* | *Aphis pomi* | restricted | Aphidinae | Aphidini | 3/08/2018 | Villers-la-ville, Belgium | 23.6 | 25.8 | 30 | 32.9 | - |
| 194 | ID 550 | Ants | *Malus domestica* | Lasius niger | / | Formicinae | Lasiini | 3/08/2018 | Villers-la-ville, Belgium | 23.6 | 25.8 | 30 | 32.9 | - |
| 194 | ID 551 | Leaves | *Malus domestica* | / | / | / | / | 3/08/2018 | Villers-la-ville, Belgium | 23.6 | 25.8 | 30 | 32.9 | + |
| 195 | ID 553 | Aphids | *Tanacetum vulgare* | *Metopeurum fuscoviride* | restricted | Aphidinae | Macrosiphini | 3/08/2018 | Villers-la-ville, Belgium | 23.6 | 25.8 | 30 | 32.9 | - |
| 195 | ID 554 | Ants | *Tanacetum vulgare* | *Lasius niger* | / | Formicinae | Lasiini | 3/08/2018 | Villers-la-ville, Belgium | 23.6 | 25.8 | 30 | 32.9 | - |
| 195 | ID 555 | Parasitoids | *Tanacetum vulgare* | *Lysiphlebus hirticornis* | / | Aphidiinae | Aphidiini | 3/08/2018 | Villers-la-ville, Belgium | 23.6 | 25.8 | 30 | 32.9 | + |
| 195 | ID 556 | Stems | *Tanacetum vulgare* | / | / | / | / | 3/08/2018 | Villers-la-ville, Belgium | 23.6 | 25.8 | 30 | 32.9 | + |
| 196 | ID 558 * | Aphids | *Leucanthemum vulgare* | *Aphis fabae* | polyphagous | Aphidinae | Aphidini | 3/08/2018 | Villers-la-ville, Belgium | 23.6 | 25.8 | 30 | 32.9 | + |
| 196 | ID 559 | Ants | *Leucanthemum vulgare* | *Lasius niger* | / | Formicinae | Lasiini | 3/08/2018 | Villers-la-ville, Belgium | 23.6 | 25.8 | 30 | 32.9 | - |
| 196 | ID 560 | Stems | *Leucanthemum vulgare* | / | / | / | / | 3/08/2018 | Villers-la-ville, Belgium | 23.6 | 25.8 | 30 | 32.9 | - |
| 197 | ID 561 | Aphids | *Sonchus asper* | *Hyperomyzus lactucae* | host alternating | Aphidinae | Macrosiphini | 16/08/2018 | Ernage, Belgium | 20.4 | 20.5 | 26.6 | 26.5 | - |
| 197 | ID 562 | Bugs | *Sonchus asper* | *Dolycoris baccarum* | / | Pentatominae | Carpocorini | 16/08/2018 | Ernage, Belgium | 20.4 | 20.5 | 26.6 | 26.5 | - |
| 197 | ID 563 | Hoverfly Larvae | *Sonchus asper* | *Paragus cooverti* | / | Syrphinae | Paragini | 16/08/2018 | Ernage, Belgium | 20.4 | 20.5 | 26.6 | 26.5 | - |
| 197 | ID 564 | Hoverfly Larvae | *Sonchus asper* | *Paragus cooverti* | / | Syrphinae | Paragini | 16/08/2018 | Ernage, Belgium | 20.4 | 20.5 | 26.6 | 26.5 | - |
| 197 | ID 565 | Stems | *Sonchus asper* | / | / | / | / | 16/08/2018 | Ernage, Belgium | 20.4 | 20.5 | 26.6 | 26.5 | - |
| 198 | ID 567 | Aphids | *Heracleum sphondylium* | *Cavariella theobaldi* | host alternating | Aphidinae | Macrosiphini | 16/08/2018 | Ernage, Belgium | 20.4 | 20.5 | 26.6 | 26.5 | - |
| 198 | ID 568 | Bugs | *Heracleum sphondylium* | *Dolycoris baccarum* | / | Pentatominae | Carpocorini | 16/08/2018 | Ernage, Belgium | 20.4 | 20.5 | 26.6 | 26.5 | - |
| 198 | ID 569 | Ladybugs | *Heracleum sphondylium* | *Hippodamia variegata* | / | Coccinellinae | Coccinellini | 16/08/2018 | Ernage, Belgium | 20.4 | 20.5 | 26.6 | 26.5 | - |
| 198 | ID 570 | Parasitoids | *Heracleum sphondylium* | *Binodoxys heraclei* | / | Aphidiinae | Aphidiini | 16/08/2018 | Ernage, Belgium | 20.4 | 20.5 | 26.6 | 26.5 | - |
| 198 | ID 571 | Hoverfly Larvae | *Heracleum sphondylium* | *Scaeva pyrastri* | / | Syrphinae | Syrphini | 16/08/2018 | Ernage, Belgium | 20.4 | 20.5 | 26.6 | 26.5 | - |
| 198 | ID 572 | Aphid Midge Larvae | *Heracleum sphondylium* | *Aphidoletes aphidimyza* | / | Cecidomyiinae | Aphidoletini | 16/08/2018 | Ernage, Belgium | 20.4 | 20.5 | 26.6 | 26.5 | - |
| 198 | ID 573 | Stems | *Heracleum sphondylium* | / | / | / | / | 16/08/2018 | Ernage, Belgium | 20.4 | 20.5 | 26.6 | 26.5 | + |
| 199 | ID 575 | Aphids | *Heracleum sphondylium* | *Aphis fabae* | polyphagous | Aphidinae | Aphidini | 16/08/2018 | Ernage, Belgium | 20.4 | 20.5 | 26.6 | 26.5 | - |
| 199 | ID 576 | Ants | *Heracleum sphondylium* | *Lasius niger* | / | Formicinae | Lasiini | 16/08/2018 | Ernage, Belgium | 20.4 | 20.5 | 26.6 | 26.5 | - |
| 199 | ID 577 | Stems | *Heracleum sphondylium* | / | / | / | / | 16/08/2018 | Ernage, Belgium | 20.4 | 20.5 | 26.6 | 26.5 | - |
| 200 | ID 579 | Aphids | *Heracleum sphondylium* | *Aphis fabae* | polyphagous | Aphidinae | Aphidini | 16/08/2018 | Chastre, Belgium | 20.4 | 20.5 | 26.6 | 26.5 | - |
| 200 | ID 580 | Ants | *Heracleum sphondylium* | *Lasius niger* | / | Formicinae | Lasiini | 16/08/2018 | Chastre, Belgium | 20.4 | 20.5 | 26.6 | 26.5 | - |
| 200 | ID 581 | Stems | *Heracleum sphondylium* | / | / | / | / | 16/08/2018 | Chastre, Belgium | 20.4 | 20.5 | 26.6 | 26.5 | - |
| 201 | ID 582 * | Aphids | *Rosa rugosa* | *Macrosiphum mordvilkoi* | specialized | Aphidinae | Macrosiphini | 16/08/2018 | Chastre, Belgium | 20.4 | 20.5 | 26.6 | 26.5 | + |
| 201 | ID 583 | Stems | *Rosa rugosa* | / | / | / | / | 16/08/2018 | Chastre, Belgium | 20.4 | 20.5 | 26.6 | 26.5 | - |
| 202 | ID 584 | Aphids | *Centranthus ruber* | *Aphis fabae* | polyphagous | Aphidinae | Aphidini | 16/08/2018 | Chastre, Belgium | 20.4 | 20.5 | 26.6 | 26.5 | - |
| 202 | ID 585 | Ants | *Centranthus ruber* | *Lasius niger* | / | Formicinae | Lasiini | 16/08/2018 | Chastre, Belgium | 20.4 | 20.5 | 26.6 | 26.5 | - |
| 202 | ID 586 | Stems | *Centranthus ruber* | / | / | / | / | 16/08/2018 | Chastre, Belgium | 20.4 | 20.5 | 26.6 | 26.5 | - |
| 203 | ID 587 | Aphids | *Sonchus asper* | *Uroleucon sonchi* | restricted | Aphidinae | Macrosiphini | 16/08/2018 | Chastre, Belgium | 20.4 | 20.5 | 26.6 | 26.5 | - |
| 203 | ID 588 | Stems | *Sonchus asper* | / | / | / | / | 16/08/2018 | Chastre, Belgium | 20.4 | 20.5 | 26.6 | 26.5 | - |
| 204 | ID 589 | Aphids | *Sonchus asper* | *Aphis fabae* | polyphagous | Aphidinae | Aphidini | 16/08/2018 | Chastre, Belgium | 20.4 | 20.5 | 26.6 | 26.5 | - |
| 204 | ID 590 | Stems | *Sonchus asper* | / | / | / | / | 16/08/2018 | Chastre, Belgium | 20.4 | 20.5 | 26.6 | 26.5 | - |
| 204 | ID 591 | Ants | *Sonchus asper* | *Lasius niger* | / | Formicinae | Lasiini | 16/08/2018 | Chastre, Belgium | 20.4 | 20.5 | 26.6 | 26.5 | - |
| 204 | ID 592 | Aphids | *Sonchus asper* | *Uroleucon sonchi* | restricted | Aphidinae | Macrosiphini | 16/08/2018 | Chastre, Belgium | 20.4 | 20.5 | 26.6 | 26.5 | - |
| 205 | ID 593 | Aphids | *Sonchus asper* | *Uroleucon sonchi* | restricted | Aphidinae | Macrosiphini | 16/08/2018 | Chastre, Belgium | 20.4 | 20.5 | 26.6 | 26.5 | - |
| 205 | ID 594 | Aphids | *Sonchus asper* | *Hyperomyzus lactucae* | host alternating | Aphidinae | Macrosiphini | 16/08/2018 | Chastre, Belgium | 20.4 | 20.5 | 26.6 | 26.5 | - |
| 205 | ID 595 | Stems | *Sonchus asper* | / | / | / | / | 16/08/2018 | Chastre, Belgium | 20.4 | 20.5 | 26.6 | 26.5 | - |
| 206 | ID 596 | Aphids | *Sonchus asper* | *Uroleucon sonchi* | restricted | Aphidinae | Macrosiphini | 16/08/2018 | Chastre, Belgium | 20.4 | 20.5 | 26.6 | 26.5 | - |
| 206 | ID 597 | Stems | *Sonchus asper* | / | / | / | / | 16/08/2018 | Chastre, Belgium | 20.4 | 20.5 | 26.6 | 26.5 | - |
| 207 | ID 598 | Aphids | *Sonchus asper* | *Uroleucon sonchi* | restricted | Aphidinae | Macrosiphini | 20/08/2018 | Walhain, Belgium | 20.4 | 20.3 | 23 | 22.9 | - |
| 207 | ID 599 | Stems | *Sonchus asper* | / | / | / | / | 20/08/2018 | Walhain, Belgium | 20.4 | 20.3 | 23 | 22.9 | - |
| 207 | ID 600 | Aphids | *Sonchus asper* | *Uroleucon sonchi* | restricted | Aphidinae | Macrosiphini | 20/08/2018 | Walhain, Belgium | 20.4 | 20.3 | 23 | 22.9 | - |
| 208 | ID 601 | Aphids | *Sonchus asper* | *Hyperomyzus lactucae* | host alternating | Aphidinae | Macrosiphini | 20/08/2018 | Walhain, Belgium | 20.4 | 20.3 | 23 | 22.9 | - |
| 208 | ID 602 | Hoverfly Larvae | *Sonchus asper* | *Eupeodes luniger* | / | Syrphinae | Syrphini | 20/08/2018 | Walhain, Belgium | 20.4 | 20.3 | 23 | 22.9 | - |
| 208 | ID 603 | Stems | *Sonchus asper* | / | / | / | / | 20/08/2018 | Walhain, Belgium | 20.4 | 20.3 | 23 | 22.9 | - |
| 209 | ID 604 | Aphids | *Cirsium arvense* | *Aphis fabae* | polyphagous | Aphidinae | Aphidini | 20/08/2018 | Walhain, Belgium | 20.4 | 20.3 | 23 | 22.9 | - |
| 209 | ID 605 * | Aphids | *Cirsium arvense* | *Capitophorus elaeagni* | host alternating | Aphidinae | Macrosiphini | 20/08/2018 | Walhain, Belgium | 20.4 | 20.3 | 23 | 22.9 | + |
| 209 | ID 606 | Bugs | *Cirsium arvense* | *Orius niger* | / | Anthocorinae | Oriini | 20/08/2018 | Walhain, Belgium | 20.4 | 20.3 | 23 | 22.9 | - |
| 209 | ID 607 | Hoverfly Larvae | *Cirsium arvense* | *Scaeva pyrastri* | / | Syrphinae | Syrphini | 20/08/2018 | Walhain, Belgium | 20.4 | 20.3 | 23 | 22.9 | + |
| 209 | ID 608 | Ants | *Cirsium arvense* | *Lasius niger* | / | Formicinae | Lasiini | 20/08/2018 | Walhain, Belgium | 20.4 | 20.3 | 23 | 22.9 | + |
| 209 | ID 609 | Stems | *Cirsium arvense* | / | / | / | / | 20/08/2018 | Walhain, Belgium | 20.4 | 20.3 | 23 | 22.9 | - |
| 210 | ID 611 | Aphids | *Verbena officinalis* | *Aphis frangulae* | polyphagous | Aphidinae | Aphidini | 20/08/2018 | Walhain, Belgium | 20.4 | 20.3 | 23 | 22.9 | - |
| 210 | ID 612 | Ants | *Verbena officinalis* | *Lasius niger* | / | Formicinae | Lasiini | 20/08/2018 | Walhain, Belgium | 20.4 | 20.3 | 23 | 22.9 | - |
| 210 | ID 613 | Hoverfly Larvae | *Verbena officinalis* | *Scaeva pyrastri* | / | Syrphinae | Syrphini | 20/08/2018 | Walhain, Belgium | 20.4 | 20.3 | 23 | 22.9 | - |
| 210 | ID 614 | Stems | *Verbena officinalis* | / | / | / | / | 20/08/2018 | Walhain, Belgium | 20.4 | 20.3 | 23 | 22.9 | - |
| 211 | ID 615 | Aphids | *Sonchus asper* | *Hyperomyzus lactucae* | host alternating | Aphidinae | Macrosiphini | 20/08/2018 | Walhain, Belgium | 20.4 | 20.3 | 23 | 22.9 | - |
| 211 | ID 616 | Aphids | *Sonchus asper* | *Uroleucon sonchi* | restricted | Aphidinae | Macrosiphini | 20/08/2018 | Walhain, Belgium | 20.4 | 20.3 | 23 | 22.9 | - |
| 211 | ID 617 | Aphids | *Sonchus asper* | *Uroleucon sonchi* | restricted | Aphidinae | Macrosiphini | 20/08/2018 | Walhain, Belgium | 20.4 | 20.3 | 23 | 22.9 | - |
| 211 | ID 618 | Hoverfly Larvae | *Sonchus asper* | *Episyrphus balteatus* | / | Syrphinae | Syrphini | 20/08/2018 | Walhain, Belgium | 20.4 | 20.3 | 23 | 22.9 | - |
| 211 | ID 619 | Stems | *Sonchus asper* | / | / | / | / | 20/08/2018 | Walhain, Belgium | 20.4 | 20.3 | 23 | 22.9 | - |
| 212 | ID 621 | Aphids | *Sonchus asper* | *Uroleucon sonchi* | restricted | Aphidinae | Macrosiphini | 20/08/2018 | Walhain, Belgium | 20.4 | 20.3 | 23 | 22.9 | - |
| 212 | ID 622 | Aphids | *Sonchus asper* | *Uroleucon sonchi* | restricted | Aphidinae | Macrosiphini | 20/08/2018 | Walhain, Belgium | 20.4 | 20.3 | 23 | 22.9 | - |
| 212 | ID 623 | Ants | *Sonchus asper* | *Lasius niger* | / | Formicinae | Lasiini | 20/08/2018 | Walhain, Belgium | 20.4 | 20.3 | 23 | 22.9 | - |
| 212 | ID 624 | Parasitoids | *Sonchus asper* | *Lysiphlebus fabarum* | / | Aphidiinae | Aphidiini | 20/08/2018 | Walhain, Belgium | 20.4 | 20.3 | 23 | 22.9 | - |
| 212 | ID 625 | Hoverfly Larvae | *Sonchus asper* | *Eupeodes luniger* | / | Syrphinae | Syrphini | 20/08/2018 | Walhain, Belgium | 20.4 | 20.3 | 23 | 22.9 | - |
| 212 | ID 626 | Stems | *Sonchus asper* | / | / | / | / | 20/08/2018 | Walhain, Belgium | 20.4 | 20.3 | 23 | 22.9 | - |
| 213 | ID 628 | Aphids | *Sonchus arvensis* | *Uroleucon sonchi* | restricted | Aphidinae | Macrosiphini | 20/08/2018 | Walhain, Belgium | 20.4 | 20.3 | 23 | 22.9 | - |
| 213 | ID 629 | Aphids | *Sonchus arvensis* | *Hyperomyzus lactucae* | host alternating | Aphidinae | Macrosiphini | 20/08/2018 | Walhain, Belgium | 20.4 | 20.3 | 23 | 22.9 | - |
| 213 | ID 630 | Parasitoids | *Sonchus arvensis* | *Tetrastichinae sp.* | / | Tetrastichinae | NA | 20/08/2018 | Walhain, Belgium | 20.4 | 20.3 | 23 | 22.9 | - |
| 213 | ID 631 | Hoverfly Larvae | *Sonchus arvensis* | *Paragus cooverti* | / | Syrphinae | Paragini | 20/08/2018 | Walhain, Belgium | 20.4 | 20.3 | 23 | 22.9 | - |
| 213 | ID 632 | Stems | *Sonchus arvensis* | / | / | / | / | 20/08/2018 | Walhain, Belgium | 20.4 | 20.3 | 23 | 22.9 | - |
| 214 | ID 633 | Bugs | *Heracleum sphondylium* | *Orthops campestris* | / | Mirinae | Mirini | 20/08/2018 | Walhain, Belgium | 20.4 | 20.3 | 23 | 22.9 | - |
| 214 | ID 634 | Bugs | *Heracleum sphondylium* | *Graphosoma lineatum* | / | Podopinae | Graphosomatini | 20/08/2018 | Walhain, Belgium | 20.4 | 20.3 | 23 | 22.9 | - |
| 214 | ID 635 | Stems | *Heracleum sphondylium* | / | / | / | / | 20/08/2018 | Walhain, Belgium | 20.4 | 20.3 | 23 | 22.9 | - |
| 215 | ID 637 | Aphids | *Heracleum sphondylium* | *Aphis fabae* | polyphagous | Aphidinae | Aphidini | 20/08/2018 | Walhain, Belgium | 20.4 | 20.3 | 23 | 22.9 | - |
| 215 | ID 638 | Ants | *Heracleum sphondylium* | *Lasius niger* | / | Formicinae | Lasiini | 20/08/2018 | Walhain, Belgium | 20.4 | 20.3 | 23 | 22.9 | - |
| 215 | ID 639 | Stems | *Heracleum sphondylium* | / | / | / | / | 20/08/2018 | Walhain, Belgium | 20.4 | 20.3 | 23 | 22.9 | - |
| 216 | ID 641 | Aphids | *Sonchus asper* | *Uroleucon sonchi* | restricted | Aphidinae | Macrosiphini | 21/08/2018 | Orbais, Belgium | 21.5 | 21.4 | 26.6 | 26.3 | - |
| 216 | ID 642 | Aphid Midge Larvae | *Sonchus asper* | *Aphidoletes aphidimyza* | / | Cecidomyiinae | Aphidoletini | 21/08/2018 | Orbais, Belgium | 21.5 | 21.4 | 26.6 | 26.3 | - |
| 216 | ID 643 | Stems | *Sonchus asper* | / | / | / | / | 21/08/2018 | Orbais, Belgium | 21.5 | 21.4 | 26.6 | 26.3 | - |
| 217 | ID 645 | Aphids | *Sonchus asper* | *Uroleucon sonchi* | restricted | Aphidinae | Macrosiphini | 21/08/2018 | Orbais, Belgium | 21.5 | 21.4 | 26.6 | 26.3 | - |
| 217 | ID 646 | Ladybugs | *Sonchus asper* | NA | / | NA | NA | 21/08/2018 | Orbais, Belgium | 21.5 | 21.4 | 26.6 | 26.3 | - |
| 217 | ID 647 | Hoverfly Larvae | *Sonchus asper* | *Eupeodes luniger* | / | Syrphinae | Syrphini | 21/08/2018 | Orbais, Belgium | 21.5 | 21.4 | 26.6 | 26.3 | - |
| 217 | ID 648 | Aphids | *Sonchus asper* | *Hyperomyzus lactucae* | host alternating | Aphidinae | Macrosiphini | 21/08/2018 | Orbais, Belgium | 21.5 | 21.4 | 26.6 | 26.3 | - |
| 217 | ID 649 | Stems | *Sonchus asper* | / | / | / | / | 21/08/2018 | Orbais, Belgium | 21.5 | 21.4 | 26.6 | 26.3 | - |
| 218 | ID 650 | Aphids | *Sonchus asper* | *Uroleucon sonchi* | restricted | Aphidinae | Macrosiphini | 21/08/2018 | Orbais, Belgium | 21.5 | 21.4 | 26.6 | 26.3 | - |
| 218 | ID 651 | Aphids | *Sonchus asper* | *Hyperomyzus lactucae* | host alternating | Aphidinae | Macrosiphini | 21/08/2018 | Orbais, Belgium | 21.5 | 21.4 | 26.6 | 26.3 | - |
| 218 | ID 652 | Hoverfly Larvae | *Sonchus asper* | *Paragus cooverti* | / | Syrphinae | Paragini | 21/08/2018 | Orbais, Belgium | 21.5 | 21.4 | 26.6 | 26.3 | - |
| 218 | ID 653 | Stems | *Sonchus asper* | / | / | / | / | 21/08/2018 | Orbais, Belgium | 21.5 | 21.4 | 26.6 | 26.3 | - |
| 219 | ID 655 | Aphids | *Hibiscus syriacus* | *Aphis fabae* | polyphagous | Aphidinae | Aphidini | 21/08/2018 | Orbais, Belgium | 21.5 | 21.4 | 26.6 | 26.3 | - |
| 219 | ID 656 | Ants | *Hibiscus syriacus* | *Lasius niger* | / | Formicinae | Lasiini | 21/08/2018 | Orbais, Belgium | 21.5 | 21.4 | 26.6 | 26.3 | - |
| 219 | ID 657 | Stems | *Hibiscus syriacus* | / | / | / | / | 21/08/2018 | Orbais, Belgium | 21.5 | 21.4 | 26.6 | 26.3 | - |
| 220 | ID 659 | Aphids | *Epilobium hirsutum* | *Aphis mamonthovae* | NA | Aphidinae | Aphidini | 21/08/2018 | Orbais, Belgium | 21.5 | 21.4 | 26.6 | 26.3 | - |
| 220 | ID 660 | Parasitoids | *Epilobium hirsutum* | *Binodoxys communis* | / | Aphidiinae | Aphidiini | 21/08/2018 | Orbais, Belgium | 21.5 | 21.4 | 26.6 | 26.3 | - |
| 220 | ID 661 | Stems | *Epilobium hirsutum* | / | / | / | / | 21/08/2018 | Orbais, Belgium | 21.5 | 21.4 | 26.6 | 26.3 | - |
| 221 | ID 663 | Aphids | *Verbena officinalis* | *Aphis mamonthovae* | NA | Aphidinae | Aphidini | 21/08/2018 | Malèves, Belgium | 21.5 | 21.4 | 26.6 | 26.3 | - |
| 221 | ID 664 | Ants | *Verbena officinalis* | *Lasius niger* | / | Formicinae | Lasiini | 21/08/2018 | Malèves, Belgium | 21.5 | 21.4 | 26.6 | 26.3 | - |
| 221 | ID 665 | Stems | *Verbena officinalis* | / | / | / | / | 21/08/2018 | Malèves, Belgium | 21.5 | 21.4 | 26.6 | 26.3 | - |
| 222 | ID 666 | Aphids | *Heracleum sphondylium* | *Aphis fabae* | polyphagous | Aphidinae | Aphidini | 21/08/2018 | Malèves, Belgium | 21.5 | 21.4 | 26.6 | 26.3 | - |
| 222 | ID 667 | Aphids | *Heracleum sphondylium* | *Cavariella pastinacae* | host alternating | Aphidinae | Macrosiphini | 21/08/2018 | Malèves, Belgium | 21.5 | 21.4 | 26.6 | 26.3 | - |
| 222 | ID 668 | Stems | *Heracleum sphondylium* | / | / | / | / | 21/08/2018 | Malèves, Belgium | 21.5 | 21.4 | 26.6 | 26.3 | - |
| 223 | ID 670 | Aphids | *Heracleum sphondylium* | *Cavariella theobaldi* | host alternating | Aphidinae | Macrosiphini | 21/08/2018 | Malèves, Belgium | 21.5 | 21.4 | 26.6 | 26.3 | + |
| 223 | ID 671 | Aphids | *Heracleum sphondylium* | *Aphis fabae* | polyphagous | Aphidinae | Aphidini | 21/08/2018 | Malèves, Belgium | 21.5 | 21.4 | 26.6 | 26.3 | + |
| 223 | ID 672 | Aphids | *Heracleum sphondylium* | *Hyadaphis foeniculi* | host alternating | Aphidinae | Macrosiphini | 21/08/2018 | Malèves, Belgium | 21.5 | 21.4 | 26.6 | 26.3 | - |
| 223 | ID 673 | Ants | *Heracleum sphondylium* | NA | / | NA | NA | 21/08/2018 | Malèves, Belgium | 21.5 | 21.4 | 26.6 | 26.3 | - |
| 223 | ID 674 | Hoverfly Larvae | *Heracleum sphondylium* | *Syrphus ribesii* | / | Syrphinae | Syrphini | 21/08/2018 | Malèves, Belgium | 21.5 | 21.4 | 26.6 | 26.3 | - |
| 223 | ID 675 | Stems | *Heracleum sphondylium* | / | / | / | / | 21/08/2018 | Malèves, Belgium | 21.5 | 21.4 | 26.6 | 26.3 | - |
| 224 | ID 677 | Aphids | *Sonchus asper* | *Aphis fabae* | polyphagous | Aphidinae | Aphidini | 21/08/2018 | Malèves, Belgium | 21.5 | 21.4 | 26.6 | 26.3 | - |
| 224 | ID 678 | Ants | *Sonchus asper* | *Lasius niger* | / | Formicinae | Lasiini | 21/08/2018 | Malèves, Belgium | 21.5 | 21.4 | 26.6 | 26.3 | - |
| 224 | ID 679 | Stems | *Sonchus asper* | / | / | / | / | 21/08/2018 | Malèves, Belgium | 21.5 | 21.4 | 26.6 | 26.3 | - |
| 224 | ID 681 | Parasitoids | *Sonchus asper* | *Pachyneuron aphidis* | / | Pteromalinae | Pachyneurini | 21/08/2018 | Malèves, Belgium | 21.5 | 21.4 | 26.6 | 26.3 | - |
| 225 | ID 682 | Aphids | *Spiraea salicifolia* | *Aphis spiraecola* | polyphagous | Aphidinae | Aphidini | 21/08/2018 | Opprebais, Belgium | 21.5 | 21.4 | 26.6 | 26.3 | - |
| 225 | ID 683 | Ants | *Spiraea sp.* | NA | / | NA | NA | 21/08/2018 | Opprebais, Belgium | 21.5 | 21.4 | 26.6 | 26.3 | - |
| 225 | ID 684 | Stems | *Spiraea sp.* | / | / | / | / | 21/08/2018 | Opprebais, Belgium | 21.5 | 21.4 | 26.6 | 26.3 | - |
| 226 | ID 685 | Aphids | *Sonchus asper* | *Hyperomyzus lactucae* | host alternating | Aphidinae | Macrosiphini | 21/08/2018 | Opprebais, Belgium | 21.5 | 21.4 | 26.6 | 26.3 | - |
| 226 | ID 686 | Parasitoids | *Sonchus asper* | *Tetrastichinae sp.* | / | Tetrastichinae | NA | 21/08/2018 | Opprebais, Belgium | 21.5 | 21.4 | 26.6 | 26.3 | - |
| 226 | ID 687 | Stems | *Sonchus asper* | / | / | / | / | 21/08/2018 | Opprebais, Belgium | 21.5 | 21.4 | 26.6 | 26.3 | - |
| 227 | ID 688 | Aphids | *Tanacetum vulgare* | NA | NA | NA | NA | 22/08/2018 | Longueville, Belgium | 20.4 | 20.2 | 24.5 | 24.9 | - |
| 227 | ID 689 | Ants | *Tanacetum vulgare* | NA | / | NA | NA | 22/08/2018 | Longueville, Belgium | 20.4 | 20.2 | 24.5 | 24.9 | - |
| 227 | ID 690 | Stems | *Tanacetum vulgare* | / | / | / | / | 22/08/2018 | Longueville, Belgium | 20.4 | 20.2 | 24.5 | 24.9 | - |
| 228 | ID 692 | Aphids | *Oenothera glazioviana* | *Aphis oenotherae* | restricted | Aphidinae | Aphidini | 22/08/2018 | Gottechain, Belgium | 20.4 | 20.2 | 24.5 | 24.9 | - |
| 228 | ID 693 | Ants | *Oenothera glazioviana* | NA | / | NA | NA | 22/08/2018 | Gottechain, Belgium | 20.4 | 20.2 | 24.5 | 24.9 | - |
| 228 | ID 694 | Stems | *Oenothera glazioviana* | / | / | / | / | 22/08/2018 | Gottechain, Belgium | 20.4 | 20.2 | 24.5 | 24.9 | - |
| 229 | ID 696 | Aphids | *Sonchus asper* | *Uroleucon sonchi* | restricted | Aphidinae | Macrosiphini | 22/08/2018 | Courtil-Noirmont, Belgium | 20.4 | 20.2 | 24.5 | 24.9 | - |
| 229 | ID 697 | Stems | *Sonchus asper* | / | / | / | / | 22/08/2018 | Courtil-Noirmont, Belgium | 20.4 | 20.2 | 24.5 | 24.9 | - |
| 230 | ID 698 | Aphids | *Cirsium arvense* | *Uroleucon sonchi* | restricted | Aphidinae | Macrosiphini | 22/08/2018 | Courtil-Noirmont, Belgium | 20.4 | 20.2 | 24.5 | 24.9 | - |
| 230 | ID 699 | Aphid Midge Larvae | *Cirsium arvense* | *Aphidoletes aphidimyza* | / | Cecidomyiinae | Aphidoletini | 22/08/2018 | Courtil-Noirmont, Belgium | 20.4 | 20.2 | 24.5 | 24.9 | - |
| 230 | ID 700 | Hoverfly Larvae | *Cirsium arvense* | *Paragus cooverti* | / | Syrphinae | Paragini | 22/08/2018 | Courtil-Noirmont, Belgium | 20.4 | 20.2 | 24.5 | 24.9 | - |
| 230 | ID 701 | Hoverfly Larvae | *Cirsium arvense* | *Scaeva pyrastri* | / | Syrphinae | Syrphini | 22/08/2018 | Courtil-Noirmont, Belgium | 20.4 | 20.2 | 24.5 | 24.9 | - |
| 230 | ID 702 | Stems | Cirsium arvense | / | / | / | / | 22/08/2018 | Courtil-Noirmont, Belgium | 20.4 | 20.2 | 24.5 | 24.9 | - |
| 231 | ID 703 | Aphids | *Cirsium arvense* | *Aphis fabae* | polyphagous | Aphidinae | Aphidini | 22/08/2018 | Courtil-Noirmont, Belgium | 20.4 | 20.2 | 24.5 | 24.9 | - |
| 231 | ID 704 | Ants | *Cirsium arvense* | *Lasius niger* | / | Formicinae | Lasiini | 22/08/2018 | Courtil-Noirmont, Belgium | 20.4 | 20.2 | 24.5 | 24.9 | - |
| 231 | ID 705 | Stems | *Cirsium arvense* | / | / | / | / | 22/08/2018 | Courtil-Noirmont, Belgium | 20.4 | 20.2 | 24.5 | 24.9 | - |

**Table S2.** Genes and primers for screening and sequencing.

| **Gene** | **Hypothtical product** | **Primer** | **Sequence (5'-3')** | **Tm** | **Fragment size (bp)** | **References** |
| --- | --- | --- | --- | --- | --- | --- |
| Insect species |  |  |  |  |  |  |
| COI | Cytochrome oxidase subunit I | LepF | ATTCAACCAATCATAAAGATATTGG | 45°C - 51°C |  | (Hajibabaei et al., 2006) |
|  |  | LepR | TAAACTTCTGGATGTCCAAAAAATCA |  | 658 |  |
| *Serratia symbiotica* |  |  |  |  |  |  |
| 16S rRNA | Ribosomal RNA | 16SA1F | AGAGTTTGATCMTGGCTCAG | 55°C |  | (Fukatsu & Nikoh, 1998) |
|  |  | PASScmpR | GCAATGTCTTATTAACACAT |  | 480 | (Fukatsu et al., 2000) |
| accD | Carboxyl transferase, subunit β | accD.S 2F | ACACCCTACTGGATAAAGGC | 60°C |  | (Henry et al., 2003) |
|  |  | accD.S 2R | GATGTTGATGTCGCCCAGC |  | 522 |  |
| gyrB | DNA gyrase, subunit B | gyrB.S 2F | TGACATTCTGGCCAAGCGCC | 64°C |  | (Henry et al., 2003) |
|  |  | gyrB.S 1R | ACTACCGCGATCAGCCCCTC |  | 615 |  |
| murE | UDP-N-acetylmuramoylalanyl-Dglutamate 2,6-diaminopimelate | murES 6F | CTGTTCGCTGGGCATGATGTGG | 55°C |  | (Henry et al., 2003) |
|  |  | murE 6R | GCCCGGTGCGTTAAACACTTCC |  | 774 |  |
| recJ | 5' –> 3' exonuclease | recJ.S 4F | GAAGCAATCGTTAATCCC | 60°C |  | (Henry et al., 2003) |
|  |  | recJ.S 4R | TATCGAGATCGTTAGCCAGC |  | 840 |  |

**Table S3.** Accession numbers of *S. symbiotica* targeted sequences.

**Table S4.** Accession numbers of *S. symbiotica* targeted sequences and genomes.
